# Insights from a survey of mentorship experiences by graduate and postdoctoral researchers

**DOI:** 10.1101/2023.05.05.539640

**Authors:** Sarvenaz Sarabipour, Natalie M Niemi, Steven J Burgess, Christopher T Smith, Alexandre W Bisson Filho, Ahmed Ibrahim, Kelly Clark

## Abstract

Mentorship is vital for early career researchers in training positions, allowing them to navigate the challenges of work and life in research environments. However, the quality of mentorship received by trainees can vary by investigator and by institution. One challenge faced by those hoping to improve trainee mentorship is that the extent to which mentorship is offered to and experienced by research trainees is not well characterized. To address this knowledge gap, we conducted a survey to examine the quality of mentorship received by trainees in research environments, to identify characteristics of positive and negative mentorship, and to highlight best practices to improve trainee mentorship. We received 2,114 responses from researchers at graduate and postdoctoral career stages worldwide. Quantitative analysis showed that at least ∼25-45% of respondents were dissatisfied with some aspects of their mentorship. Qualitative responses revealed that common issues in mentorship include unclear expectations in research and mentoring interactions, lack of guidance, and inadequate support of trainee independence and career goals. Our findings also identified key mentorship elements desired by trainee mentees. Based on trainee suggestions, we describe strategies for individual mentors, departments, and institutions to improve the training experience for graduate and postdoctoral researchers.

## Introduction

Mentorship is regarded as guidance provided by a more experienced and knowledgeable person as a guide and sponsor (i.e., the mentor) to a less experienced individual, the mentee. A recent National Academies report defines academic mentorship as “a professional, working alliance in which individuals work together over time to support the personal and professional growth, development, and success of the relational partners through the provision of career and psychosocial support” (National Academies of Sciences, 2019). Mentorship can also occur in the setting of academic groups (e.g., in laboratories) defined as unilateral support from an advisor (i.e., a research group leader) to mentees (i.e., research group members). In research environments, mentorship for trainees is predominantly expected from their faculty advisor or the principal investigator of the research group with which they are associated. It has been suggested that interactions with their research advisor exerts the highest influence amongst the experiences of research trainees (Barnes and Austin, 2009; Lovitts, 2001; Sverdlik et al., 2018; Zhao et al., 2007). This single mentor model may be complemented by a variety of mentorship formats, such as mentoring networks (e.g., principal investigators of collaborating labs for postdoctoral researchers or thesis committees for graduate students), peer-mentoring, group mentoring, or distance mentoring, which can occur on an informal or formal basis (Bielczyk et al., 2019; Guardia et al., 2021; Lorenzetti et al., 2019; Sarabipour et al., 2021).

Graduate students and postdoctoral fellows constitute the majority of the academic scientific community and contribute significantly to the research enterprise (Boothby et al., 2022). Despite their importance in academic research environments, trainees can experience a number of challenges during their training, research endeavors, and career. These include poor mental health and attrition (Maher et al., 2020; Sverdlik et al., 2018), poor work-life balance, internal and external stress (Evans et al., 2018), and impostor syndrome. A contributor to these challenges may be poor mentorship (Jeste et al., 2009). Multiple studies have shown that mentorship is vital for the success and well-being of trainees in research environments (Eby et al., 2008; Ragins et al., 2000; Sosik and Godshalk, 2000). However, merely being in a mentoring relationship does not always lead to positive outcomes for trainees as not all mentorship interactions are beneficial to mentees (Eby et al., 2000; Ragins et al., 2000). Constructive mentorship can improve trainee research performance and self-efficacy (Tenenbaum et al., 2001), accelerate their time to degree completion (Lunsford, 2012), promote satisfaction with their academic program (Lovitts, 2001; Sverdlik et al., 2018; Zhao et al., 2007), enhance their career progression (Allen et al., 2004), and improve prospects of securing academic positions (Fernandes et al., 2020; Liénard et al., 2018).

In contrast, lack of mentorship can isolate early career researchers who are facing the challenges of academic research, limiting their career development. Several recent studies identified factors influencing the well-being of graduate students and postdoctoral researchers, aiming to raise awareness on these issues (Evans et al., 2018; Jeste et al., 2009; Loissel, 2020). Poor mentorship can be detrimental to researchers’ career progression and well-being (Christian et al., 2021; Fernandes et al., 2020; Grinstein and Treister, 2018; McConnell et al., 2018), and lack of effective mentorship may have particularly negative effects on select trainee demographics, including those that face additional challenges such as being from an underrepresented background, identifying as a first-generation scientist, or moving to a foreign country for their training. For instance, international or immigrant researchers make up around 25% of all science and technology workers and around 50% of the doctoral-level science workforce in the United States (“Foreign-born STEM Workers in the United States,” 2022), and temporary visa holders exceed citizens and permanent residents combined at the level of postdoctoral scholars in the United States (“Survey of Graduate Students and Postdoctorates in Science and Engineering Fall 2017: Citizenship of graduate students and postdoctoral appointees in science, engineering, and health: 1980–2017,” n.d.). Furthermore, more than 30% of the total scientific workforce in Canada, Sweden, Switzerland, UK and Australia is foreign-born (Franzoni et al., 2012). This population can experience additional challenges during their research training career due to their foreign status (Fleming, 2022; Johnson et al., 2018; Moss and Mahmoudi, 2021), as can trainees who identify as women and/or underrepresented (Batut et al., 2021; Campbell, and Rodríguez, 2018; Johnson et al., 2018; National Academies of Sciences, 2019; Thomas et al., 2007).

Despite its recognition as vital for the development of trainees in STEM fields, mentorship as experienced by trainees in research environments remains understudied compared to other work environments (National Academies of Sciences, 2019). It thus remains difficult to quantify the quality and frequency of mentorship in individual laboratories, departments, and universities. Academic mentors and leaders would benefit from understanding trainee mentees’ expectations and experiences of mentorship interactions. Furthermore, there is a growing interest in improving mentorship, as it is increasingly recognized as a key factor in creating systemic change by improving researcher training and career outcomes (National Academies of Sciences, 2019). In this regard, multiple national institutions in the United States have created resources on recommending best practices for mentoring relationships (“National Institute of Neurological Disorders and Stroke,” 2022; Pfund et al., 2015; Reynolds, 2019; Sorkness et al., 2017). However, it remains unclear the extent to which these practices are employed within mentoring relationships, particularly with respect to the trainee’s experiences and perception in this space. Identifying desirable and undesirable mentorship traits as experienced by trainees would allow substantial improvements in mentorship relationships. Unfortunately, there is a paucity of knowledge on trainee mentorship structures in research environments (National Academies of Sciences, 2019), and this requires attention given the dependence of trainees on their advisors for support and the significant role that advisors play during their education and research. Notably, mentor training is neither typically required in faculty hiring nor tenure and promotion practices (Fernandes et al., 2019; Wright and Vanderford, 2017) and mentoring programs for trainee mentees and mentor training programs for faculty mentors are scarce. Finally, there is a paucity of data and guidelines for mentor-mentee expectations and interactions at departmental and institutional levels.

To address these gaps in knowledge, we conducted a survey over 2,000 trainees to understand the general nature of their mentoring relationships and how these interactions relate to their satisfaction with the mentoring guidance they received. In our survey, a mentee is defined as a student or researcher in a training position. We asked these researchers to provide anonymous feedback on the mentorship received from their primary mentor. The results show that the quality of mentorship varies greatly across labs and between mentees. Additional insight was gained on issues at the level of laboratories, departments, institutions, and scientific disciplines that impact mentorship interactions and professional development. Based on these findings, we recommend strategies for mentors to employ to strengthen their mentoring skills and improve experiences for their mentees. In addition, we suggest that departments, funders, and institutions provide oversight and guidelines to foster effective mentoring interactions in research environments.

## Results

We received responses from 2,114 trainees (i.e., graduate students and postdoctoral researchers) from 66 countries (Figure 1). By continent 52% of our responses originated from North America, 29% from Europe, 8% from Asia, 6% from Latin America, 4% from Oceania, and 2% from Africa. The majority of our responses were from researchers working in the United States (48%) followed by the United Kingdom (7%), Germany (4%), Spain (4%), Canada (4%), France (3%), Australia (3%), Argentina (3%), India (2%), Switzerland (2%) and under 2% each from over fifty other countries. The majority (94%) of respondents performed their research at an academic institution (Figure 1B). Over half (52%) of respondents performed basic research in life and biomedical sciences, 17% in physical and mathematical sciences, computer science or engineering, 11% in social and behavioral sciences and humanities disciplines, and under ten percent in all other fields (Figure 1C). Most respondents held a bachelor’s degree (26%), or a master’s degree (31%), while 37% held a doctoral degree, with a minority holding dual degrees, a professional degree only, or a high school diploma (Figure 1D). Survey respondents were distributed across self-identified genders, with 38% identifying as male, 61% as female, and 1% identified as non-binary or preferred to not disclose this information (Figure 1E).

**Figure 1.**
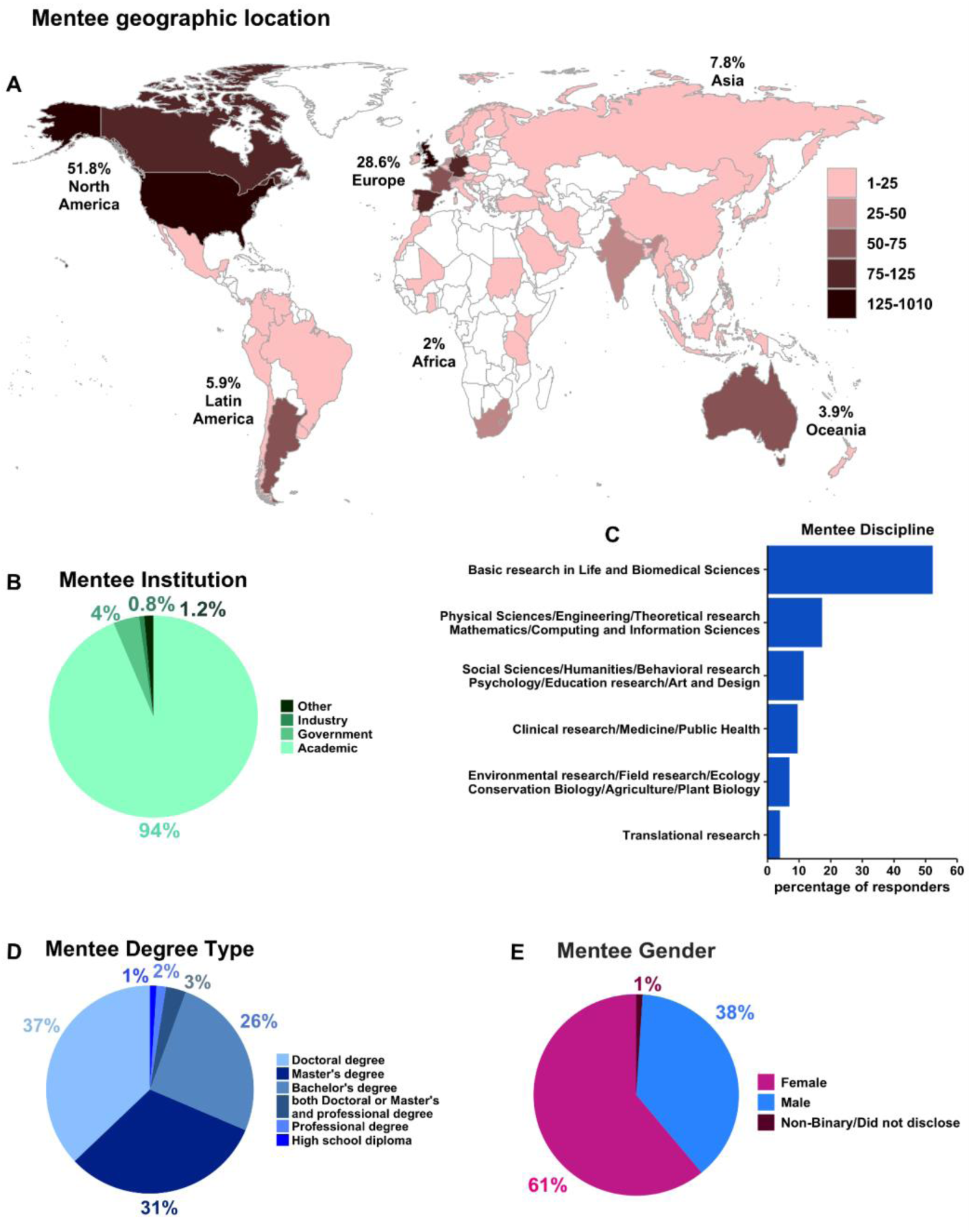
Demographics of trainee mentees. Distribution of the 2,114 researchers who responded to the mentoring survey by (A) Country of research (where institutional research lab affiliation was based Table S1), (B) The type of research institution in which the trainee mentee performed research. “Other” institutions were non-profit organizations and the hospital sector (Table S2), (C) Mentee scientific discipline spanned basic research in life and biomedical sciences (52%), physical sciences, theoretical research, engineering, mathematics (17%), social sciences, humanities, psychology, education research, art and design (11%), clinical research, medicine, public health (9%), environmental research, field research, ecology, agriculture, conservation biology, plant biology (7%), and translational research (4%) (Table S3), (D) Highest advanced degree mentees held: Doctoral degree (PhD), professional degree (e.g., MD, DDS, RD, PT, PharmD, etc.), both PhD and professional degree (MD/PhD, MD/MPH, PharmD/MS, etc.), Master’s degree (science, arts, humanities), Bachelor’s degree (science, arts, humanities) or high school diploma (Table S4), (E) Mentee gender (Table S5). All responses were self-identified.

### Characteristics of mentorship interactions

To examine how mentorship is offered to research trainees, we queried whether mentorship was taking place and, if so, how mentoring relationships functioned among our respondents. Most respondents (57%) were graduate students while 40% were postdoctoral researchers, with a small number of post-graduate professionals, fellows, and undergraduate students (3%) (Figure 2A). These trainees reported variable desired mentorship levels, with the majority of trainees preferring moderate management with some guidance or input (Figure 2B). Most mentees worked in an experimental setting (49%), while 23% worked in theoretical or computational groups, 23% of trainees report working in a lab with both components, and 6% performed research in other settings (Figure 2C). Overall, most respondents (71%) found their interactions with their mentor constructive (Figure 2D). The majority used face-to-face meetings and email to communicate with their mentor (41%) (Figure 2E). Most mentees met with their mentor weekly, every two weeks, or scheduled their meetings when necessary (Figure 2F). The majority of mentees chose their mentors voluntarily (93%) while a minority had a mentor assigned by their institution (7%) (Figure 3A). The majority of respondents (>99%) indicated that they were receiving some mentorship, a small fraction (0.4%) of respondents reported not having a mentor. Over half (53%) reported one key mentor, while 46% had two or more mentors (Figure 3B). The majority of survey respondents (70%) were 25-35 years of age (Figure 3C). Over 85% of respondents had worked with their mentor for at least one year (Figure 3D). Most respondents (70%) regarded their mentoring match experience as good-to-excellent, although a significant portion (30%) characterize the quality of mentorship match as poor-to-fair (Figure 3E).

**Figure 2.**
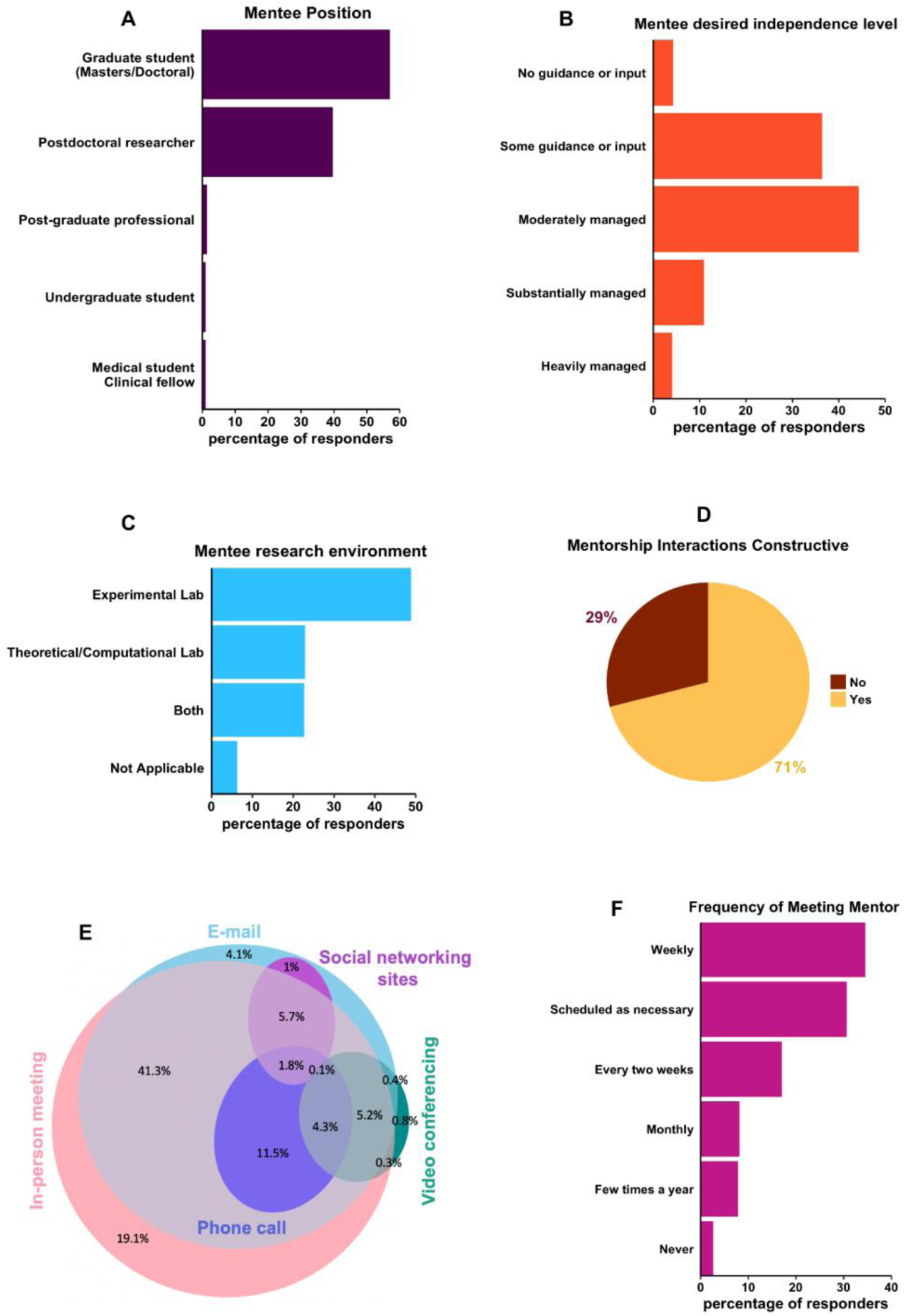
Characterization of mentorship environment and mentor accessibility. (A) Mentee academic positions spanned graduate students (in Master’s or Doctoral training) (57%) postdoctoral researchers (40%), post-graduate professionals (1%), undergraduate students (1%) and medical students or clinical fellows (1%) (Table S6), (B) Mentee desired level of independence (Table S7), (C) Mentee research environment (i.e., wet lab/bench research, theoretical/experimental research or both formats) (Table S8), (D) Quality of mentee-mentor interactions (Table S9), (E) Mentee responses to the question on format of communication with mentor. 24% of respondents only used one of the in-person (face-to-face) meeting, email, phone call, video/web conferencing, chat/asynchronous interactions via social networking sites (such as Slack) format and 76% used a combination of these formats (Table S10), (F) Mentee-mentor meeting frequency was scheduled as necessary (31%), weekly (34%), every two weeks (17%), monthly (8%), yearly (8%), and never (3%) (Table S11). All responses were self-identified.

**Figure 3.**
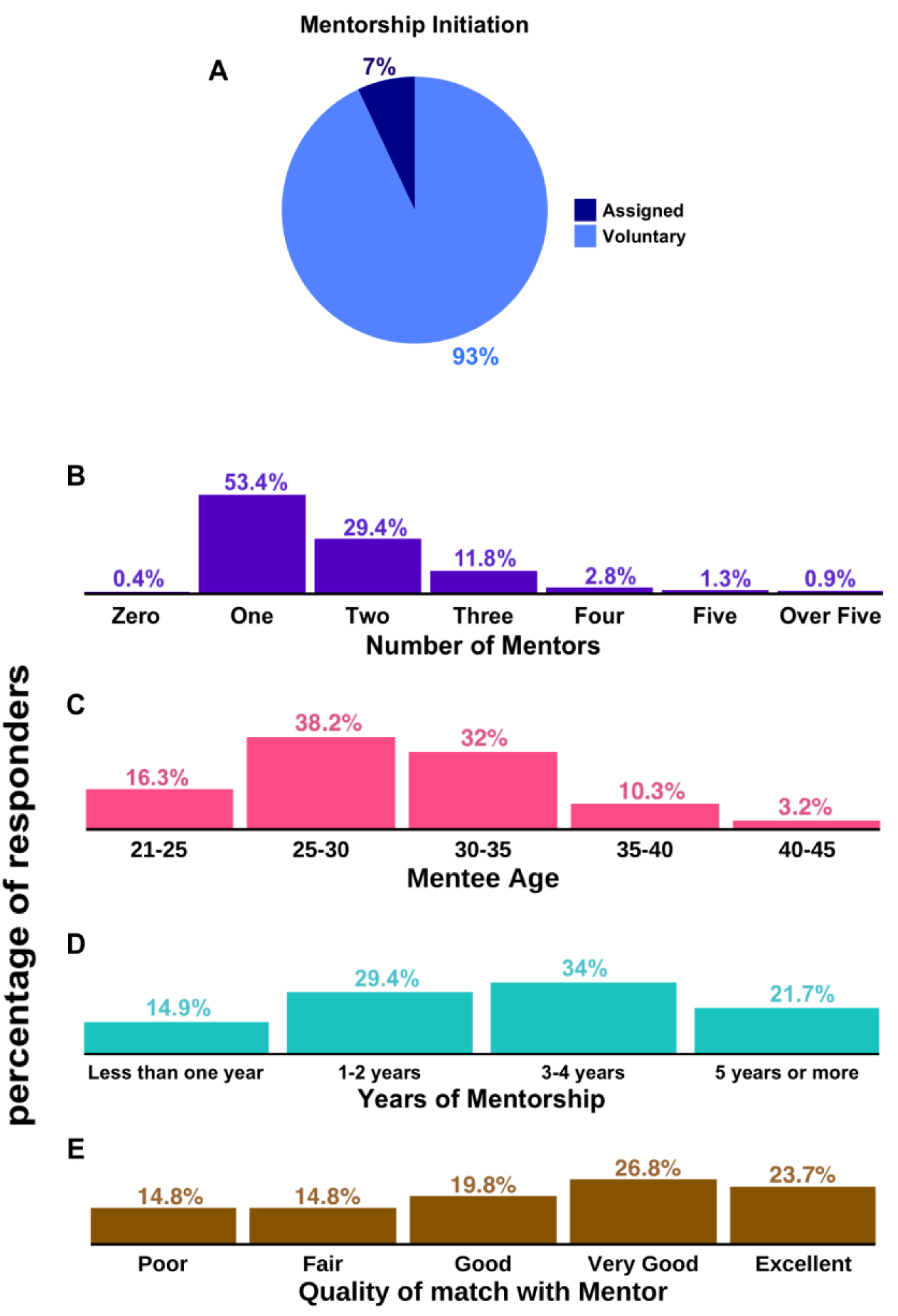
Characteristics of mentees and mentorship interactions. (A) Mentorship initiation mode (Table S12), (B) Number of mentors (Table S13), (C) Mentee age distribution (Table S14), (D) Years that mentorship was received or years of working with the mentor(s) (Table S15), (E) Quality of mentee-mentor interactions (Table S16). All responses were self-identified.

For our geographic analyses, we stratified our analyses to North America; Europe; and Latin America, Africa, Oceania and Asia combined; this classification resulted in reasonably sized subdivisions of our data. Looking across continents, trainee mentees in Europe expressed higher dissatisfaction with their mentorship interactions compared to mentees researchers in other continents (Figure 4A-D). Cross-analysis with respondent citizenship status in country of research shows that while majority of respondents working in North America, Latin America/Africa/Asia/Oceania were citizens in their country of research (67% citizens for North America and 76% citizens for researchers in Latin America/Africa/Asia/Oceania), the majority of respondents from Europe were not citizens in their country of research (59%). Thus, the connection between respondent citizenship status and continent of research location may contribute to the higher dissatisfaction rates of European respondents, as international trainee status appears to serve as a negative factor in mentoring experiences (Table S35-S36).

**Figure 4.**
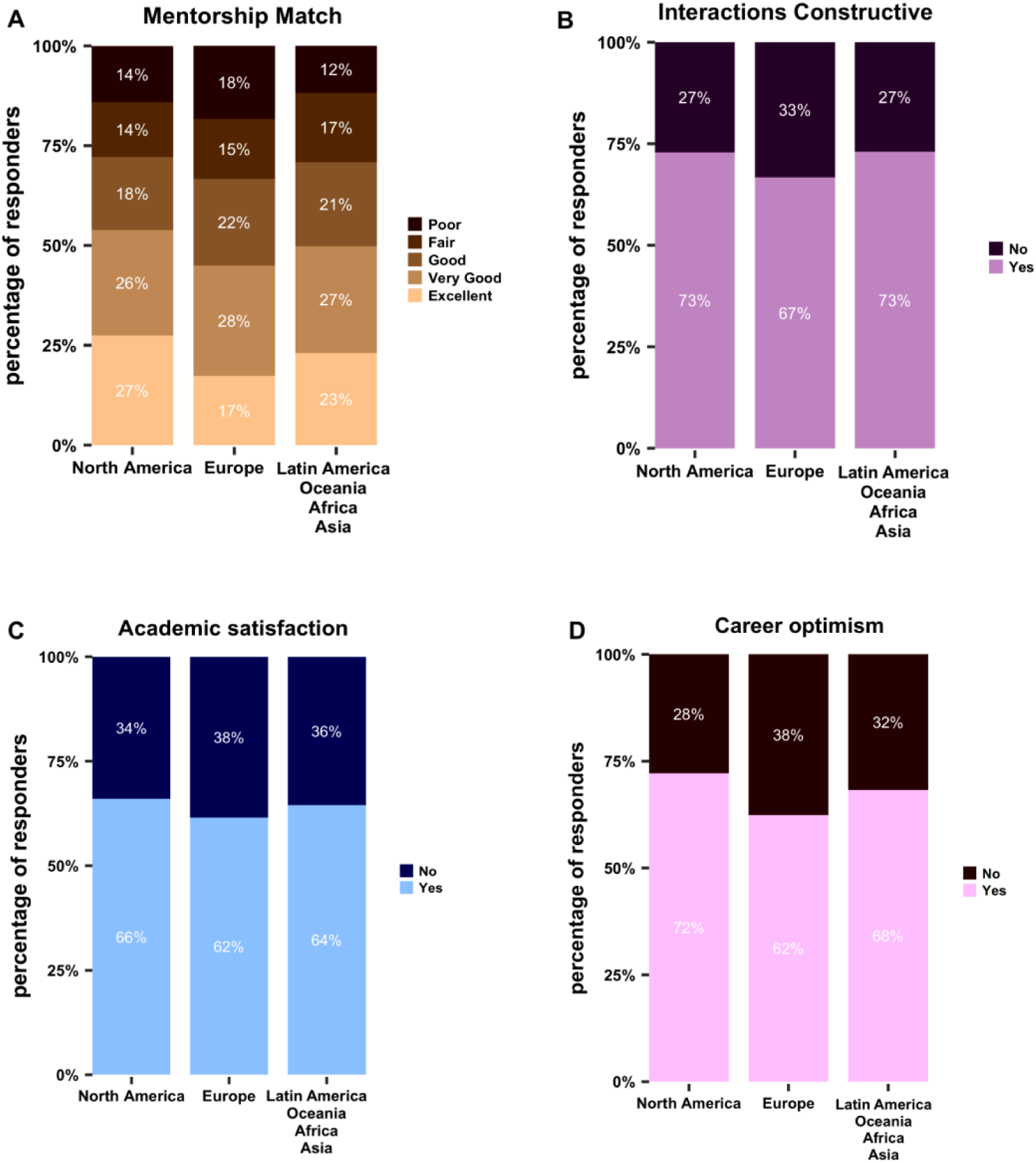
Mentorship interactions by trainee mentee geographic location. (A) Quality of mentorship match (Table S17) (B) Quality of mentorship interactions (Table S18) (C) Academic satisfaction (Table S19) (D) Career optimism (Table S20).

We further examined mentoring satisfaction for those working within their home countries (i.e., national or citizen mentees) versus those working outside of their home countries (i.e., international or non-citizen mentees). These results suggest disparities in access to mentorship opportunities and lower satisfaction for international trainees. Comparing mentees by their citizenship status, international mentees expressed less positive mentorship matches (Figure 5A, S4B, p<0.0001), fewer constructive interactions (Figure 5B, S4C, p<0.001), lower academic satisfaction (Figure 5C, S4D, p<0.001) and lower career optimism (Figure 5D, S4E, p<0.01).

**Figure 5.**
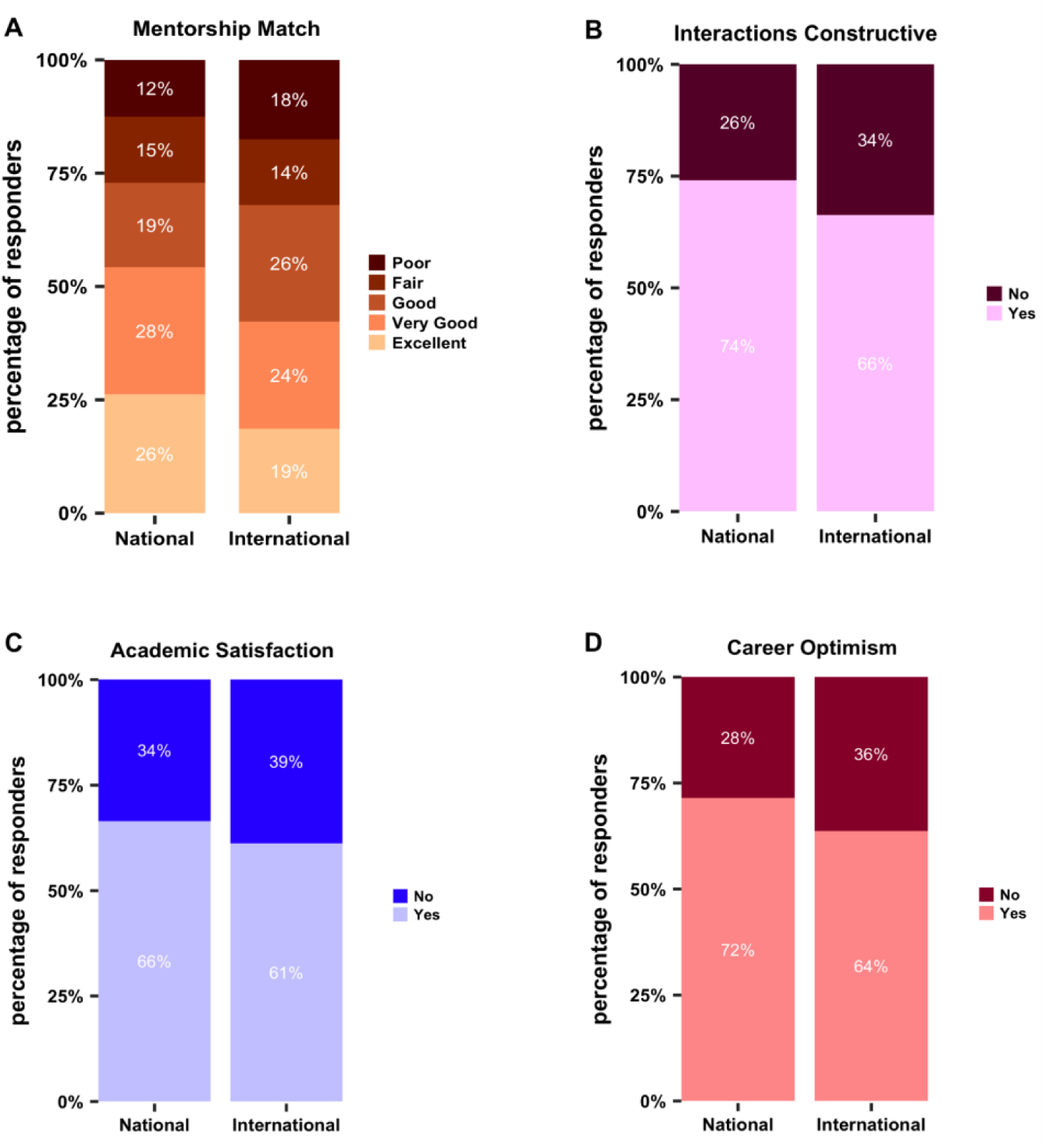
Mentorship interactions by trainee mentee citizenship status. (A) Quality of mentorship match (Table S21) (B) Quality of mentorship interactions (Table S22) (C) Mentee academic satisfaction (Table S23) (D) Mentee career optimism (Table S24).

We further sought to analyze trends in gender disparities in access to and satisfaction with mentors. In examining gender, men and women expressed similar satisfaction with mentorship matches, constructive interactions, academic satisfaction (Figure 6A-C, S2E, p=n.s., S3B, p=n.s., Figure 7A-B). Men expressed higher career optimism compared to women (Figure 6D, S4A, p<0.01). Our analysis of respondent citizenship status by gender, shows that a larger percentage of women were international researchers compared to men (Figure 7). Examining both gender and citizenship status, the survey results indicate higher dissatisfaction in mentoring experiences of international women trainees compared to national women, and national compared to international men (Figure 8A-D, 9A-D, S5A-F,H, 10E-H, S6A-E).

**Figure 6.**
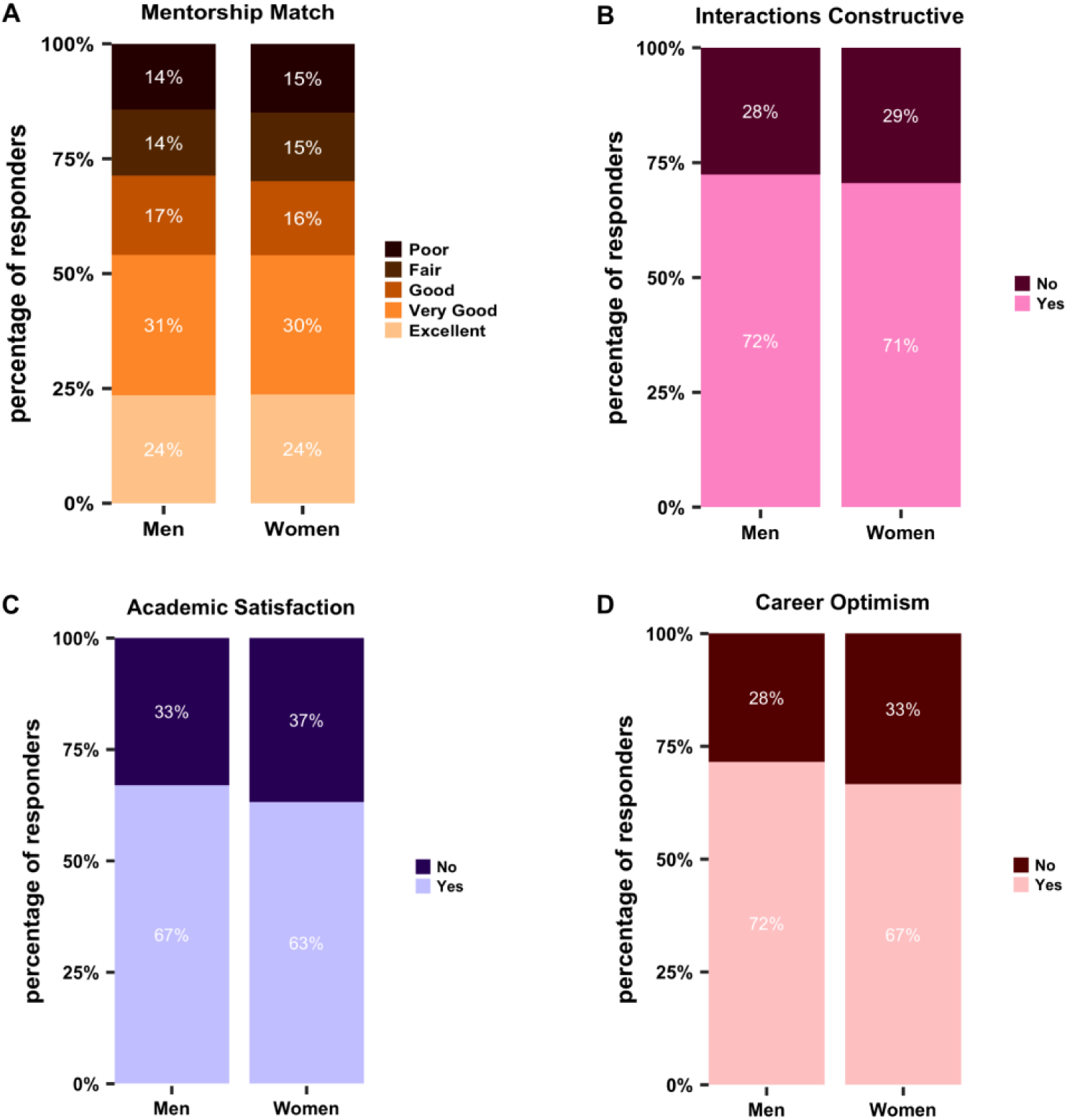
Mentorship interactions by trainee mentee gender. (A) Quality of mentorship match (Table S25), (B) Quality of mentorship interactions (Table S26), (C) Mentee academic satisfaction (Table S27), (D) Mentee career optimism (Table S28). Survey respondents were distributed across self-identified genders, with 38% identifying as male, 61% as female, and 1% identified as non-binary or preferred to not disclose this information.

**Figure 7.**
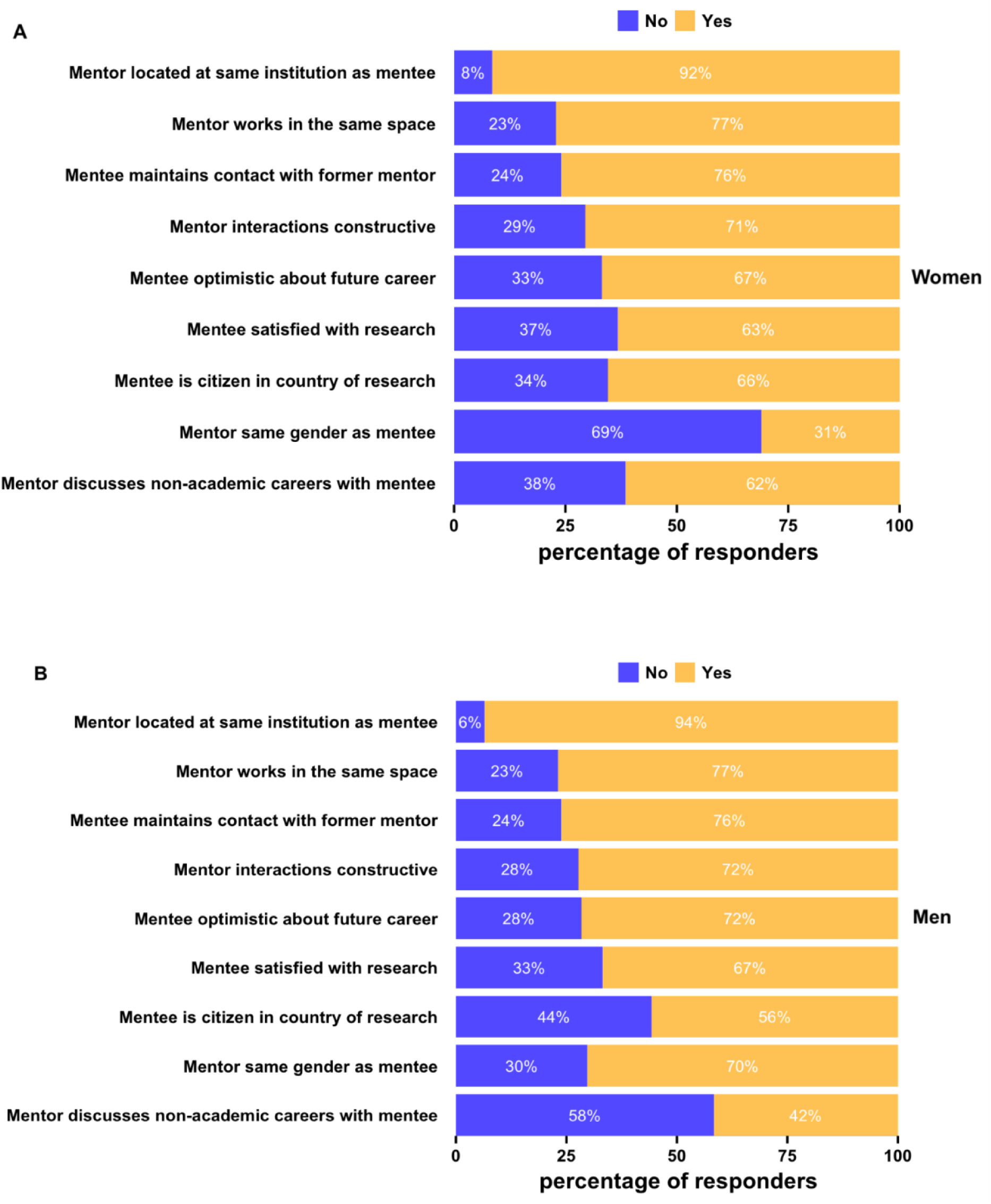
Mentee and mentorship characteristics by trainee mentee gender. Mentee characterization of mentor location (same department/institution or not), mentor physical location (same laboratory/team office space or not), mentor matching method, mentor accessibility, mentor gender (same as mentee or not), maintaining contact with former mentor(s), citizenship status in the country of research for (A) Women and (B) Men (Tables S26-S34).

**Figure 8.**
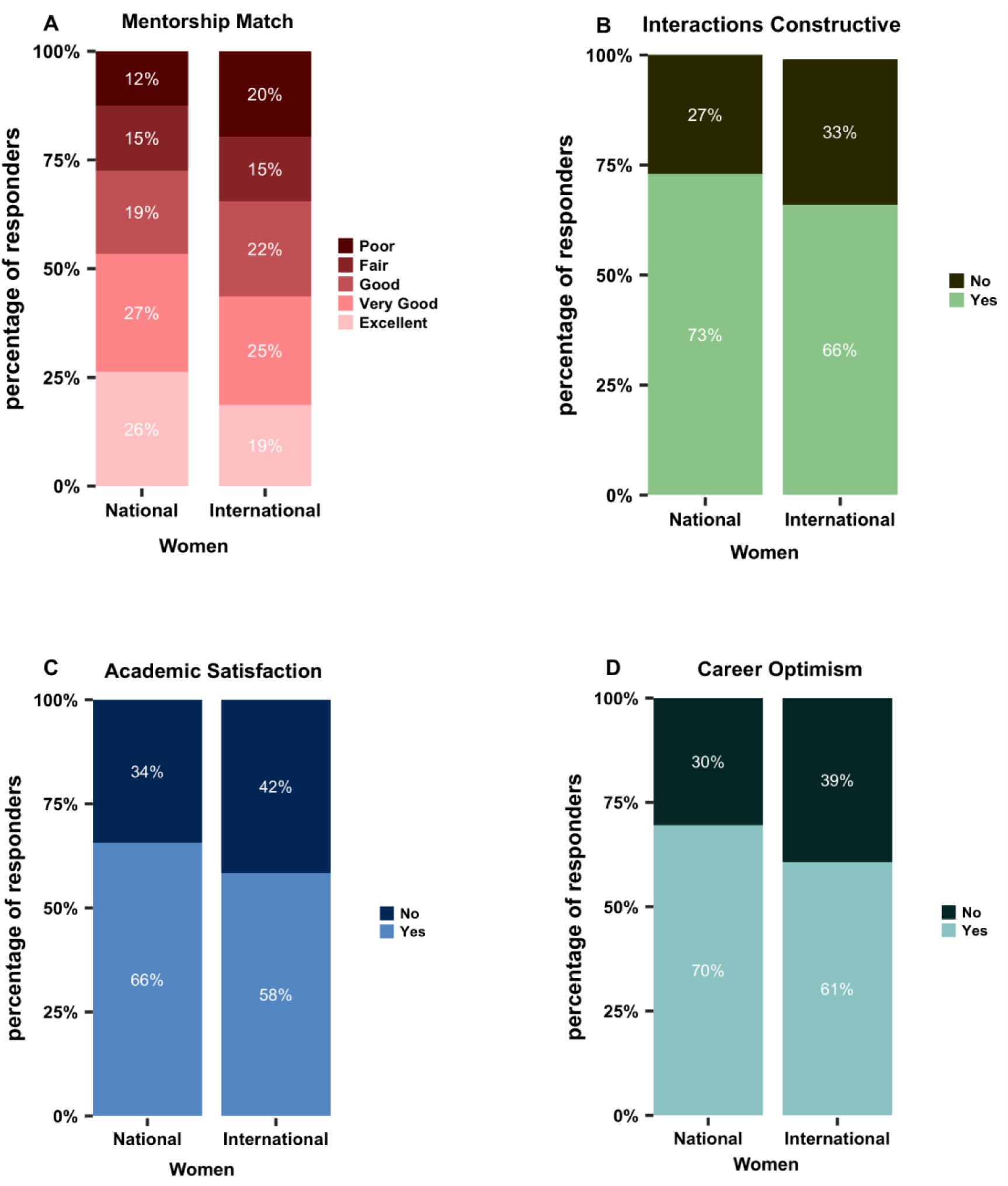
Women trainee mentee mentorship interactions by mentee citizenship status. (A) Quality of mentorship match (Table S37), (B) Quality of mentorship interactions (Table S38), (C) Mentee academic satisfaction (Table S39), (D) Mentee career optimism (Table S40).

**Figure 9.**
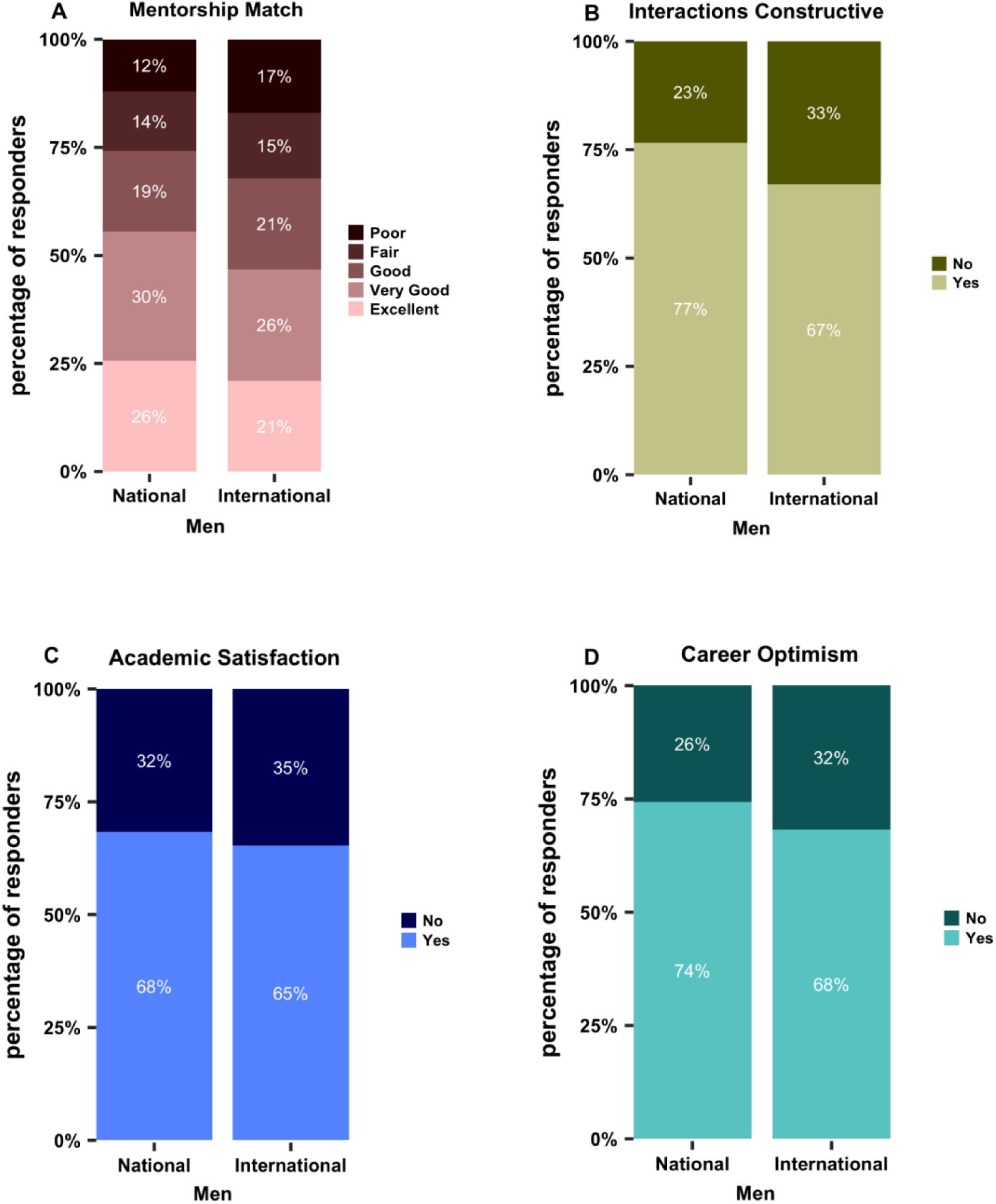
Men trainee mentee mentorship interactions by mentee citizenship status. (A) Quality of mentorship match (Table S37), (B) Quality of mentorship interactions (Table S38), (C) Mentee academic satisfaction (Table S39), (D) Mentee career optimism (Table S40).

We further asked survey respondents to rate the characteristics of their mentor and the extent to which their mentor met their expectations. This included questions regarding mentor interactions (e.g., used active listening, provided constructive feedback), mentor advocacy (e.g., acknowledged professional contributions, helped strategize and prioritize goals), and access to a hospitable working environment (e.g., decreased or eliminated workplace discrimination and harassment, valued and promoted diverse backgrounds, and acknowledged potential biases and prejudices) (Figure 10A-B). We did not, however, ask respondents to rank each of these mentor characteristics as valued in their mentorship interactions. Despite this, mentees indicated several areas in which mentors could improve. First, mentors should help mentees develop strategies to meet career goals (negatively perceived by 39% of women and 34% men). Second, mentors should help mentees negotiate a path to professional independence (negatively perceived by 40% of women and 35% of men). Third, mentors should consider bias and prejudice in mentorship interactions (negatively perceived by 42% of women and 35% of men). Finally, mentees would like mentors to work with them to set expectations for mentoring interactions (negatively perceived by 42% of women and 41% of men). The results show lower satisfaction of international women mentees compared to national men mentees in these categories (Figure S6A-E).

**Figure 10.**
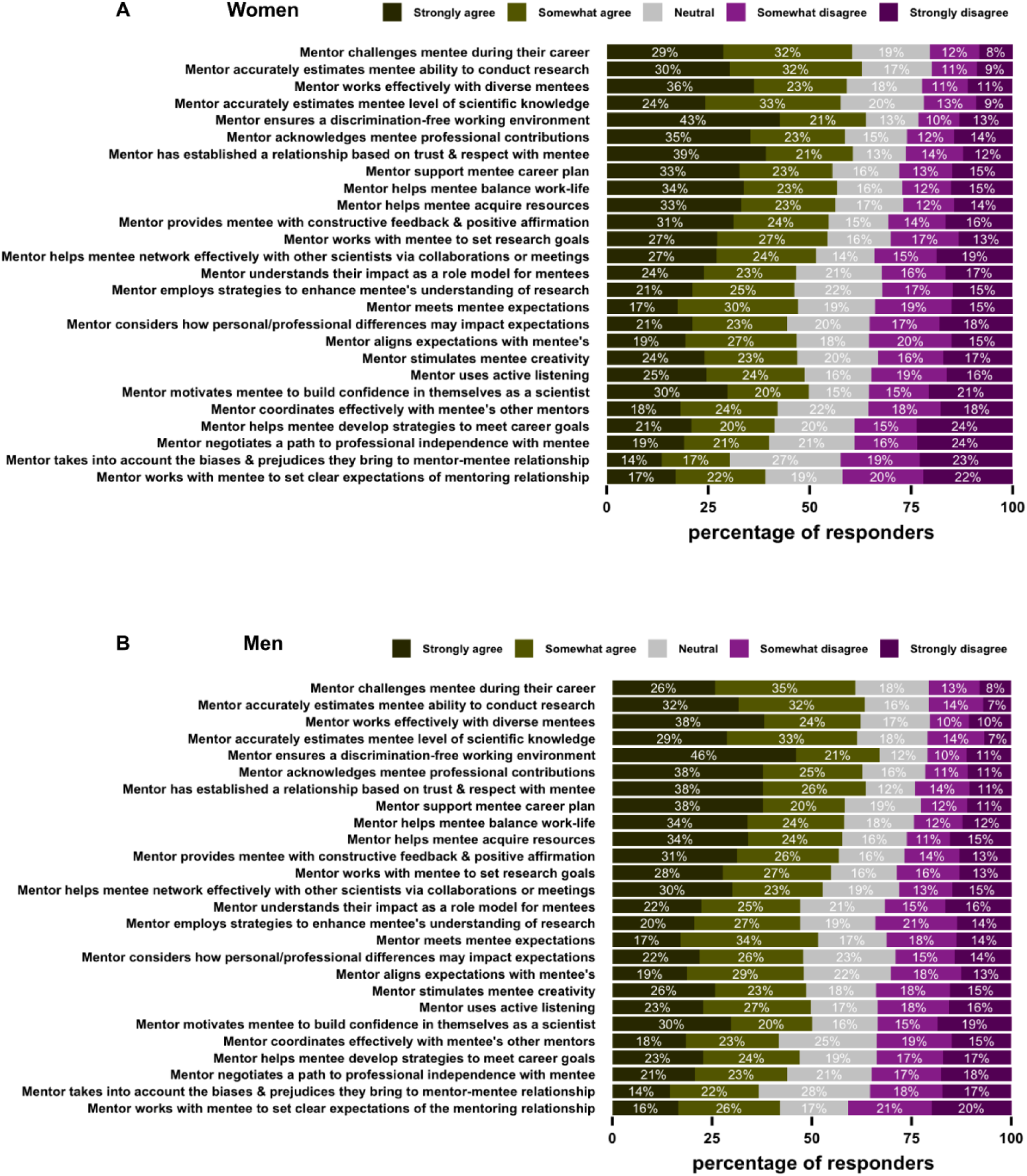
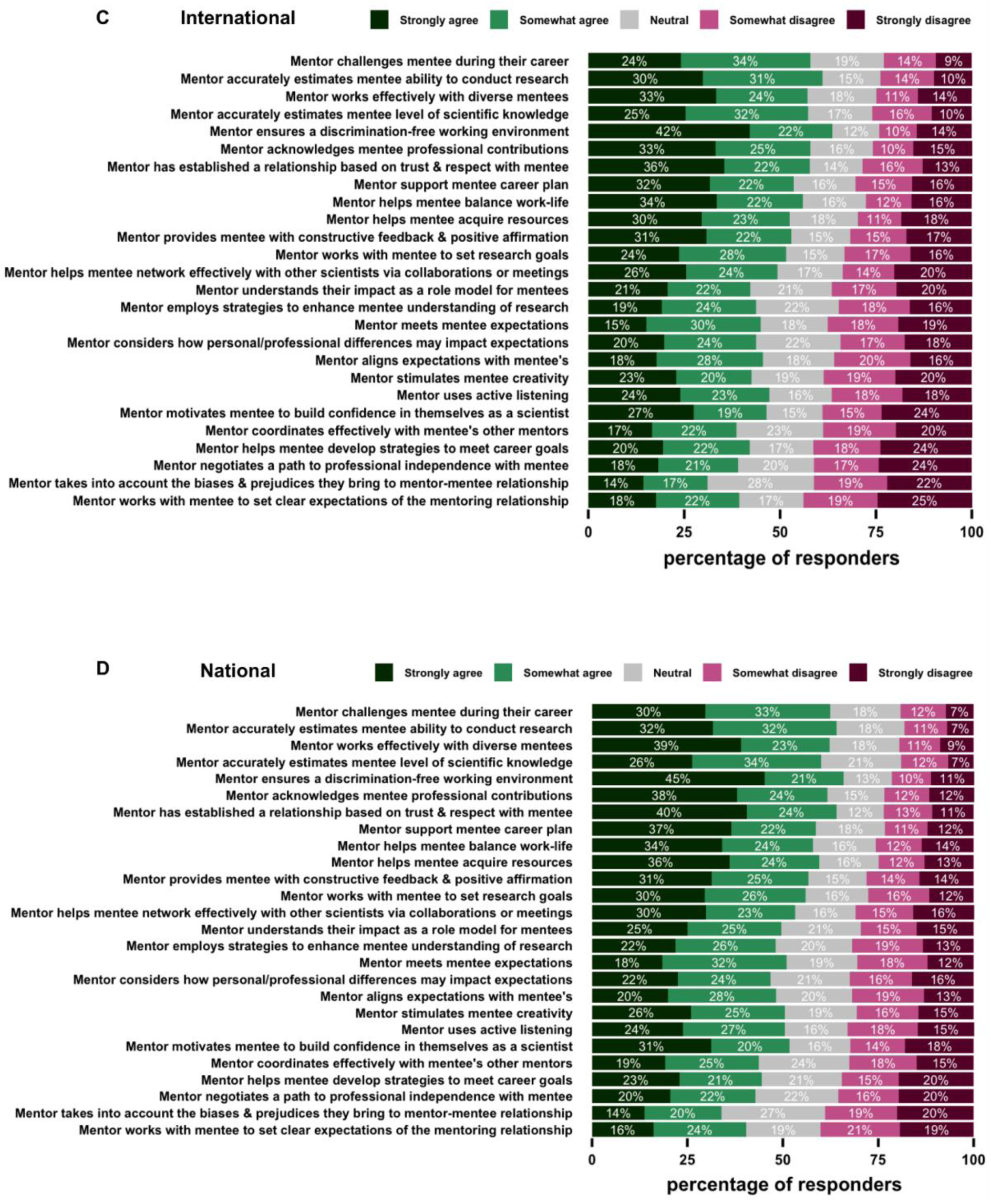
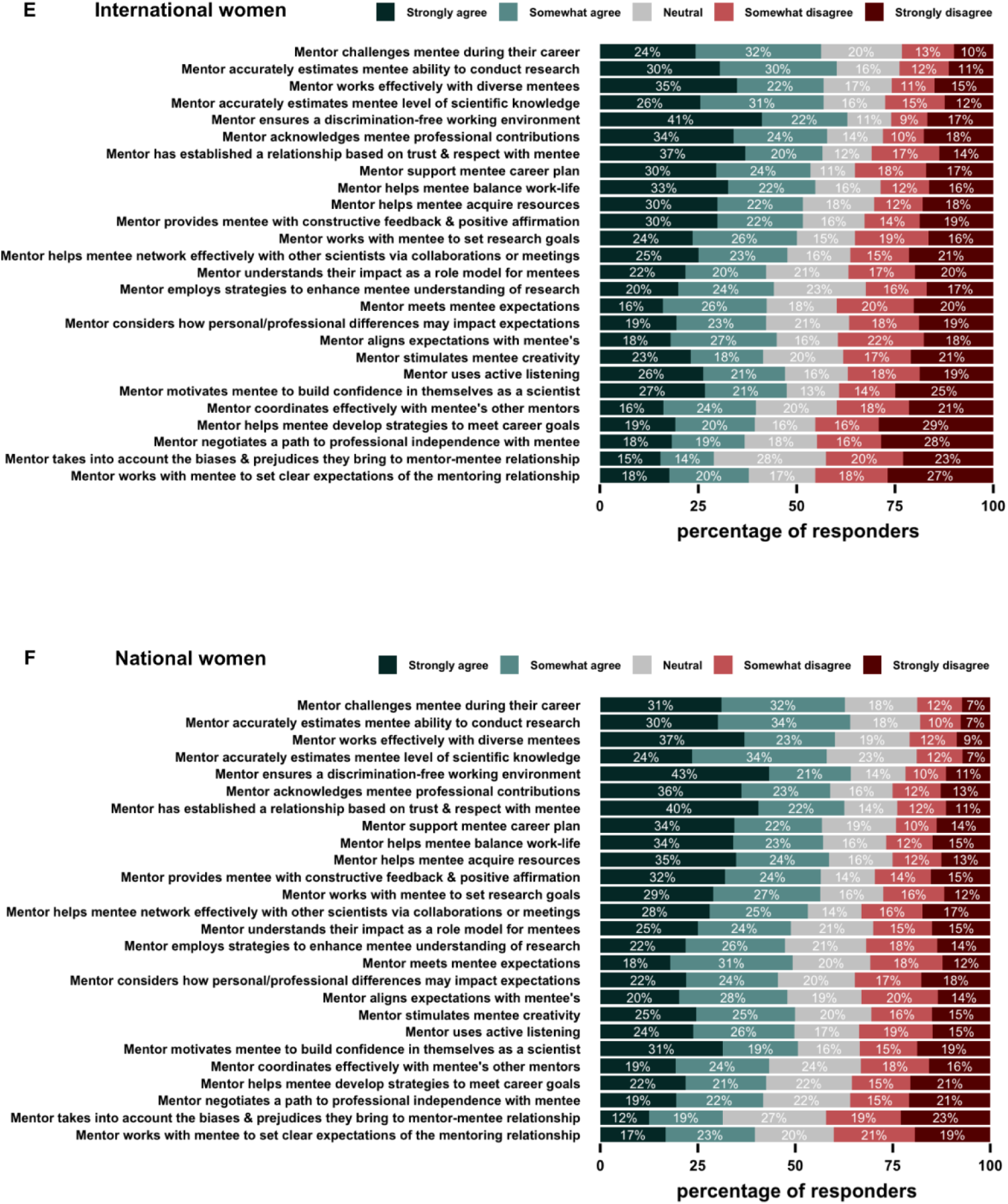
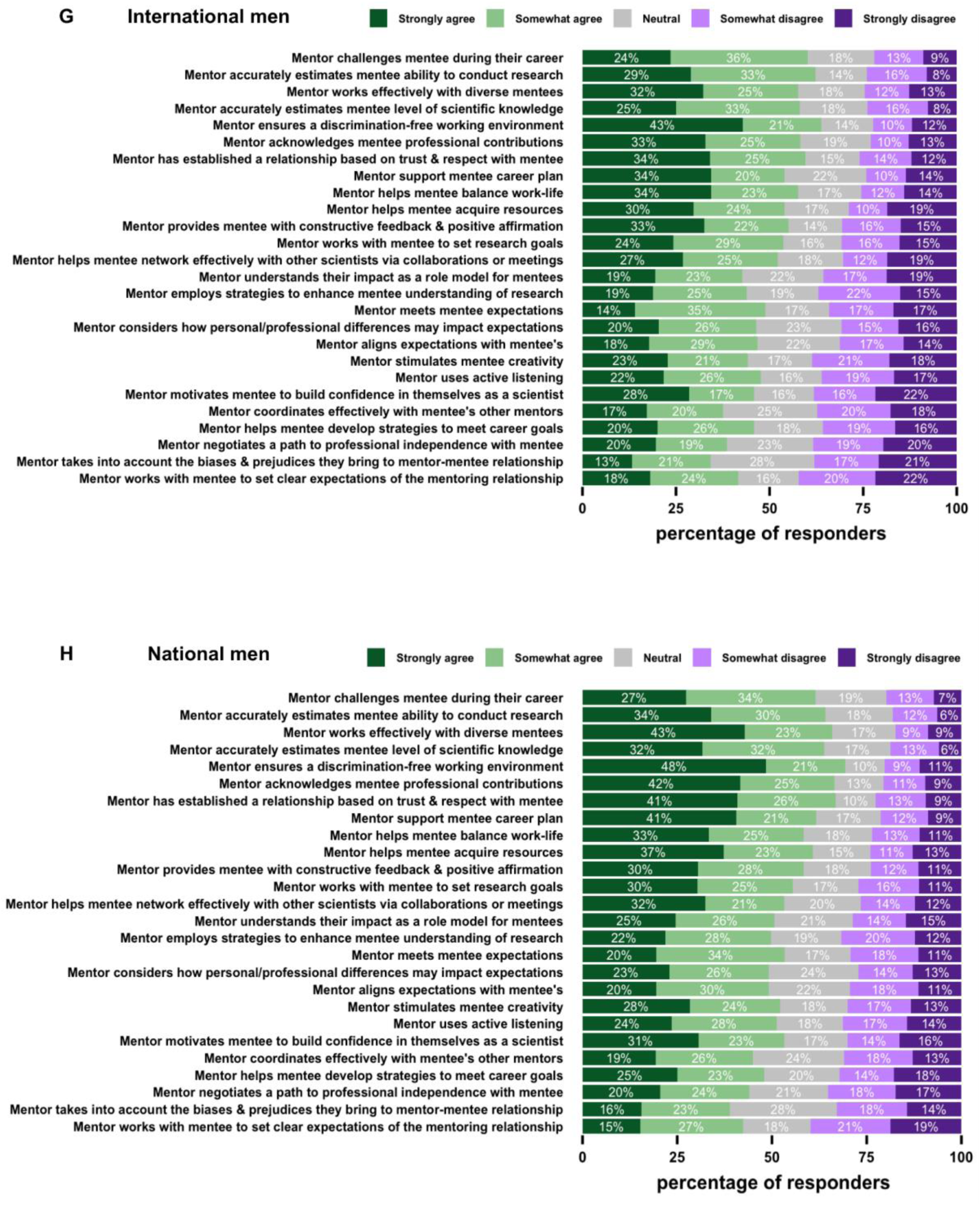
Trainee mentee valuation of mentorship interactions by gender and citizenship status. Mentee rating of mentorship characteristics on use of active listening (i.e., Identifying and accommodating different communication styles and employing strategies to improve communication with you), establishment of a relationship based on trust, quality of mentor feedback, quality of working environment, mentor valuation of colleagues with a diverse background, mentor’s consideration of biases, acknowledges mentee professional contributions, mentor’s helps in acquiring resources (e.g., grants, instrumentation, collaborations etc.), help in developing strategies to meet career goals, help with work-life balance, introduction to speakers and other professors at meetings and seminars and scientific society gatherings, working effectively with mentee whose personal background is different from their own (age, race, gender, class, region, culture, religion, family composition etc.), mentor’s help in developing strategies for mentee to better mentor their own mentees, overall extent of mentor meeting mentee expectations. (A-B) All respondents by gender, (C-D) All respondents by citizenship status, (E-F) Women respondents by citizenship status, (G-H) Men respondents by citizenship status (Table S41-S118).

### Analysis of qualitative responses

We asked respondents, in the form of long response questions, if their interactions with mentors were valuable. If not, we asked mentees to articulate what they were seeking in the mentorship relationship. Using these data, we summarized key mentorship features displayed by mentors who were noted as helpful, desired, unique, and needed (Figure 11). Many of the issues raised by mentees appear to be caused by poor alignment of expectations, including mentees’ desires for emotional support, professional development, career guidance, and support in attaining independence (Figure 13B). Through a compilation of responses from mentees on their mentoring relationships, we have identified common pitfalls, helpful mentorship traits and recommendations for various stakeholders as noted by trainee mentees.

**Figure 11.**
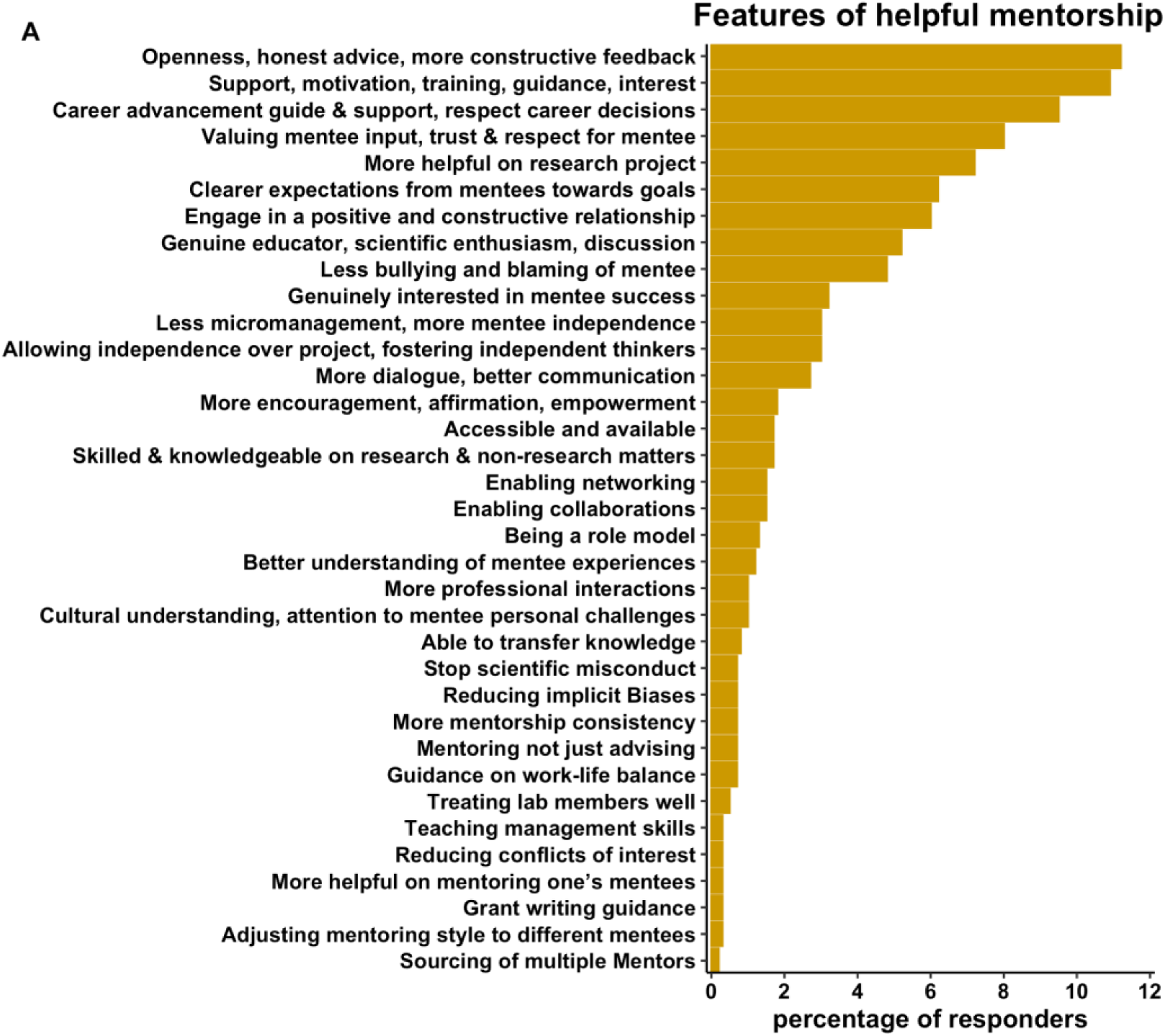
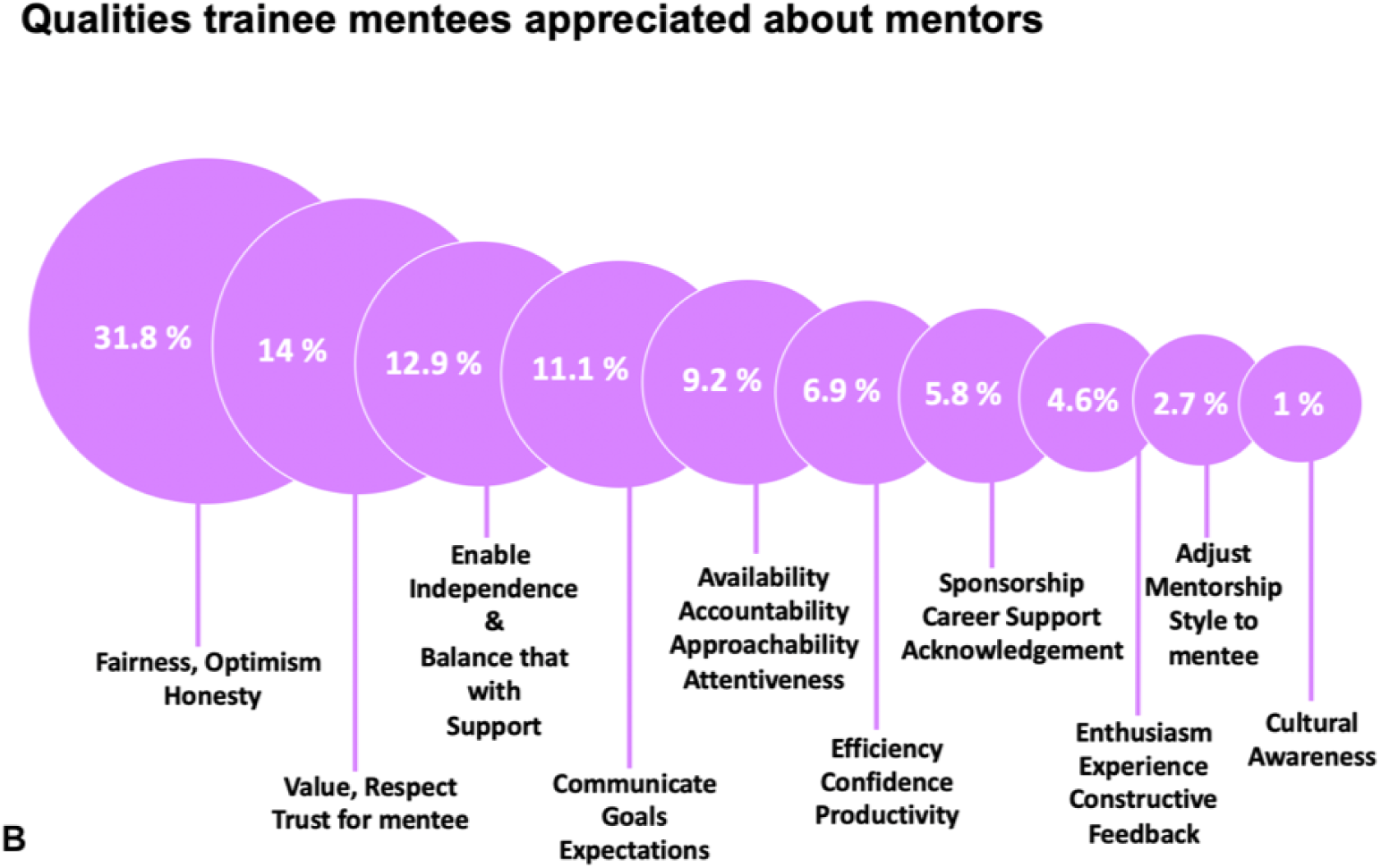
Analysis of qualitative comments on positive mentorship traits. Percentage of responses to (A) What do you like about your mentoring experience? (Total responses n=1,093), (B) If your interactions with your mentor were not deemed constructive, what were you ideally looking for? (Total responses n=604).

**Figure 12.**
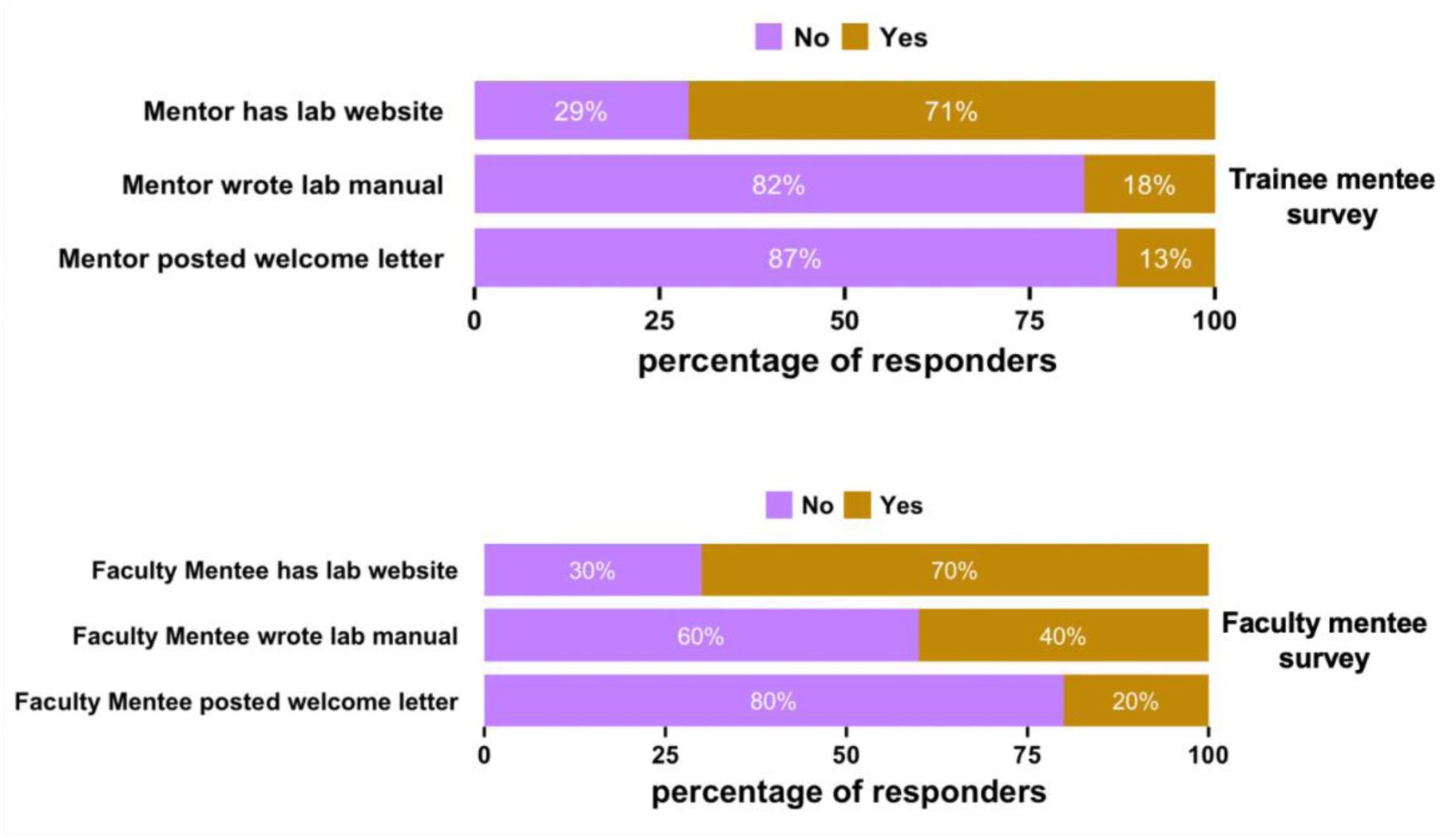
Improving mentorship initiation and interactions. Responses by all trainee mentee respondents and all faculty mentee respondents to questions on whether the trainee mentee’s faculty mentor or faculty mentee (in a faculty-to-faculty mentorship survey) created a lab website, wrote a lab welcome letter, wrote a lab manual (also known as lab culture expectations document) (Table S119-S124). Faculty mentee data for this figure panel originates from a faculty-to-faculty mentoring experiences survey (Sarabipour et al., 2023), shown here for the first time for comparison with faculty mentors of trainees in the trainee mentorship survey.

**Figure 13.**
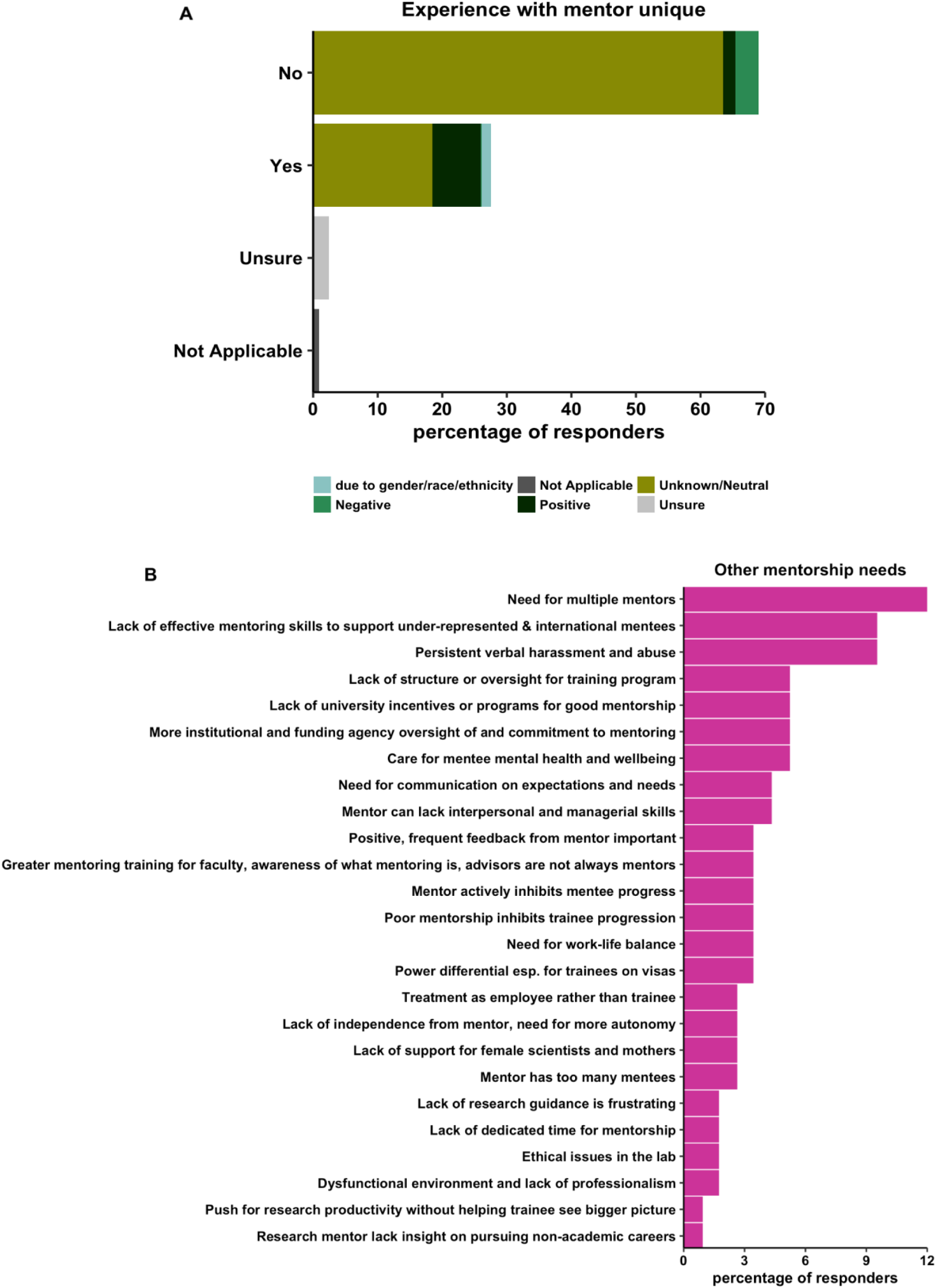
Analysis of qualitative comments on uniqueness of mentorship and other mentorship needs. Percentage of responses to (A) Do you feel your experience is unique in the lab (i.e., are you treated differently from other lab members by your mentor)? unique situations were cases where the respondent was the only trainee mentee (total responses n=1,027), (B) Other comments on mentorship (total responses n=116).

### Helpful mentorship traits as desired by trainee mentees

#### Genuinely supportive while providing guidance and mentorship

Amongst the most common helpful mentorship traits identified by trainees is genuine support from their mentors–not only of their scientific research, but also of their development outside of the lab. Many survey respondents noted that highly regarded mentors express genuine concern for the success and well-being of their mentees, showing an interest in who they are outside of their role in the laboratory. Many trainees found that supportive mentorship is highly motivational, instilling self-confidence in their abilities and their work. A number of trainees perceive that their mentor believes in their abilities even when they lack belief in themselves, which enables them to overcome impostor syndrome and grow scientifically in the lab. Similarly, mentees appreciate mentors who create a positive and helpful environment, helping them to feel that they belong both in the group as well as in research in general.

Mentees noted that mentors can show genuine support and guidance through a set of common traits, including being generally available, approachable, and attentive to the needs of trainees. Survey respondents appreciated their mentor being regularly available for unplanned short chats when mentees seek out help or advice on scientific and non-scientific topics. Trainees further appreciate mentors who offer their undivided attention during meetings, creating the mental space to think deeply about the work and ideas of the mentee. Some survey respondents acknowledged and appreciated their mentor’s patience and willingness to thoroughly explain concepts and goals for mentees still new to conducting research. Regardless of stage, trainees value mentors who are generous with their time to encourage and reassure mentees when they are on the right track. Similarly, they appreciate mentors who are kind and forthcoming about mentees’ shortcomings when trainees are not on the right track.

Trainees also commented on how their mentors can facilitate effective scientific mentorship. Many survey respondents appreciate mentors who address big questions in their field but still take the time to discuss how these questions fit in with the work being done in the lab. Many also positively commented on their mentor’s ability to simplify complex problems into manageable and actionable steps to allow the translation of these big picture ideas into meaningful progress in their day-to-day work. Beyond incorporating new ideas, trainees value insightful advice given from an experienced and wide knowledge base geared toward gaining the best results. Many trainees noted that guidance is most effective when given as needed, while simultaneously providing the trainee autonomy to direct their own day-to-day work. Finally, our survey respondents valued mentors who have solution-oriented approaches when things in the lab go awry, rather than focusing on blame or excuses.

#### Openness, honesty, and transparency

Trainees seek a mentor who has positive and honest conversations while maintaining a professional relationship. This allows trainees to express their views to have mutually open and constructive conversations. Mentees appreciate mentors who are direct, honest, respectful, and encouraging while lacking judgment. Survey respondents noted that great mentors maintain open and safe lines of communication and free expression of views surrounding scientific projects and goals. However, trainees also appreciate mentors who set clear, tangible expectations about progress and performance in the laboratory. Specifically, trainees desire mentors with an organized, strategic approach to projects who share these insights with mentees to facilitate a mutually beneficial ‘team science’ attitude. Survey respondents value mentors who engage frequently – answering questions, advising on approaches, and providing general guidance on how to advance projects and address struggles in the lab. Trainees seek a mentor who can challenge them intellectually and discuss big picture research ideas but remain open to discussions about new research directions, or simply makes time to creatively brainstorm about projects and ideas. Mentees value guidance in these discussions, but also seek freedom to raise concerns or bring forth opposing opinions about research directions and experimental design. Some trainees additionally seek mentors who can lend specific technical advice on experimental design and troubleshooting, including hands-on training within the laboratory. Regardless of the topic on which mentees seek advice, they appreciate feedback that is accurate and constructive, particularly when delivered positively and in a manner that clearly signals that the mentor is trying to be helpful and supportive.

Coupled to this, trainees value honest feedback and engagement around work expectations. This includes consistent and timely feedback on written documents (e.g., fellowship proposals, manuscripts, and job application essays) but also mentors’ willingness to listen to mentee problems and questions and providing perspective and solutions on how to address these issues. These conversations could extend beyond their projects. Indeed, mentees noted that open, positive, and honest conversations about a variety of topics – from benchwork to professional development – were highly valued from a mentor. Mentees appreciate mentors who maintain high expectations in regard to mentee performance in the lab while simultaneously providing the resources and support to allow mentees to excel and reach these expectations.

#### Valuing, trusting, and respecting mentees and their experiences

Our survey respondents describe ideal mentors as those who value the opinions and ideas of their mentees. Some noted mentors who are as enthusiastic about mentee ideas as the mentor’s own ideas, instilling motivation and confidence in their trainees. Numerous trainees desire a mentor who trusts and respects their ideas and expresses confidence in mentees while simultaneously supporting trainees should the need arise. Trainees noted that this respect promotes intellectual discourse between the mentor and mentee, allowing mentee growth and eventual independence. Over time, this allows mentees to grow into colleagues capable of making original intellectual contributions to the research process – a critical step in their scientific growth. Mentees also value mentors who request mentee input and offer the mentee freedom to express their opinions before making important decisions.

Mentees appreciate mentors who respect their individual experiences and actively work to let the mentee know that they belong in their group. Mentees value mentors who appreciate and understand that mentees have a life and responsibilities outside of the lab, and who take steps to accommodate their individual needs. Beyond simple accommodations, mentees value mentors’ advocacy and assistance with both personal and professional problems. Specific examples cited by survey respondents include trainees valuing mentors’ efforts to navigate a poor relationship with a different mentor or taking action to fight harassment on behalf of the trainee.

#### Enabling the transition to independence while maintaining support

One recurrent theme identified by trainees as key to their success in the laboratory is that their mentor actively worked to enable their transition to independence. Our survey respondents noted that mentors should encourage their mentees to perform research – alone or in collaboration with other groups – but should continue to guide the mentee on critical aspects of the work. Mentees noted that this guidance needs to be dynamic and adjusted throughout their scientific growth. For instance, graduate students within their first years of training may need more specific help on technical aspects of their work, appropriate experimental design, or guidance as to which literature is most relevant to their project. Indeed, multiple comments from our survey respondents noted frustrations with how much independence their advisors expected, leading mentees to feel alone and discouraged at their progress and prospects. Alternatively, multiple postdoctoral fellows that took our survey found their mentors to be over engaged, disallowing independent thought and micromanaging their projects. The qualitative responses seem to indicate that the level of independence desired by a given trainee not only correlates with their seniority and experience in the lab, but also depends on their personal preferences. Thus, mentors should not only check in with trainees often about the level of independence expected but should also plan to grow to trust the mentee’s independence in the laboratory over time.

During their transition to independence, mentees noted that they appreciated the ability to perform work on their own, even when they felt slightly underprepared to do so. Allowing trainees to become independent may inevitably be accompanied by errors and mistakes in the lab, which allow mentees to learn and grow. Mentors who respond to such mistakes with patience and a positive mindset are highly valued by mentees at all levels. Mentees also appreciate being empowered to make their own scientific decisions so long as they have reasonable justification. Our survey respondents note that they value the opportunity to do their own reading, make their own presentations, and draft manuscript papers independently to feel a level of ownership over their research directions and projects. Finally, mentees value growing into mentors themselves by working with and training students within the lab or gaining experience through teaching others.

#### Enabling career advancement and supporting diverse career options

Trainees rely heavily on their mentors for career advancement advice and opportunities, and desire mentors who will support them in both endeavors. To enable career advancement, respondents hoped that their mentor would help to establish network connections with other labs (both nationally and internationally) for research collaborations. Mentees wished to be included in discussions in early stages of project planning with these collaborators to establish relationships with potential leaders on relevant projects within the field. Giving mentees a voice in the process is well-aligned with respecting and trusting mentee opinions and helping them grow independence but also allows them to be personally invested in projects. Beyond collaborations, mentees hoped for mentors who would promote their work, both by presenting mentee findings at conferences and allowing the mentee to present their work to a wider scientific community at conferences and workshops. Mentees appreciate mentors who encourage and support grant and award applications, and who generally support the ambitions of mentees overall. As these ambitions will be distinct for each trainee, mentees value a mentor who accommodates mentee career plans by aligning their long-term goals and choices when prioritizing projects and professional development activities. Furthermore, mentees appreciate acknowledgement and support from their mentors to pursue both academic and non-academic positions.

### Common mentorship pitfalls noted by trainee mentees

#### Lack of trainee knowledge on importance and benefits of mentorship

Some respondents noted that they didn’t know much about mentorship before entering a graduate training position. This includes lack of knowledge in desired mentor qualities and being uninformed of appropriate expectations of mentorship. Some mentees noted their opinions concerning their mentor came from observing how other labs are managed or the relationships that their colleagues had with their faculty mentor. Due to the high variability in mentorship styles and training environments, this informal gathering of information did not always translate to the best fit in mentorship for the trainee. These responses indicate a need for mentee training in seeking and managing mentorship interactions.

#### Treatment as employee rather than mentee

Based on qualitative responses, the majority of survey respondents regarded their research advisor or principal investigator (PI) of their project as their key mentor. Some respondents noted that they were not treated by their research advisor as a mentee but rather as an employee. These mentees felt that their mentors used them more as a source of labor and set of lab hands rather than a mentee with goals and aspirations. Others reported that their mentors seemed to care more about their own career and publications than the development of the mentee. Respondents noted that this led to mentees seeking opportunities for appropriate mentorship and support elsewhere.

#### Mentor often absent or has too many trainees

Respondents noted that some mentors were inaccessible due to frequent absences or insufficient time to devote to mentorship. Some noted their mentor was too busy for their students, unreachable by email and distracted during face-to-face meetings. Some respondents noted that their mentors were not completing basic commitments such as being prepared for their students’ degree milestones such as thesis meetings and completion. Some respondents noted their mentor had insufficient mentoring abilities because they had too many students. Similarly, some mentees noted that the size of a mentor’s lab has an important bearing on the quality of mentorship, as mentors running large labs may be stretched thin for the time and resources required to successfully guide junior scientists. Some respondents noted that they actually preferred a more independent mentorship experience as postdoctoral researchers, but noted they still require feedback and bigger picture guidance during this training period (also see Figure 2B).

#### Lack of mentee independence caused by mentor micromanagement

In contrast to mentees feeling abandoned and unsupported, some respondents noted that they needed more independence and trust from their mentors. These mentees desired mentors who were open to new ideas and allowed mentees to develop independent research projects, apply for their own grants, and transition to independent academic jobs. Some noted that even if the mentee generated the ideas for the experiments, the mentee had no independence because the mentor micromanaged what the mentee was doing within the lab. Some noted that their mentor did not support any endeavor outside of the primary research endeavors of the mentor and would only agree to allow the mentee to perform other science-related work (e.g., write grants or perform outreach) begrudgingly. This may be due to mentors delivering a message against spreading too thin or about alternative activities serving as diversions that prevent progress to degree milestones. Some noted having little freedom in work or thought due to mentor-issued demands for specific endpoints in their work. Others noted that they did receive support for developing their own research trajectory, but that this did not happen until the mentee asked for it directly.

#### Power imbalance between mentor and mentee

Some survey respondents noted that many significant challenges arise due to the unequal power dynamics present in many academic mentoring relationships. Mentees often endure non-ideal mentoring situations (from minor annoyances to major burdens) because they need their mentor’s approval for grant submissions, publications, and letters of recommendation. Respondents noted that mentees in poor mentoring situations were not able to self-advocate if they were treated poorly. This can be especially difficult for international scholars, who are particularly vulnerable due to their visa status being dependent on their mentor’s willingness to keep them employed in their lab.

#### Insufficient professional respect and boundaries

Respondents noted that they hoped for a professional and supportive relationship with their mentor. However, mentees reported that some mentors harbor behaviors that would be considered unprofessional at best and outright abusive at worst. Some mentees noted that their mentor treated them more like a friend than advisee or attempted to become too involved in the mentee’s personal life, creating an uncomfortable dynamic. Respondents instead hoped for interactions where their personal lives didn’t interfere with the professionalism expected in the mentoring relationship. Other survey respondents noted their advisor created a dysfunctional and unproductive lab environment, with reports of negativity, overt bullying (e.g., closed-door abusive meetings) and exploitation from their mentors. Respondents desired a mentor who showed trust and encouragement in the mentee rather than a continuous push to produce the data that matches the mentor’s desired hypotheses. Beyond the research, some mentees noted mentors who engaged in personal attacks rather than constructively criticizing the work. These behaviors highlight the need for expectations of professionalism to be made explicitly clear to mentors by their institutions.

#### Lack of consideration for mentee work-life balance, mental health and well-being

Respondents also noted that some mentors expect the mentees to dedicate their life to research. In such cases, most mentees are constantly under stress and high pressure, creating a poor atmosphere in the lab. Some respondents noted that they were punished for taking time off on weekends despite the fact that they actually did work on most weekends. Respondents hoped for a mentor who recognizes the importance of work-life balance and allows mentees to take a reasonable number of breaks and vacation time.

#### Insufficient communication and unclear expectations for mentees

Respondents hoped for clearly defined goals and guidelines from their mentor, particularly with regards to reasonable expectations for their day-to-day goals and overall progress. Some mentees noted that their expectations did not match with what their mentors desired, resulting in frequent misunderstandings. Some noted that they were met with pressure and unreasonable deadlines towards the end of their training. For example, some mentees report that their mentor set unrealistic timelines for completing their work while subsequently ignoring their finalized publications. Some trainees note that their mentors do not have (or at least do not effectively communicate) a set of goals for their lab to work toward, leaving the mentee without much direction.

#### Poor support for mentees with non-traditional backgrounds or experiences

Some respondents noted that they were discriminated against and scientifically dismissed due to overt gender or racial discrimination, or due to the fact that they were completing their training in a foreign country. Some respondents noted differences in how their mentor related with other graduate students from the same lab, indicating bias. Some noted that they felt that they were treated differently from national students based on their international status. Some noted that certain personal or health issues, such as having a child or managing a chronic illness, negatively affected their experience because there was little to no support from their mentor or institution for managing these challenges. Some noted that being mentored by mentors with similar backgrounds was helpful in these situations, and enhanced mentee confidence and abilities.

#### Lack of discussions on diverse careers

Many mentees found that their mentor does not provide significant guidance regarding non-research and non-academic careers. This could be because the mentor does not have the appropriate experience to speak to these opportunities, but could also be due to the fact that they do not support these career paths. Importantly, our data show that 60% of mentees reported that their mentor had not discussed non-academic careers with them (Figure 7A-B). Regardless of the reasons for this lack of discussion, mentors should respect and support the career paths chosen by their mentees. Respondents seemed discouraged by this lack of support for non-academic career paths, noting that mentors should be aware that due to paucity of permanent academic positions, the majority of their trainees will not end up in careers as full-time academic faculty.

#### Mentorship quality may vary for mentees of the same mentor

We asked respondents if their experience with their mentor was unique (Figure 13A). Many found this question challenging, as some aspects of their mentorship relationship seemed unique, while others overlapped with the experience of those in their laboratory. For instance, some trainees noted that there should be inherent unique interactions that are tailored to the specific project, career stage, and goals of a specific mentee. However, other aspects may not be unique, such as mentor availability to mentees. Some respondents noted that it was difficult to compare their mentorship experience with other lab members, or that they had select characteristics (e.g., gender differences) that made it hard to assess ‘uniqueness’ without potential biases. Some mentees found their relationship with their mentor isolating due to a persistent sense of not belonging in the research group or overt favoritism for other members in the lab.

#### Persistent exhibition of favoritism

A number of respondents noted a strong culture of favoritism cultivated by mentors. In some laboratories, this is perceived to occur due to differences in experience level, scientific skill set, or background. Some mentees found that their mentor pays the most attention to those who are the most productive, despite the fact that those who are struggling would likely most benefit from additional mentoring. Similarly, some mentees noted being given more or less independence in the pursuit of their projects based on their skill level or how closely the project intersects with their mentor’s own interests and skills. Some mentees reported unequal opportunities are given within their lab for co-authoring publications or presenting their work. Interestingly, at least one mentee noted receiving favoritism in the lab, which led to unhealthy behaviors and poor work-life balance. Thus, trainees reported negative experiences with favoritism on either side of the experience.

Other mentees reported problematic favoritism in their labs that were rooted in mentor bias. These include examples of gender bias, with women treated with less respect and courtesy, given less support, or being asked to handle a heavier administrative burden than their male counterparts. Others note that their nationality created bias, with international students reporting that they were ignored due to communication barriers or aspects of their cultural background which make them less likely to self-advocate. Finally, multiple mentees felt that those in the lab that desired to stay in academia were treated better than those who were interested in pursuing non-academic career paths.

#### Research ethics issues

A number of respondents noted highly concerning behavior in their laboratories, witnessing data manipulation, rigging, and corruption during their research training. Some noted that this was due to careless behavior of their mentor, who was focused on professional advancement rather than on the robustness and reproducibility of the research in their group. Respondents noted mentors who pushed mentees to accomplish undoable projects or to ‘fit the data to the model’ rather than accept the conclusions of the actual experiments. Respondents expressed a desire for a mentor who conducts science in the most ethical way possible, who is open to change, and who is able to recognize and acknowledge when they are wrong about scientific or non-scientific issues.

#### Comparing the trainee mentee survey with the faculty mentee survey

We asked faculty mentees in a previously published survey (Sarabipour et al., 2023) on whether they created a lab website, posted a welcome letter (Bennett et al., 2014) on their website and whether they composed a lab expectations manual for their group. We asked the same questions this time about mentors from trainee mentees in the trainee survey. The findings of both surveys consistently show that while many faculty have a lab website, composing a welcome letter for prospective trainees and a lab manual are still not commonly practiced in research environments (Figure 12). The trainee survey responses on rating of mentorship characteristics shows the lack of mentor communication on clear expectations for the mentorship relationship as the most prominent source of trainee mentee dissatisfaction (Figure 10). Thus, initiating dialogue and setting expectations through a lab website, welcome letter and a laboratory manual could improve the mentorship experiences of trainees.

## Recommendations for improving mentorship for trainees

### Mentors can help improve and optimize mentorship for mentees

#### Balancing equal treatment and tailored mentorship for mentees

Analysis of the qualitative responses from our survey revealed that mentees have a strong desire for both equal treatment as well as tailored and personalized (i.e., equitable) mentorship. While these aspects of mentorship may seem in conflict with one another, mentors can cultivate both traits in their laboratories and in their individual mentoring relationships. Mentors should promote a healthy lab culture that respects work-life balance and promotes a positive work environment. Equal treatment of mentees can be achieved by allotting similar time for meetings and project planning for each lab member (e.g., via weekly one-on-one meetings) and by giving attention to all projects being worked on in the lab. Furthermore, mentors should actively work to hold each lab member to the same professional standards within what is reasonable for the career status of the mentee. While maintaining similar expectations, communication, and respect for all trainees, mentors should also prepare to adapt their mentoring style to the unique needs of each member of their laboratory. This includes understanding the individual motivations and long-term goals of each trainee and adapting project goals and professional development to align the direction of the laboratory with the goals of the trainee. Mentors should understand that there is not one single path to becoming a scientist, and that trainees have different backgrounds and lived experiences that impact their work in the lab. Furthermore, respondents noted they appreciated when their relationship with their mentor evolved over time, suggesting mentors should anticipate dynamic and changing interactions with their mentees.

#### Setting clear expectations for individual mentees and laboratory practices

Mentors should be prepared to clearly communicate goals, guidelines, and expectations for the trainees in their lab. This should include frequent communication on scientific approaches and the current status of experiments, but should also extend to the state of the lab or long-term goals relevant to funding applications and publication strategies. Principal investigators should also work to communicate a set of standard expectations for certain laboratory roles and how these expectations may differ, for instance, between graduate students and technicians. Principal investigators can achieve this through the creation of a laboratory handbook, which can include details of such expectations, as well as basic lab information including policies on time off, working hours, and other administrative tasks. The lab manual additionally can provide standard forms for the laboratory (e.g., Individual development plan (IDP) or professional development templates, meeting note templates, etc.) and can comment on the culture and values cultivated by the research team. Having such information both communicated and easily accessible can be highly useful for promoting best and consistent laboratory practices for mentees. Additionally, mentors can assess the opinions of their lab members regarding aspects of research and research culture by running anonymous lab surveys. Mentees also need to work to manage their mentor’s expectations and match these with their own goals. Mentorship should be normalized in academic environments for trainees.

#### Supportive of mentee well-being and mental health

Recent studies suggest that a substantial fraction of mentees struggle to achieve work-life balance or have issues with their mental health (Evans et al., 2018; Hyun et al., 2006; Loissel, 2020). Mentors should take steps to proactively support their trainees and to promote their work-life balance and mental health. Mentors should minimally have an understanding of common mental health issues and work to support mentees with these struggles. Actionable ways to support mentees include having flexibility for meeting times, work schedules, and understanding the need for time off. Mentors should also be aware of resources on their campus that can assist in mental health and well-being, and should promote these resources to their trainees. Finally, mentors should communicate with their mentees about hard work and high expectations, while understanding the threshold at which such expectations may lead to burnout. As all mentees will have different backgrounds and needs in regards to their mental health, mentors can work to both determine and honor these individual needs amongst the mentees within their laboratory.

#### Career development guidance and support

One of the most important roles a mentor has is to prepare their mentees and enable them to achieve their career goals by supporting their professional and career development. Mentors should take the time to understand the long-term goals of their trainees or assist their mentees in figuring out what career paths interest them. One mechanism that may assist with this is to complete and discuss Individual Development Plans (IDPs) with their trainees at least annually. This gives mentees a formalized opportunity to discuss their ideas for professional development and to allow the mentor to advise on these goals. Upon establishing these goals for their mentees, mentors can then use their network to find opportunities for their mentees to connect with established members of the fields that interest them. Mentors should be open to careers outside of traditional academic roles, and can connect mentees with recent program graduates or former postdoctoral fellows within their department that have moved into non-academic roles. For mentees that would like to stay in academia, mentors can find ways to promote mentee work through presentations at conferences or establishing collaborations with others in their field.

#### Improve mentorship and training conditions to increase trainee career satisfaction

It is worth noting that poor mentorship practices have far reaching effects on the personal and professional trajectories of mentees in academia. This largely is driven by an inherent hierarchy that exists between faculty members and their mentees, with mentors in a position of power that can result in the exploitation, abuse, or general mistreatment of trainees. Mentors should not only recognize that such a power dynamic exists, but should work to minimize the negative connotations and actions that can result from such a dynamic. This includes ensuring that the mentor’s own behavior is appropriate and professional with their direct mentees and other trainees at their institution and beyond. Beyond their own behaviors, mentors should understand their obligations in reporting inappropriate interactions and behaviors in academia.

### Departments, institutions, and funders can promote excellence in trainee mentorship

#### More structure or oversight of mentorship programs

Some respondents noted that their graduate program was dysfunctional, and there were no good means to approach a bad, but not abusive, mentor relationship. Respondents noted they found it frustrating that there seems to be no mechanism to ensure that professors are performing their basic functions as mentors, such as providing feedback on dissertation chapters, showing up on time to qualifying exams, and meeting with trainees regularly. Some noted that complaints regarding their mentor were ignored or swept under the rug by departmental leadership. Respondents noted that university policies could influence the quality of mentorship, for instance by including such information in faculty evaluations or within their tenure packages. Others noted that mentorship practices should be reviewed by institutions. Furthermore, mentees of all levels need to be able to confidentially report research ethics or harassment issues to departmental or university leadership without fear of repercussions or retaliation.

#### Greater mentorship training and accountability for faculty

One recurrent observation from our study is the lack of infrastructure for mentorship oversight or training programs. Furthermore, the results from our survey suggest a need for greater awareness of what constitutes ideal mentoring practices among principal investigators. Respondents noted that leadership and mentoring courses should be required of incoming tenure track assistant professors, who often have not received such training despite beginning a career that relies heavily on successful mentorship practices. Classes on mentoring and leadership can formalize some of the core expectations of what mentors can and should do to effectively train mentees in their laboratory, and such expectations can and should be shared with mentees to create clear communication about these subjects. Such training should be supported at the department and institutional level, or could be supported by funding agencies within grant applications. Our survey indicates that mentorship practices could be improved by implementing formal review policies by institutions and funding agencies to provide oversight for mentor engagement and support. Finally, department and institutional leadership could proactively initiate and facilitate positive discussions between faculty and trainees through informal networking events to establish potential informal mentorship connections (Figure 14).

**Figure 14.**
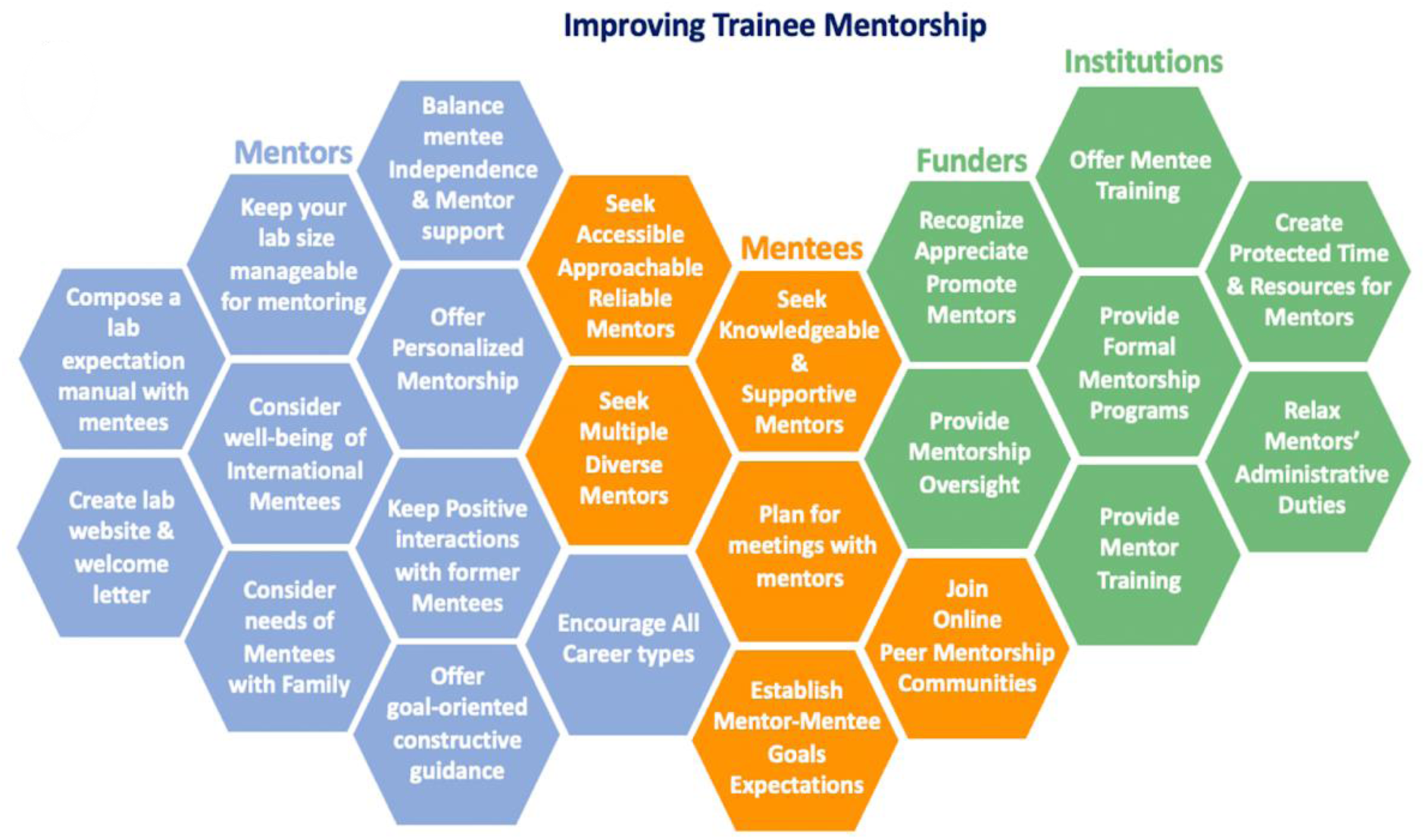
Summary of key recommendations for trainee mentees, mentors, institutions and funders.

## Discussion

We analyzed findings of a survey of 2,114 graduate and postdoctoral researchers from over 65 countries to characterize features of mentorship interactions. One of the key issues noted by respondents was unavailability of mentors for sufficient training and mentorship of their trainees. Mentors should be honest about their capacity to provide meaningful guidance and support to trainees in their lab, either limiting the number of scientists in their lab or establishing supportive networks within their laboratories, such as teams of senior and junior lab members, to facilitate mentee success. Conversely, a large lab size opens new opportunities for co-mentorship, which can be highly beneficial for trainees at all career stages. Principal investigators should recognize the need to alter their mentorship strategy based on growth or shrinkage in their labs and should consider that time devoted to mentorship may be limiting as they grow their lab as their current mentorship model may not scale accordingly. This suggests that faculty mentors leading large labs may need a *team* to provide adequate mentorship, which may come from a lab culture of co-mentorship.

Mentors can run annual or semi-annual anonymous lab surveys to gauge what lab members think about various aspects of research and laboratory culture and how mentors can best support them (Vosshall, 2022). A lab welcome letter (Bennett et al., 2014) and laboratory manual or handbook, complete with expectations, could be composed by the principal investigator to help with trainee management and mentorship (Hainer et al., 2020; Maestre, 2019; Masters and Kreeger, 2017). Mentees need to also make an effort towards managing academic work and life expectations, particularly as the progress toward independence in the lab (Bartlett et al., 2021; Martin and Grimes Stanfill, 2023; Sarabipour et al., 2021).

Some mentorship issues trainees reported in this survey are shared with our recent faculty mentee survey (Sarabipour et al., 2023). Consistent with our faculty mentee survey, the need for multiple mentors is important for trainees, as it can be challenging if not impossible to find expertise and guidance on all issues in a sole advisor. However, it can be difficult for trainees to initiate interactions or source mentorship with faculty. Department leadership, scientific societies, graduate training programs, and graduate or postdoctoral associations all could play a positive role in facilitating such interactions or providing diverse perspectives to help trainees’ scientific and career development (Bielczyk et al., 2019; Choi et al., 2019; Hillebrand and Leysinger, 2023; Treasure et al., 2022). However, it is worth noting that mentees should be proactive in seeking and securing the mentorship they need. Such proactive actions will not only boost their confidence but also secure guidance from a diverse set of mentors ranging from peers to faculty (Figure 14). The effectiveness of mentorship can be different from the perception of mentorship by the mentee; for instance, mentees may not feel that mentorship is effective when in fact it is or vice versa (i.e., can feel it’s great when it is not actually helpful to them in shaping their career). To assess this, mentorship should be evaluated longitudinally according to mentee perception against some objective measurement (Andrews and Chilton, 2000). Defining objective measures can be a challenge, but mentors and institutions could, for instance, assess dropout rates from high-stress or toxic environments as a proxy for student dissatisfaction. Furthermore, our findings identify a blueprint for trainee mentorship needs in research environments.

In the area of mentoring trainees, the pitfalls noted by survey respondents may be driven by both a lack of mentorship training for PIs, as well as systemic challenges. For instance, tying graduate and postdoctoral training to specific grant funding may incentivize some of the identified pitfalls. We hope that these benchmarks are discussed by research groups and departments to clarify best mentorship practices for mentors and mentees. Ongoing conversations on mentor and mentee roles, expectations and best practices can further alleviate issues before they arise.

### Study strengths and limitations

In this work, we report the findings of a large and independent survey of trainees (predominantly graduate students and postdoctoral researchers) in research environments worldwide. While some institutions may run climate and mentorship surveys internally, their findings are seldom made public, suggesting a need for data curation on the broad needs of trainee mentorship. As our survey was completed before the COVID-19 pandemic, mentoring perceptions may have changed in some aspects that are not reflected in our survey. The COVID-19 pandemic also may have reduced outside mentoring, at least for those who were funded to travel to conferences and informally seek external mentorship. Our responses are highly enriched in researchers in the life and biomedical sciences and medical research areas currently residing in North America or Europe, and conclusions about mentorship practices across other disciplines may differ substantially. Furthermore, the survey was presented in English, influencing our reach. Nations with English-speaking researchers, on social media and English-speaking universities, have the highest number of researchers per million inhabitants, so it is common for similar research culture surveys to have responses concentrated in North America and Europe. Despite this commonality, this concentration of responses influences the inferences we can make about mentorship relationships worldwide, particularly in geographic regions not well represented in our survey responses. We did not collect data on the race or ethnicity of respondents and therefore cannot know how this may have influenced some aspects of the findings of our survey. In future surveys, it would be interesting to compare an expansion to non-tenure track research assistant professors and research scientists as their growth and mentoring path is often poorly defined, with some faculty career expectations (e.g., writing grants), and some postdoctoral training expectations (e.g., driving key grant-funded projects).

## Materials and Methods

### Survey Materials

The text for the survey used in this work is included in the supplemental material. A Google form was used to conduct the survey and was distributed on various social media platforms including academic Slack groups the Grad Student Slack and the Future PI Slack and via academics on X (formerly Twitter) worldwide. The survey was distributed from March 2019 to March 2020 and contained both scaled-response and open-ended questions. Respondents could choose to not respond to any of the questions (leave the response section blank) thus partial and complete survey participation was included in data analysis. The respondents to the survey were asked to self-report, and the information collected was not independently verified.

### Data Analysis

Microsoft Excel, RStudio and the ggplot package (Wickham, 2016) and the eulerr package (Larsson and Gustafsson, 2018) were used to graph survey findings shown in Figures 1-11A, Figures 12-13 and Supplementary Figure S1. The qualitative survey comments were categorized by theme (themes, keywords and context) describing each comment and the frequency of comments pertaining to a particular theme are summarized in Results and Discussion. Bar charts were generated from distinct themes in qualitative responses (Figures 11 and 13). Figures 11B and 14 were created using Microsoft Powerpoint. For statistical analyses, Prism 9 was used to perform Ordinary one-way Analysis of variance (ANOVA) (Figure S2A-E, S3A-B, S6A-E), two-tailed Mann-Whitney U (Wilcoxon Rank Sum) test (Figure S2F, S3C, S5A-H, S4A-E) and the Tukey multiple comparisons test (Figure S2A-E, S3A-B, S6A-E).

### Data availability

The authors confirm that, for approved reasons, access restrictions apply to the data underlying the findings. Raw data underlying this study cannot be made publicly available in order to safeguard participant anonymity and that of their organizations. Ethical approval for the project was granted on the basis that only aggregated data is provided (as has been provided in the supplementary tables, with appropriate anonymization) as part of this publication.

### Statement of Ethics

This survey was conducted by researchers and educators listed as authors on this publication, affiliated with universities in the United States in an effort to identify the challenges that researchers in training positions (graduate students and postdoctoral researchers) face in receiving mentorship from senior colleagues. The authors respect the confidentiality and anonymity of all respondents. No identifiable private information has been collected by the survey presented in this publication. Participation has been voluntary and the respondents could choose to stop responding to the survey at any time. The survey has been verified by the Johns Hopkins University Institutional Review Board (IRB) as Exempt IRB-00012432.

## Supporting information

Supplementary Information_Trainee_Mentorship_Sarabipour et al 2023

## Author contributions

SS and SJB designed and distributed the survey. SS, SJB and NMN performed the data analysis. SS performed the tabulation and visualization. SS and NMN wrote and reviewed the manuscript. SS, SJB, NMN, CTS, AWBF, AI, KC wrote, edited and reviewed the manuscript.

## Conflicts of Interest

The authors declare no competing interests.

## Acknowledgments

The authors thank all the early career researchers worldwide who took the time to respond to the survey and offer their valuable input in support of this work. The authors thank Feilim Mac Gabhann, Richard Sever, Paul Macklin, Gunnar Blohm, Christian Frezza, Iain Cheeseman, Bradley Wyble, Uri Manor, Harmit Malik, Stephen Klusza and Inez Lam for their valuable comments on an earlier version of this manuscript and the early career researchers of *eLife* community 2018-2020 program for their support of this endeavor. NMN was supported by National Science Foundation grant #2327631.

## Notes

### Competing Interest Statement

The authors have declared no competing interest.

### Summary of Updates

In this revision, we have expanded on our analysis of data presented in Figures 5,6,8,9 and 10. New analysis are included in the supplementary information as Figures S2,S3,S4,S5 and S6 and Tables S125-S129.

