## Supplementary Information_Trainee_Mentorship_Sarabipour et al 2023 for "Insights from a survey of mentorship experiences by graduate and postdoctoral researchers"

### Table of Contents

**Survey Questions……...........................................................................................................2**
**Supplementary Figures……………………………………………………………………….9**
**Supplementary Tables .......................................................................................................15**

### Survey Materials

#### Trainee Survey questions

The purpose of this anonymous is to gain an overview of mentee's perspectives on the quality of mentoring received from their mentor(s) in research environments. The aim is to collate the results and use the findings to produce recommendations.

Our definition of a mentor includes: A Mentor is someone who guides trainees through their scholarly training in higher education institutions or other industry or government research environments. A Mentee is a graduate or postdoctoral trainee.

You are the Mentee. You may have research and non-research mentors. You may have one mentor or a mentorship team (Mentors).

A mentor is different from a manager. If you have multiple mentors, please focus on the one mentor that covers your most important requirements.

If you would like to comment on one of your previous mentors (Graduate school mentor, first postdoc mentor) as well as your current mentor, please fill the survey separately so that we can hear about your previous experiences as well.

Thank you

*Required

1) Please select how much you like independence in your job*. Mark one option only. (*Required).

No guidance or input                Very heavily managed

1    2    3    4    5

2) Overall, to what extent do you feel that your current mentor is meeting your expectations? (If you have multiple mentors, please focus on the one mentor that covers your most important requirements) *

3) Your Mentor's Communication Skills (If you have multiple mentors, please focus on the one mentor that covers your most important requirements).
Please rate how skilled you feel your mentor is in each of the following areas: [We understand that you can only speak from your personal experience. Please try to rate a skill whenever possible, reserving the 'not observed' category for cases where you have no basis for assessment].

Rating scale:

1    2    3    4    5

Strongly Disagree                Strongly Agree

A. Has established a relationship based on trust and respect with you

B. Uses active listening (Meaning Identifying and accommodating different communication styles and employing strategies to improve communication with you)

C. Coordinates effectively with other mentors with whom you work

D. Works with you to set clear expectations of the mentoring relationship

E. Provides you with respectful, positive affirmation & constructive feedback

F. Your Mentor's commitment to your (mentee) Scientific Growth

G. Challenges you during your career

H. Aligns his/her expectations with your own

I. Considers how personal and professional differences may impact expectations

J. Works with you to set research goals

K. Ensures a working environment free from discrimination and harassment & values

L. Accurately estimates your level of scientific knowledge

M. Accurately estimates your ability to conduct research

N. Employs strategies to enhance your understanding of the research

O. Motivates you to build confidence in yourself as a scientist

P. Stimulates your creativity

Q. Acknowledges your professional contributions

4) Your Mentor's Support of your Professional Development Support your career plan?

R. Helps you develop strategies to meet career goals?

S. Negotiates a path to professional independence with you

T. Takes into account the biases and prejudices s/he brings to your mentor/mentee relationship

U. Works respectfully and effectively with mentees whose personal background is different from his/her own (age, race, gender, class, region, nationality, culture, religion, family composition etc.)

V. Helps you network effectively with other scientists via collaborations or meetings?

W. Helps you balance work with your personal life (for example: take family and/or vacation time off)

X. Understands his/her impact as a role model for you

Y. Helps you acquire resources (e.g., grants, fellowships, etc.)

Z. Discusses non-academic career options with you for future/upon graduation? (e.g., Industry or government research jobs, jobs in publishing industry, science communication jobs, Teaching positions, Science policy jobs...) Mark one option only.

Yes
No

**About You (the Mentee): Professional Background**
5) Which category below describes you best? * Mark one option only.
Undergraduate student
Medical or health professional student
Graduate student (Masters or PhD)
Assistant scientist
Associate scientist
Clinical fellow
Postdoctoral researcher
Assistant professor (Tenure-track)
Associate professor (Tenured)
Full Professor (Tenured)
Other:

6) What type of advanced degree do you hold? [check only One] * Mark one option only.
High school Diploma
Bachelor's degree
Master’s degree
PhD
Professional degree (e.g., MD, DDS, RD, PT, PharmD, etc.)
Both PhD or Masters and professional degree (MD/PhD, MD/MPH, PharmD/MS, etc.)
Other:

7) Which category(s) best describes the focus of your research? [check all that apply] * Choose all that apply.*
Basic (Fundamental) research in Life Sciences
Basic research in Physical Sciences
Engineering
Theoretical/Mathematical research
Translational research (specify type: T1, T2, etc.)

Clinical research
Behavioral research
Field/applied research
Social Sciences
Environmental Sciences
Humanities
Medicine
Psychology
Neuroscience
Other:
If you answered "Other" to the previous question please provide your field of research here:

8) Is the majority of your work in the wet-lab/at the benchtop? * Mark one option only.
Yes
No
Both wet lab and computational work
Not Applicable to my field

9) What type of institution do you study or perform your research at? *Mark one option only.
Academic
Industry
Government
Other:

**Mentor(s) background**
10) How many research mentors do you currently have (i.e., your current advisor in your lab/department/program or other professors that provide mentorship)?

11) Is your main mentor located in the same department/Institution as you? Mark one option only.
Yes
No

12) How long have you been working with your current mentor? Mark one option only.
Not Applicable
Less than one year
1-2 years
3-4 years
5 years or more

13) How were you originally matched to this mentor? Mark one option only.
Voluntarily (I chose the mentor)
I was assigned to the mentor.
It was a combination of both.
By technical field
By Age
At random

Other:

14) Are you the same gender as your mentor? Mark one option only.
Yes
No

15) Do you share a workspace or have an office close to this mentor (so that you have unplanned meetings or encounters)? Mark one option only.
Yes
No

16) How frequently do you have formal meetings with your mentor? Mark one option only.
Weekly
Every 2 weeks
Monthly
Scheduled as necessary
a few times a year
Never

17) What format(s) do you use to communicate with your Mentor? Choose all that apply
Face-to-Face meeting
Telephone
Email
Web/Video conferencing (e.g., Skype, Zoom)
Social networking sites
Other:

18) Does your current lab have a website? Mark one option only.
Yes
No

19) Has your mentor posted a lab welcome letter for prospective students on your lab website? Mark one option only.
Yes
No

20) Has your mentor written a lab manual for current lab members? Mark one option only.
Yes
No

21) What has been the most helpful method for you in getting advice from your mentor? Mark one option only.
Face-to-Face meeting
Telephone conversations
E-mail communication
Web/Video conferencing (e.g., Skype, Zoom)
Other:

22) Please rate the quality of the match between you and your mentor. Mark one option only.
Excellent
Very good
Good
Fair
Poor

23) Do you find your interactions with your Mentor constructive? Mark one option only.
Yes
No

24) If you answered “NO” to the previous question what type of mentoring relationship were you looking for? What were you hoping to achieve with your Mentor?

25) If you answered “YES” to the previous question, what do you like in your current mentoring relationship?

26) Do you feel your experience is unique in the lab (Are you treated differently from other lab members by your mentor)?

27) Do you feel happy and satisfied about your current research career? Mark one option only.
Yes
No
Please rate your happiness and satisfaction about your current research career?
Mark only one oval.

| | | | | |
|---|---|---|---|---|
| 1 | 2 | 3 | 4 | 5 |
| very Dissatisfied | | | | very Satisfied |

28) Do you feel optimistic about your future career? Mark one option only.
Yes
No

Please rate your optimism about your future career? Mark one option only.

| | | | | |
|---|---|---|---|---|
| 1 | 2 | 3 | 4 | 5 |
| very Negative | | | | very Positive |

29) Do you maintain contact with your former mentors? Mark one option only.

Yes
No

**Your (Mentee) Demographics**

30) Your Gender. Mark one option only.
Female
Male
Prefer not to say
Female non-binary
Male non-binary

31) What is your age? Mark one option only.
<21
21-25
25-30
30-35
35-40
>40

32) Are you a citizen in the country you conduct your research? Mark one option only.
Yes
No

33) Which country do you study/perform you research at currently? *
Answer:---------------------------------

34) Other Comments about the survey or your experience that was not addressed in the Questions?
Long Answer:---------------------------------

### Supplementary Figures

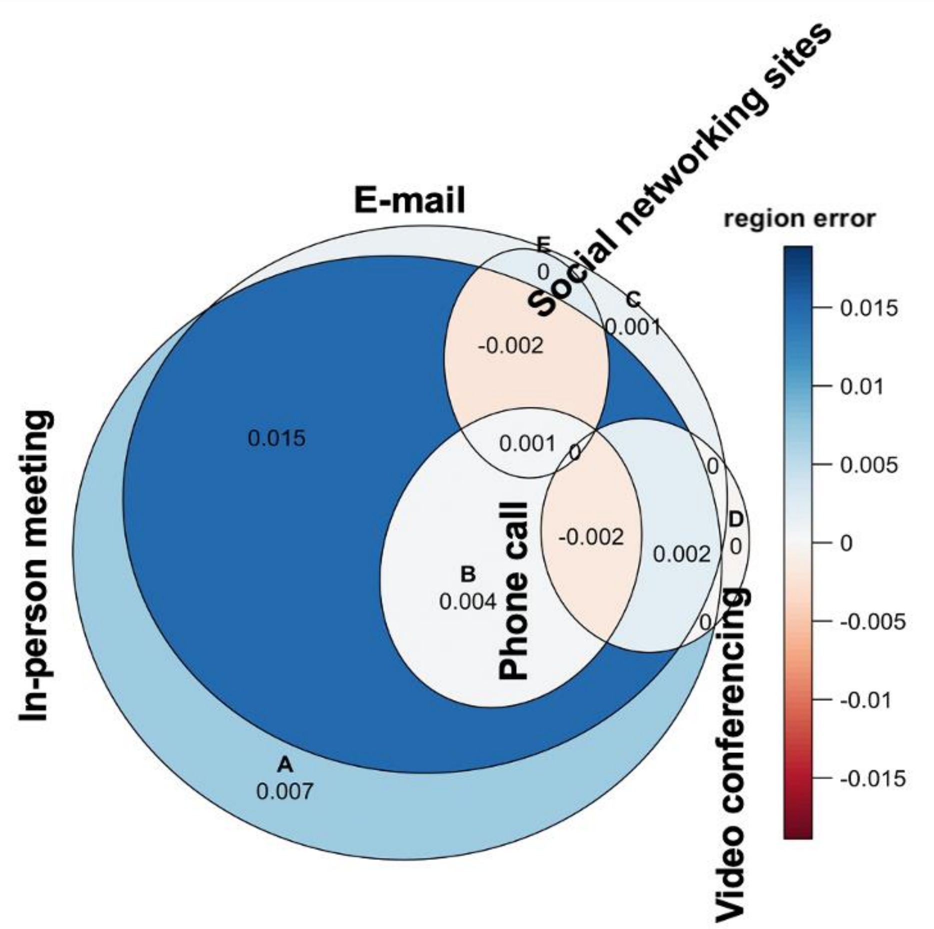

**Figure S1.** Error plot for ‘euler’ * objects shown in Figure 2E, calculated using the *eulerr package* in Rstudio*.* The error plot visually evaluates the fit to the call *euler()* for the optimization performed during plotting of Figure 2E.

* Wickham H. 2016 *ggplot2: elegant graphics for data analysis*.
Larsson J, Gustafsson P. 2018 A Case Study in Fitting Area-Proportional Euler Diagrams with Ellipses Using eulerr. *Proc. Int. Workshop Set Vis. Reason.* **2116**, 84–91.
https://cran.r-project.org/package=eulerr
https://www.rdocumentation.org/packages/eulerr/versions/7.0.0

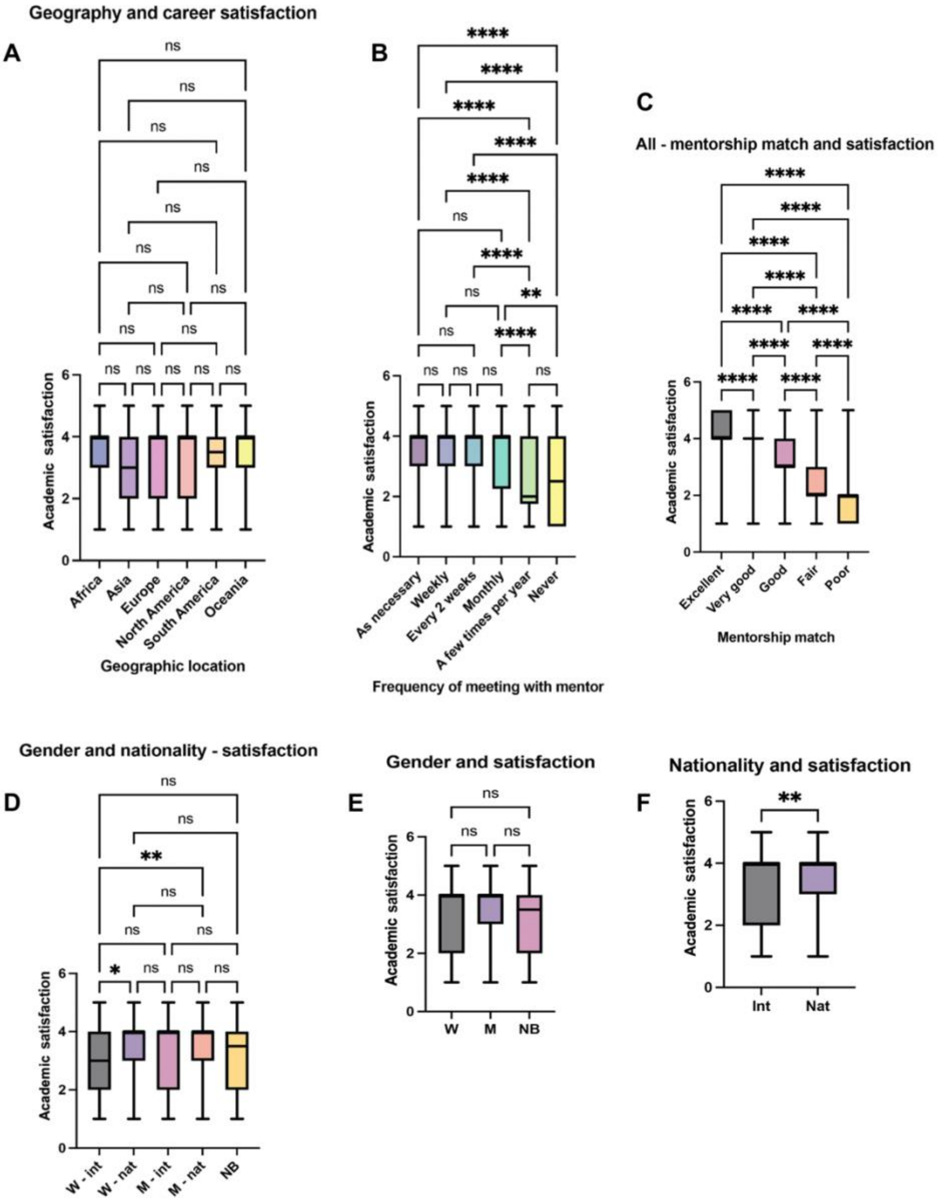

**Figure S2.** Statistical analysis of relationships between respondent academic satisfaction about their research, geographic location (region of research), gender, number of mentors. (A) Academic satisfaction reported by mentees in North America, Europe, or other continents. n.s. = not significant, Ordinary one-way ANOVA (Table S125a), (B) Academic satisfaction reported by mentees by meeting frequency. n.s. = not significant, ** $p<0.01$, **** $p<0.0001$, Ordinary one-way ANOVA (Table S40b), (C) Academic satisfaction reported by mentees by mentorship match. **** $p<0.0001$, Ordinary one-way ANOVA (Table S125c), (D) Academic satisfaction reported by mentees according to their gender (W = women, M = men, NB = non-binary) and nationality (int = international, nat = national). n.s. = not significant, * $p<0.05$, ** $p<0.01$, Ordinary one-way ANOVA (Table S125d), (E) Academic satisfaction reported by mentees according to their gender (W = women, M = men, NB = non-binary). n.s. = not significant, Ordinary one-way ANOVA (Table S125e), (F) Academic satisfaction reported by mentees according to their nationality (int = international, nat = national). ** $p<0.01$, Ordinary one-way ANOVA (Table S125f), For all plots, box and whiskers represent the $5^{th}$-$95^{th}$ percentile of data, with the box representing the $25^{th}$-$75^{th}$ percentile of data. The middle line represents the median.

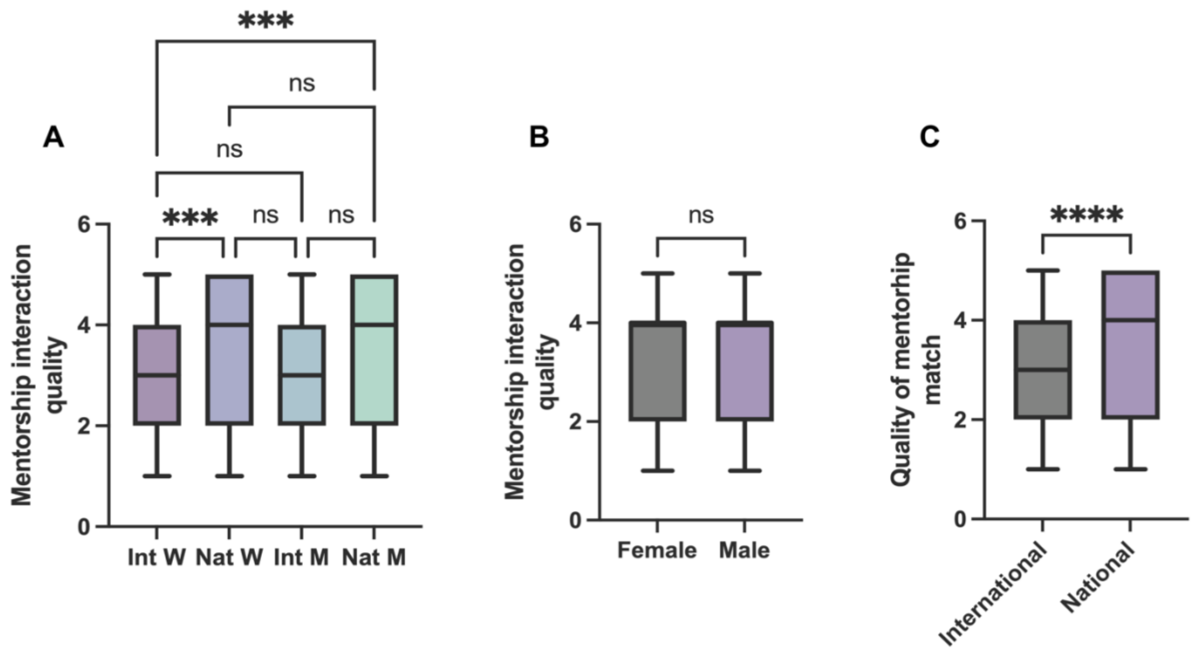

**Figure S3.** Statistical analysis of relationships between respondents' (mentee's) mentor meeting mentee expectations. (A) Quality of mentorship match reported by mentees by gender and nationality. n.s. = not significant, *** p<0.001, Ordinary one-way ANOVA, (Table S126a), (B) Quality of mentorship match reported by mentees by gender (Table S126b), n.s. = not significant, Wilcoxon rank sum test. (C) Quality of mentorship match reported by mentees by nationality. **** p<0.0001, Wilcoxon rank sum test (Table S126c). For all plots, box and whiskers represent the 5$^{th}$-95$^{th}$ percentile of data, with the box representing the 25$^{th}$-75$^{th}$ percentile of data. The middle line represents the median.

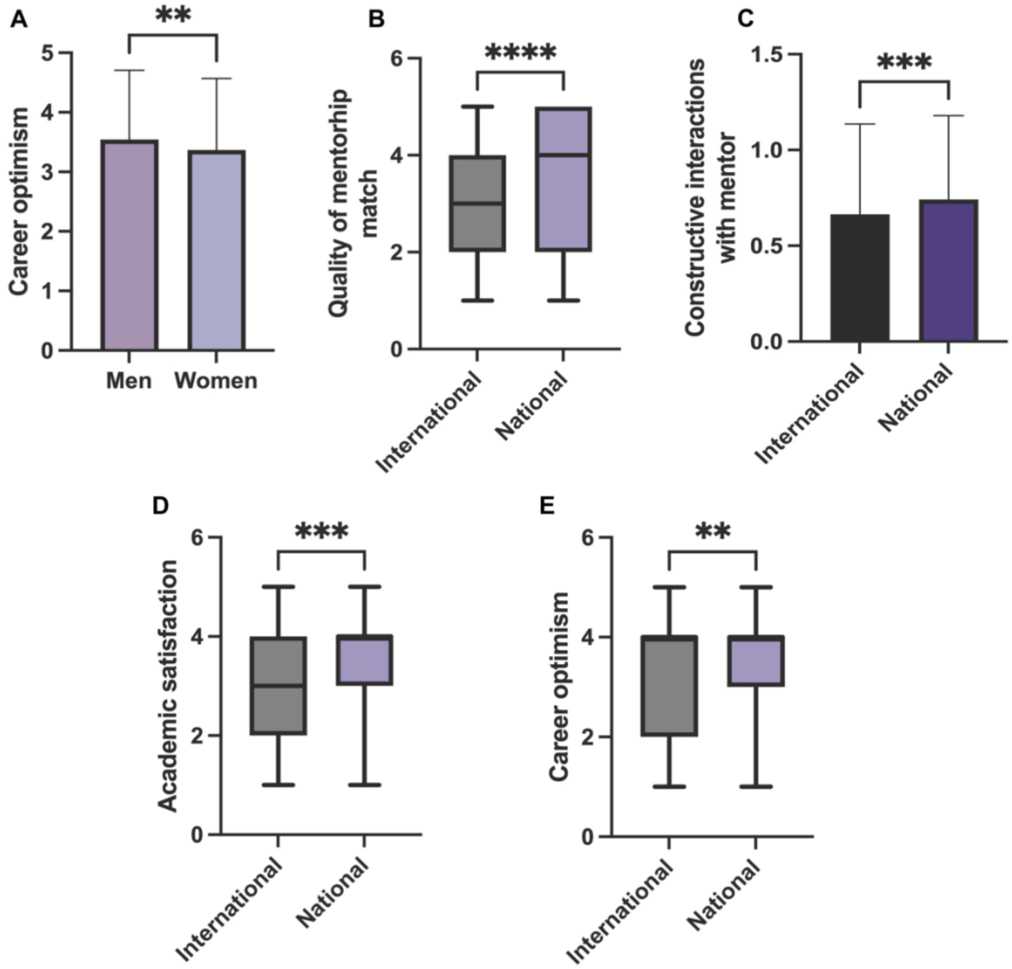

**Figure S4. Statistical analysis for Figure 5-6.** (A) Career optimism as reported by mentees by gender. ** p<0.01, Wilcoxon rank sum test (Table S128a), (B) Quality of mentorship match as reported by nationality. **** p<0.0001, Wilcoxon rank sum test (Table S128b), (C) Constructiveness of interactions with mentor as reported by mentees by nationality. *** p<0.001, Wilcoxon rank sum test (Table S128c), (D) Academic satisfaction match as reported by mentees by nationality. *** p<0.001, Wilcoxon rank sum test (Table S128d), (E) Career optimism as reported by mentees by nationality. ** p<0.01, Wilcoxon rank sum test (Table S128e). For all plots, box and whiskers represent the 5$^{th}$-95$^{th}$ percentile of data, with the box representing the 25$^{th}$-75$^{th}$ percentile of data. The middle line represents the median.

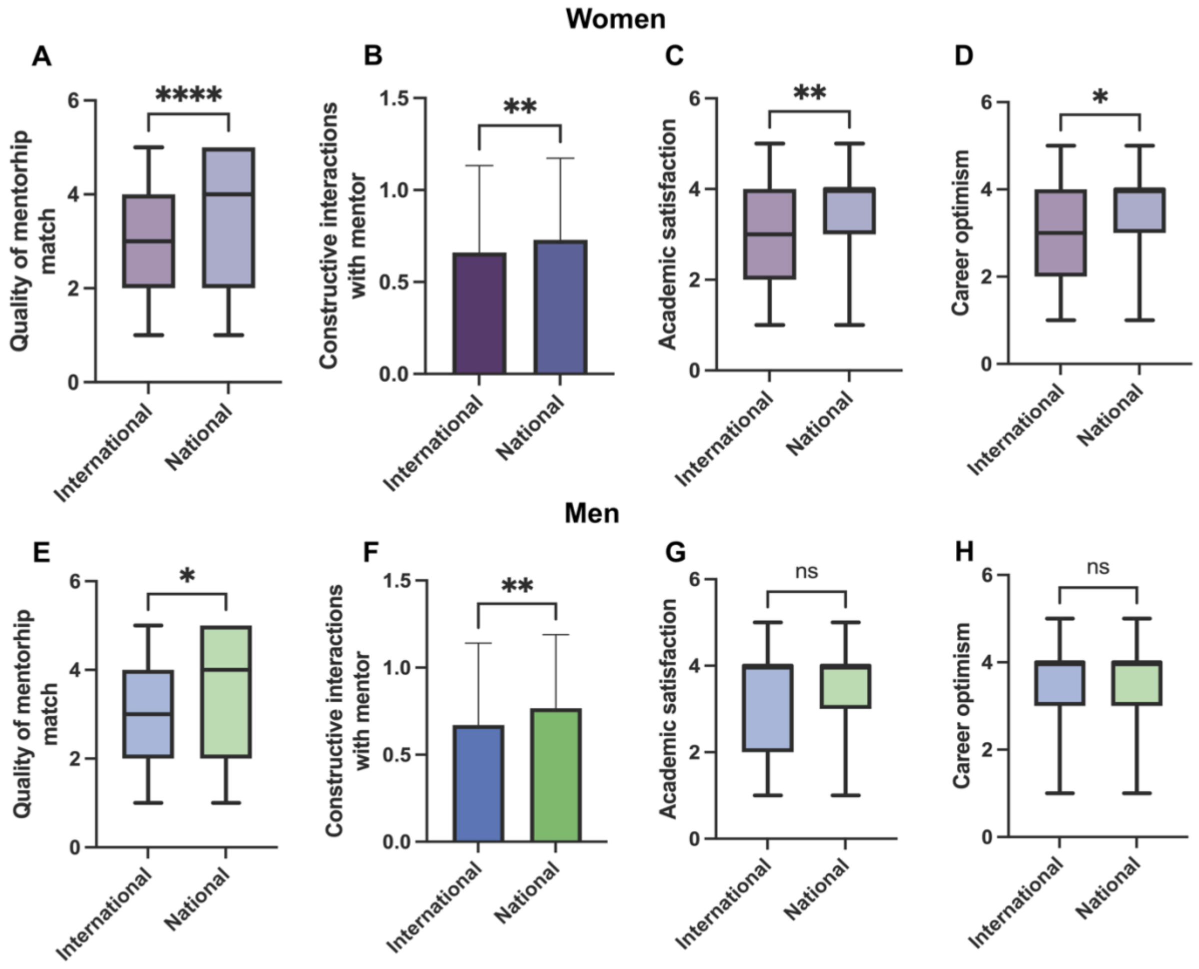

**Figure S5. Statistical analysis for Figure 8-9.** (A) Quality of mentorship match reported by mentee women by nationality. **** $p<0.0001$, Ordinary one-way ANOVA (Table S127a), (B) Constructive interactions as reported by mentee women by nationality. ** $p<0.01$, Wilcoxon rank sum test (Table S127b), (C) Academic satisfaction match reported by mentee women by nationality. ** $p<0.01$, Wilcoxon rank sum test (Table S127c), (D) Career optimism reported by mentee women by nationality. * $p<0.05$, Wilcoxon rank sum test (Table S127d), (E) Quality of mentorship match reported by mentee men by nationality. * $p<0.05$, Ordinary one-way ANOVA (Table S127e), (F) Constructiveness of interactions as reported by mentee men by nationality. ** $p<0.01$, Wilcoxon rank sum test (Table S127f), (G) Academic satisfaction match reported by mentee men by nationality. n.s. = not significant, Wilcoxon rank sum test (Table S127g), (H) Career optimism reported by mentee men by nationality. n.s. = not significant, Wilcoxon rank sum test (Table S127h). For all plots, box and whiskers represent the $5^{th}$-$95^{th}$ percentile of data, with the box representing the $25^{th}$-$75^{th}$ percentile of data. The middle line represents the median.

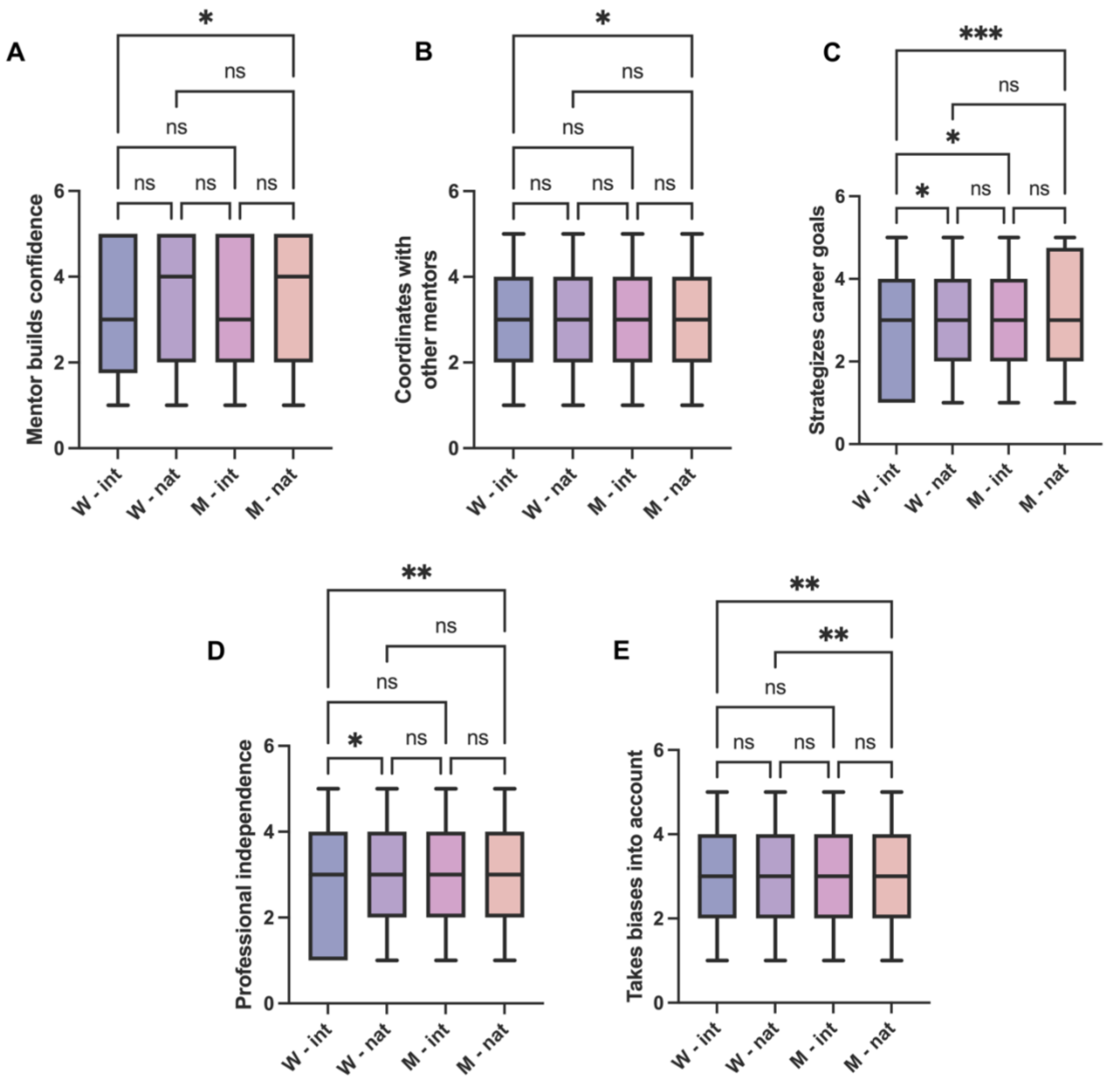

**Figure S6. Statistical analysis for Figure 10.** (A) Assessment of mentor building confidence in mentee as reported by mentees by gender and nationality. n.s. = not significant, * $p<0.05$, Ordinary one-way ANOVA (Table S129a), (B) Assessment of coordination across mentors as reported by mentees by gender and nationality. n.s. = not significant, * $p<0.05$, Ordinary one-way ANOVA (Table S129b), (C) Assessment of mentor's ability to strategize mentee career goals as reported by mentees by gender and nationality. n.s. = not significant, * $p<0.05$, *** $p<0.001$, Ordinary one-way ANOVA (Table S129c), (D) Assessment of mentor promoting mentee professional independence as reported by mentees by gender and nationality. n.s. = not significant, * $p<0.05$, ** $p<0.01$, Ordinary one-way ANOVA (Table S129d), (E) Assessment of mentors taking own biases into account in mentorship interactions as reported by mentees by gender and nationality. n.s. = not significant, ** $p<0.01$, Ordinary one-way ANOVA (Table S129e). For all plots, box and whiskers represent the $5^{th}$-$95^{th}$ percentile of data, with the box representing the $25^{th}$-$75^{th}$ percentile of data. The middle line represents the median.

### Supplementary Tables

**Data availability**

The authors confirm that, for approved reasons, access restrictions apply to the data underlying the findings. Raw data underlying this study cannot be made publicly available in order to safeguard participant anonymity and that of their organizations. Ethical approval for the project was granted on the basis that only aggregated data is provided (as has been provided in the supplementary tables, with appropriate anonymization) as part of this publication.

**Table S1. Trainee Mentee Demographics**
Overview of respondents' country of research.
Countries which had fewer than four respondents in our survey were aggregated.

| Respondent geographic region (Geographic region in which the trainee mentee conducted research in a laboratory) | Number of trainees |
|---|---|
| **by Continent** | |
| **North America** | 51.8% (1093 out of 2112) |
| **Europe** | 28.6% (605 out of 2112) |
| **Asia** | 7.8% (165 out of 2112) |
| **Latin America** | 5.9% (124 out of 2112) |
| **Oceania** | 3.9% (83 out of 2112) |
| **Africa** | 2% (42 out of 2112) |
| **by Country** | |
| **United States** | 47.7% (1008 out of 2112) |
| **United Kingdom** | 7.2% (153 out of 2112) |
| **Germany** | 4.2% (89 out of 2112) |
| **Spain** | 4% (85 out of 2112) |
| **Canada** | 4% (85 out of 2112) |
| **France** | 3.3% (69 out of 2112) |
| **Australia** | 3.2% (67 out of 2112) |
| **Argentina** | 3% (63 out of 2112) |
| **India** | 2.3% (49 out of 2112) |
| **Switzerland** | 2% (43 out of 2112) |
| **The Netherlands** | 1.8% (39 out of 2112) |

| | |
|---|---|
| **South Africa** | 1.6% (33 out of 2112) |
| **Sweden** | 1.2% (25 out of 2112) |
| **Japan** | 1% (22 out of 2112) |
| **Chile** | 1% (21 out of 2112) |
| **New Zealand** | 0.75% (16 out of 2112) |
| **Brazil** | 0.66% (14 out of 2112) |
| **Belgium** | 0.66% (14 out of 2112) |
| **Portugal** | 0.75% (16 out of 2112) |
| **Italy** | 0.56% (12 out of 2112) |
| **Malaysia** | 0.52% (11 out of 2112) |
| **Israel** | 0.43% (9 out of 2112) |
| **Denmark** | 0.52% (11 out of 2112) |
| **Bangladesh** | 0.52% (11 out of 2112) |
| **Albania** | 0.47% (10 out of 2112) |
| **Turkey** | 0.47% (10 out of 2112) |
| **Iran** | 0.37% (8 out of 2112) |
| **Singapore** | 0.37% (8 out of 2112) |
| **Uruguay** | 0.43% (9 out of 2112) |
| **Austria** | 0.37% (8 out of 2112) |
| **Morocco** | 0.33% (7 out of 2112) |
| **Hong Kong** | 0.37% (8 out of 2112) |
| **Ireland** | 0.33% (7 out of 2112) |
| **Venezuela** | 0.3% (6 out of 2112) |
| **Taiwan** | 0.3% (6 out of 2112) |
| **Mexico** | 0.2% (5 out of 2112) |
| **Nepal** | 0.2% (4 out of 2112) |
| **Finland** | 0.2% (4 out of 2112) |
| **Georgia (1) + Ghana (1) + Indonesia (2) + Mali (1) + Nigeria (1) +** | 2.2% (47 out of 2112) |

| | |
|---|---|
| **Thailand (1) + Norway (3) + United Arab Emirates (1) + Croatia (2) + Cambodia (1) + Colombia (1) + Panama (3) + Czech Republic (2) + Hungary (1) + Luxembourg (3) + Kenya (2) + Myanmar (1) + Peru (2) + Philippines (1) + Poland (3) + Russia (2) + Saudi Arabia (1) + Slovenia (2) + Slovakia (1) + South Korea (1) + China (3) + Sudan (1) + Tanzania (3)** | |
| **Did not respond to this survey Question** | 0.1% (2 out of 2114) |

**Table S2. Type of Trainee Mentee Research Institution**

| Theme | count |
|---|---|
| **Academic** | 93.8% (1983 out of 2114) |
| **Industry** | 0.8% (17 out of 2114) |
| **Government** | 4% (85 out of 2114) |
| **Other** | 1.4% (29 out of 2114) |
| **Did not respond to this survey Question** | 0% (0 out of 2114) |

**Table S3. Trainee mentee by their Field of Research**
Respondent self-identified scientific discipline.

| Theme | count |
|---|---|
| **Basic (fundamental) research in Life and Biomedical Sciences** | 52% (1099 out of 2112) |
| **Basic research in Physical Sciences, Engineering research, Theoretical or Mathematical research, Computing and Information Sciences** | 17% (359 out of 2112) |
| **Social Sciences, Humanities, Psychology, Behavioral research, Art and Design, Education research** | 11.2% (237 out of 2112) |
| **Environmental Sciences, Conservation Biology, Field or applied research, Agriculture, Plant Biology** | 6.7% (142 out of 2112) |
| **Clinical Research, Medicine, Public Health** | 9.3% (196 out of 2112) |
| **Translational research (type: T1, T2, etc.)** | 3.7% (79 out of 2112) |
| **Did not respond to this survey Question** | 0.09% (2 out of 2114) |

| **Table S4. Mentee Academic Degree**<br>Type of advanced degree trainee mentee held | |
|---|---|
| **Theme** | **count** |
| **Doctoral Degree (PhD)** | 37.1% (784 out of 2114) |
| **Master's Degree (Sciences, Arts, Engineering)** | 31.5% (666 out of 2114) |
| **Bachelors' Degree (Sciences, Arts, Engineering)** | 25.8% (545 out of 2114) |
| **High School diploma** | 1% (22 out of 2114) |
| **Professional degree (e.g., MD, DDS, RD, PT, PharmD, etc.)** | 1.5% (32 out of 2114) |
| **both PhD and professional degree**<br>**(MD/PhD, MD/MPH, PharmD/MS, etc.)** | 3.1% (65 out of 2114) |
| **Did not respond to this survey Question** | 0% (0 out of 2114) |

| **Table S5. Trainee Mentee Gender**<br>Respondent self-identified gender | |
|---|---|
| **Theme** | **count** |
| **Women (n=1292)** | 61% (1292 out of 2114) |
| **Men (n=802)** | 38% (802 out of 2114) |
| **Non-binary/did not disclose gender (n=20)** | 1% (20 out of 2114) |

| **Table S6. Trainee Mentee Academic Position**<br>Respondent self-identified academic research position | |
|---|---|
| **Theme** | **count** |
| **Graduate researcher**<br>**(researcher in a Master's or Doctoral degree program)** | 57% (1206 out of 2114) |
| **Postdoctoral researcher (has obtained a PhD or MD/PhD)** | 39.7% (839 out of 2114) |
| **Undergraduate researcher**<br>**(researcher in a Bachelor's degree program)** | 1% (20 out of 2114) |
| **Medical student/Health professional/Clinical fellow** | 0.9% (19 out of 2114) |
| **Other**<br>**(researcher, research intern, research scientist (assistant or associate), research officer with a bachelor's or master's degree)** | 1.4% (30 out of 2114) |
| **Did not respond to this survey Question** | 0% (0 out of 2114) |

| **Table S7. Trainee mentee desired independence level**<br>Respondent self-identified desired independence level from mentor | |
|---|---|
| **Theme** | **count** |
| **No guidance or input (level 1)** | 4.3% (90 out of 2099) |
| **Some guidance or input (level 2)** | 36.4% (765 out of 2099) |
| **Moderately managed (level 3)** | 44.3% (930 out of 2099) |
| **Substantially managed (level 4)** | 10.9% (228 out of 2099) |
| **Very heavily managed (level 5)** | 4.1% (86 out of 2099) |
| **Did not respond to this survey Question** | 0.7% (15 out of 2114) |

| **Table S8. Trainee mentee research environment type**<br>Respondent self-identified research environment | |
|---|---|
| **Theme** | **count** |
| **Experimental (wet) lab** | 48.7% (1023 out of 2099) |
| **Computational (dry) lab** | 22.7% (476 out of 2099) |
| **both types**<br>**(hybrid experimental and computational lab)** | 22.5% (472 out of 2099) |
| **Not Applicable** | 6.1% (128 out of 2099) |
| **Did not respond to this survey Question** | 0.7% (15 out of 2114) |

| **Table S9. Do you (trainee mentee) find your interactions with your Mentor constructive?** | |
|---|---|
| **Theme** | **count** |
| **All respondents (n=2091)** | 71.2% said Yes (1488 out of 2091)<br>28.8% said No (603 out of 2091) |
| **Did not respond to this survey Question (n=19)** | 0.9% (19 out of 2114) |

| **Table S10. What format do you (trainee mentee) use to communicate with your Mentor?** | |
|---|---|
| **Theme** | **format of Mentor-Mentee communication** |
| **Total**<br>**(n=2102)** | **Use of a single approach Only:(n=511 or 24.3%)**<br>19.1% said Face-to-Face in-person (400 out of 2102)<br>0.05% said Telephone (1 out of 2102)<br>4.1% said E-mail (86 out of 2102)<br>0.8% said Web/Video conferencing (e.g., Skype, Zoom) (17 out of 2102)<br>0.2% said Social networking sites (5 out of 2102) |

| | |
|---|---|
| | **Use More than one approach:(n=1593 or 75.7%)**<br>0.3% said Telephone+E-mail (6 out of 2102)<br>1.0% said E-mail+Social networking sites (e.g., Slack chat or instant messaging on WatsApp) (21 out of 2102)<br>0.2% said Telephone+E-mail+Social networking sites (e.g., Slack chat or instant messaging on WatsApp) (4 out of 2102)<br>41.3% said Face-to-Face+E-mail (870 out of 2102)<br>0.5% said Face-to-Face+Telephone (10 out of 2102)<br>0.4% said E-mail+Web/Video conferencing (e.g., Skype, Zoom) (9 out of 2102)<br>0.3% said Face-to-Face+Web/Video conferencing (e.g., Skype, Zoom) (6 out of 2102)<br>5.7% said Face-to-Face+E-mail+Social networking sites (e.g., Slack chat or instant messaging on WatsApp) (119 out of 2102)<br>11.5% said Face-to-Face+Telephone+E-mail (242 out of 2102)<br>0.2% said Telephone+E-mail+Web/Video conferencing (e.g., Skype, Zoom) (4 out of 2102)<br>5.2% said Face-to-Face+E-mail+Web/Video conferencing (e.g., Skype, Zoom) (110 out of 2102)<br>0.1% said Face-to-Face+Web/Video conferencing (e.g., Skype, Zoom)+Social networking sites (e.g., Slack chat or instant messaging on WatsApp)(3 out of 2102)<br>1.7% said Face-to-Face+E-mail+Web/Video conferencing (e.g., Skype, Zoom)+Social networking sites (e.g., Slack chat or instant messaging on WatsApp)(36 out of 2102)<br>1.8% said Face-to-Face+E-mail+Telephone+Social networking sites (e.g., Slack chat or instant messaging on WatsApp)(37 out of 2102)<br>4.3% said Face-to-Face+E-mail+Telephone+Web/Video conferencing (e.g., Skype, Zoom) (91 out of 2102)<br>0.1% said E-mail+Telephone+Web/Video conferencing (e.g., Skype, Zoom)+Social networking sites e.g., Slack chat or instant messaging on WatsApp) (2 out of 2102)<br>0.1% said Face-to-Face+Telephone+Web/Video conferencing (e.g., Skype, Zoom)+Social networking sites e.g., Slack chat or instant messaging on WatsApp) (2 out of 2102)<br>1.0% said Face-to-Face+E-mail+Telephone+Web/Video conferencing (e.g., Skype, Zoom)+Social networking sites e.g., Slack chat or instant messaging on WatsApp) (21 out of 2102) |
| **Did not respond to this survey Question (n=10)** | 0.05% (10 out of 2114) |

| Table S11. Trainee mentee meeting frequency with mentor | |
|---|---|
| **Theme** | **count** |
| **Scheduled as necessary** | 30.5% (642 out of 2100) |
| **Weekly** | 34.4% (723 out of 2100) |
| **Every 2 weeks** | 16.9% (355 out of 2100) |
| **Monthly** | 8% (169 out of 2100) |
| **Few times a year** | 7.7% (162 out of 2100) |

| | |
|---|---|
| **Never** | 2.3% (49 out of 2100) |
| **Did not respond to this survey Question** | 0.5% (10 out of 2110) |

| **Table S12. Method of original mentee-mentor matching**<br>Respondent self-identified mentorship initiation mode | |
|---|---|
| **Theme** | **count** |
| **Total (n=2088)** | 93% said Voluntary (I chose Mentor) (1942 out of 2088)<br>7% said I was assigned to my Mentor (146 out of 2088) |
| **Did not respond to this survey Question (n=24)** | 1.1% (24 out of 2112) |

| **Table S13. Number of mentors that trainee mentee had**<br>All percentages are calculated out of the total number of respondents to this particular survey question, not the total number of overall survey respondents (n=2114). | |
|---|---|
| **Theme** | **count** |
| **Total (n=1990)** | 0.4% had 0 mentors (6 out of 1990)<br>53.4% had 1 mentor (1061 out of 1990)<br>29.4% had 2 mentors (584 out of 1990)<br>11.8% had 3 mentors (238 out of 1990)<br>2.8% had 4 mentors (55 out of 1990)<br>%1.3 had 5 mentors (25 out of 1990)<br>0.9% had >5 mentors (17 out of 1990)<br>Average= 1.7<br>Median= 1 |

| **Table S13a. Number of mentors Trainee had by gender**<br>All percentages are calculated out of the total number of respondents to this question n=2011. | |
|---|---|
| **Theme** | **count** |
| **Women (n=1226)** | 0.3% had 0 mentors (3 out of 1226)<br>52.4% had 1 mentors (643 out of 1226)<br>28.2% had 2 mentors (346 out of 1226)<br>13.1% had 3 mentors (161 out of 1226)<br>3.3% had 4 mentors (41 out of 1226)<br>1.5% had 5 mentors (18 out of 1226)<br>1.1% had >5 mentors (14 out of 1226)<br>Average=1.8;<br>Median=1 |
| **Men (n=736)** | 0.5% had 0 mentors (3 out of 736)<br>55.1% had 1 mentors (406 out of 736)<br>31.2% had 2 mentors (230 out of 736)<br>9.8% had 3 mentors (72 out of 736)<br>1.8% had 4 mentors (13 out of 736)<br>0.9% had 5 mentors (7 out of 736)<br>0.7% had >5 mentors (5 out of 736) |

| | |
|---|---|
| | Average=1.6;<br>Median=1 |
| **Non-binary or preferred not to disclose gender (n=22)** | 0% had 0 mentors (0 out of 22)<br>40.9% had 1 mentors (9 out of 22)<br>31.8% had 2 mentors (7 out of 22)<br>9.1% had 3 mentors (2 out of 22)<br>9.1% had 4 mentors (2 out of 22)<br>9.1% had 5 mentors (2 out of 22)<br>0% had >5 mentors (0 out of 22)<br>Average=2.1;<br>Median=2 |
| **Did not respond to this survey Question** | 0.1% (3 out of 2114) |

**Table S14. Trainee Age distribution**
All percentages are calculated out of the total number of respondents to this question.

| Theme | count |
|---|---|
| **Total (n=2112)** | 16.3% said 21-25 (345 out of 2112)<br>38.1% said 25-30 (805 out of 2112)<br>32.1% said 30-35 (677 out of 2112)<br>10.3% said 35-40 (218 out of 2112)<br>3.2% said 40-45 (67 out of 2112) |
| **Preferred not to say (n=2)** | 0.09% (2 out of 2114) |

**Table S15. Number of years trainee mentee has worked with their mentor**
All percentages are calculated out of the total number of respondents to this question n=2095.

| Theme | count |
|---|---|
| **Total (n=2095)** | 14.9% Less than one year (313 out of 2095)<br>29.4% 1-2 years (616 out of 2095)<br>34% 3-4 years (711 out of 2095)<br>21.7% 5 or more years (455 out of 2095) |
| **Women (n=1294)** | 15.4% Less than one year (200 out of 1294)<br>28% 1-2 years (362 out of 1294)<br>35.2% 3-4 years (455 out of 1294)<br>21.4% 5 or more years (277 out of 1294) |
| **Men (n=794)** | 14% Less than one year (111 out of 794)<br>31.2% 1-2 years (248 out of 794)<br>32.9% 3-4 years (261 out of 794)<br>21.9% 5 or more years (174 out of 794) |
| **Non-binary or did not disclose gender (n=14)** | 14.3% Less than one year (2 out of 14)<br>28.6% 1-2 years (4 out of 14)<br>35.7% 3-4 years (5 out of 14)<br>21.4% 5 or more years (3 out of 14) |
| **Did not respond to this survey Question** | 0.9% (19 out of 2114) |

| Table S16. Respondents (trainee mentees) rating of the quality of the match between mentee and mentor, responses by gender. | |
|---|---|
| **Theme** | **count** |
| **Total (n=2105)** | 23.7% said Excellent (499 out of 2105)<br>26.8% said Very Good (564 out of 2105)<br>19.8% said Good (417 out of 2105)<br>14.9% said Fair (313 out of 2105)<br>14.8% said Poor (308 out of 2105) |

| Table S17. Trainee mentees rating of the quality of the match between mentee and mentor. analysis by continent | |
|---|---|
| **Theme** | **count** |
| **Total (n=2105)** | 23.7% said Excellent (499 out of 2105)<br>26.8% said Very Good (565 out of 2105)<br>19.8% said Good (417 out of 2105)<br>14.9% said Fair (313 out of 2105)<br>14.8% said Poor (311 out of 2105) |
| **North America (n=1091)** | 27.4% said Excellent (299 out of 1091)<br>26.5% said Very Good (289 out of 1091)<br>18.3% said Good (200 out of 1091)<br>13.7% said Fair (150 out of 1091)<br>14% said Poor (153 out of 1091) |
| **Europe (n=591)** | 17.3% said Excellent (102 out of 591)<br>27.7% said Very Good (164 out of 591)<br>21.7% said Good (128 out of 591)<br>15.1% said Fair (89 out of 591)<br>18.3% said Poor (108 out of 591) |
| **Latin America+Oceania+Africa+Asia (n=420)** | 23.1% said Excellent (97 out of 420)<br>26.7% said Very Good (112 out of 420)<br>21.2% said Good (89 out of 420)<br>17.4% said Fair (73 out of 420)<br>11.7% said Poor (49 out of 420) |
| **Did not respond to this survey Question (n=9)** | 0.4% (9 out of 2114) |

| Table S18. Do you (trainee mentee) find your interactions with your Mentor constructive? analysis by continent | |
|---|---|
| **Theme** | **count** |
| **All respondents (n=2095)** | 71.1% said Yes (1490 out of 2095)<br>28.9% said No (605 out of 2095) |
| **North America (n=1087)** | 72.9% said Yes (792 out of 1087)<br>27.1% said No (295 out of 1087) |
| **Europe (n=586)** | 66.7% said Yes (391 out of 586)<br>33.3% said No (195 out of 586) |

| | |
|---|---|
| **Latin America+Asia+Africa+Oceania (n=419)** | 73% said Yes (306 out of 419)<br>27% said No (113 out of 419) |
| **Did not respond to this survey Question (n=19)** | 0.9% (19 out of 2114) |

| **Table S19. Do you feel happy and satisfied with your current research and position as a trainee mentee, analysis by continent** | |
|---|---|
| **Theme** | **count** |
| **Total (n=1899)** | 64.5% Yes (1225 out of 1899)<br>35.5% No (674 out of 1899) |
| **North America (n=972)** | 66% Yes (642 out of 972)<br>34% No (330 out of 972) |
| **Europe (n=525)** | 61.5% Yes (323 out of 525)<br>38.5% No (202 out of 525) |
| **Latin America+Oceania+Africa+Asia (n=399)** | 64.9% Yes (259 out of 399)<br>35.1% No (140 out of 399) |
| **Did not respond to this survey Question (n=215)** | 10.1% (215 out of 2114) |

| **Table S19a. Please rate your feeling of happiness and satisfaction with your current research and position as a trainee mentee, analysis by continent.** on a Likert Scale (1 lowest-5 highest). | |
|---|---|
| **Theme** | **count** |
| **Total (n=1705)** | 12% rated 1 (lowest score) (205 out of 1705)<br>17.3% rated 2 (295 out of 1705)<br>21.7% rated 3 (370 out of 1705)<br>42.4% rated 4 (723 out of 1705)<br>18.6% rated 5 (highest score) (317 out of 1705) |
| **North America (n=980)** | 11.2% rated 1 (lowest score) (110 out of 980)<br>14.4% rated 2 (141 out of 980)<br>17.8% rated 3 (174 out of 980)<br>37.1% rated 4 (364 out of 980)<br>19.5% rated 5 (highest score) (191 out of 980) |
| **Europe (n=537)** | 11% rated 1 (lowest score) (59 out of 537)<br>17.9% rated 2 (96 out of 537)<br>18.6% rated 3 (100 out of 537)<br>40.2% rated 4 (216 out of 537)<br>12.3% rated 5 (highest score) (66 out of 537) |
| **Latin America+Oceania+Africa+Asia (n=390)** | 9.2% rated 1 (lowest score) (36 out of 390)<br>14.4% rated 2 (56 out of 390)<br>24.6% rated 3 (96 out of 390)<br>36.4% rated 4 (142 out of 390)<br>15.4% rated 5 (highest score) (60 out of 390) |
| **Did not respond to this survey Question (n=3)** | 0.2% (3 out of 1705) |

| **Table S20. Do you (trainee mentee) feel optimistic about your future career?**<br>**analysis by continent** | |
|---|---|
| **Theme** | **count** |
| **Total (n=1901)** | 68.6% Yes (1304 out of 1901)<br>31.4% No (597 out of 1901) |
| **North America (n=975)** | 72.2% Yes (704 out of 975)<br>27.8% No (271 out of 975) |
| **Europe (n=529)** | 62.4% Yes (330 out of 529)<br>37.6% No (199 out of 529) |
| **Latin America+Oceania+Africa+Asia (n=394)** | 68.3% Yes (269 out of 394)<br>31.7% No (125 out of 394) |
| **Did not respond to this survey Question (n=213)** | 10% (213 out of 2114) |

| **Table S20a. Please rate your optimism about the future as a trainee mentee.**<br>on a Likert Scale (1 lowest-5 highest), analysis by continent | |
|---|---|
| **Theme** | **count** |
| **Total (n=1911)** | 8.5% rated 1 (lowest score) (163 out of 1911)<br>14.2% rated 2 (271 out of 1911)<br>21.2% rated 3 (406 out of 1911)<br>37.4% rated 4 (714 out of 1911)<br>18.7% rated 5 (highest score) (347 out of 1911) |
| **North America (n=981)** | 8.2% rated 1 (lowest score) (80 out of 981)<br>13.6% rated 2 (133 out of 981)<br>19.4% rated 3 (190 out of 981)<br>38.1% rated 4 (374 out of 981)<br>20.8% rated 5 (highest score) (204 out of 981) |
| **Europe (n=539)** | 8.5% rated 1 (lowest score) (46 out of 539)<br>16.7% rated 2 (90 out of 539)<br>22.8% rated 3 (123 out of 539)<br>37.7% rated 4 (203 out of 539)<br>14.3% rated 5 (highest score) (77 out of 539) |
| **Latin America+Africa<br>+Asia+Oceania (n=388)** | 9.5% rated 1 (lowest score) (37 out of 388)<br>11.9% rated 2 (46 out of 388)<br>23.7% rated 3 (92 out of 388)<br>35.3% rated 4 (137 out of 388)<br>19.6% rated 5 (highest score) (76 out of 388) |
| **Did not respond to this survey Question (n=3)** | 0.15% (3 out of 1911) |

| Table S21. Respondents (trainee mentees) rating of the quality of the match between mentee and mentor, responses by citizenship in the country of research status | |
|---|---|
| **Theme** | **count** |
| **Total (n=2105)** | 23.7% said Excellent (499 out of 2105)<br>26.8% said Very Good (565 out of 2105)<br>19.8% said Good (417 out of 2105)<br>14.9% said Fair (313 out of 2105)<br>14.8% said Poor (311 out of 2105) |
| **International (non-citizen) Mentee (n=848)** | 18.6% said Excellent (158 out of 848)<br>23.7% said Very Good (201 out of 848)<br>25.7% said Good (218 out of 848)<br>14.5% said Fair (123 out of 848)<br>17.5% said Poor (148 out of 848) |
| **National (citizen) Mentee (n=1302)** | 26,2% said Excellent (341 out of 1302)<br>28% said Very Good (364 out of 1302)<br>18.7% said Good (244 out of 1302)<br>14.6% said Fair (190 out of 1302)<br>12.5% said Poor (163 out of 1302) |
| **Did not respond to this survey Question (n=9)** | 0.4% (9 out of 2114) |

| Table S22. Do you (trainee mentee) find your interactions with your Mentor constructive? Analysis by citizenship status | |
|---|---|
| **Theme** | **count** |
| **All respondents (n=2095)** | 71.1% said Yes (1490 out of 2095)<br>28.9% said No (605 out of 2095) |
| **International (non-citizen) Mentee (n=799)** | 66.3% said Yes (530 out of 799)<br>33.7% said No (269 out of 799) |
| **National (citizen) Mentee (n=1296)** | 74.1% said Yes (960 out of 1296)<br>25.9% said No (336 out of 1296) |
| **Did not respond to this survey Question (n=19)** | 0.9% (19 out of 2114) |

| Table S23. Do you feel happy and satisfied with your current research and position as a trainee mentee by citizenship in their country of research status. | |
|---|---|
| **Theme** | **count** |
| **Total (n=1899)** | 64.5% Yes (1225 out of 1899)<br>35.5% No (674 out of 1899) |
| **International (non-citizen) Mentee (n=708)** | 61.2% Yes (433 out of 708)<br>38.8% No (275 out of 708) |
| **National (citizen) Mentee (n=1191)** | 66.5% Yes (792 out of 1191)<br>33.5% No (399 out of 1191) |
| **Did not respond to this survey Question (n=215)** | 10.1% (215 out of 2114) |

| Table S24. Do you feel optimistic about your future career?<br>Responses by citizenship in country of research status | |
|---|---|
| **Theme** | **count** |
| **Total (n=1901)** | 68.6% Yes (1304 out of 1901)<br>31.4% No (597 out of 1901) |
| **International (non-citizen) Mentee (n=708)** | 63.7% Yes (451 out of 708)<br>36.3% No (257 out of 708) |
| **National (citizen) Mentee (n=1193)** | 71.5% Yes (853 out of 1193)<br>28.5% No (340 out of 1193) |
| **Did not respond to this survey Question (n=213)** | 10% (213 out of 2114) |

| Table S25. Respondents (trainee mentees) rating of the quality of the match between mentee and mentor, responses by gender. | |
|---|---|
| **Theme** | **count** |
| **Total (n=2105)** | 23.7% said Excellent (499 out of 2105)<br>26.8% said Very Good (565 out of 2105)<br>19.8% said Good (417 out of 2105)<br>14.9% said Fair (313 out of 2105)<br>14.8% said Poor (311 out of 2105) |
| **Women Mentee (n=1281)** | 23.7% said Excellent (304 out of 1281)<br>30.3% said Very Good (388 out of 1281)<br>16.1% said Good (206 out of 1281)<br>14.9% said Fair (191 out of 1281)<br>15% said Poor (192 out of 1281) |
| **Men Mentee (n=798)** | 23.5% said Excellent (188 out of 798)<br>30.6% said Very Good (244 out of 798)<br>17.2% said Good (137 out of 798)<br>14.4% said Fair (115 out of 798)<br>14.3% said Poor (114 out of 798) |
| **Non-binary or did not disclose gender (n=23)** | 30.4% said Excellent (7 out of 23)<br>13% said Very Good (3 out of 23)<br>17.4% said Good (4 out of 23)<br>21.7% said Fair (5 out of 23)<br>17.4% said Poor (4 out of 23) |
| **Did not respond to this survey Question (n=9)** | 0.4% (9 out of 2114) |

| Table S26. Do you find your interactions with your Mentor constructive? by gender | |
|---|---|
| **Theme** | **count** |
| **All respondents (n=2095)** | 71.1% said Yes (1490 out of 2095)<br>28.9% said No (605 out of 2095) |

| | |
|---|---|
| **Women (n=1272)** | 70.6% said Yes (898 out of 1272)<br>29.4% said No (374 out of 1272) |
| **Men (n=797)** | 72.4% said Yes (577 out of 797)<br>27.6% said No (220 out of 797) |
| **Non-binary or did not disclose gender (n=23)** | 65.2% said Yes (15 out of 23)<br>34.8% said No (8 out of 23) |
| **Did not respond to this survey Question (n=19)** | 0.9% (19 out of 2114) |

**Table S27. Do you feel happy and satisfied with your current research and position as a trainee mentee.**

| **Theme** | **count** |
|---|---|
| **Total (n=1899)** | 64.5% Yes (1225 out of 1899)<br>35.5% No (674 out of 1899) |
| **Women Mentee (n=1152)** | 63.2% Yes (728 out of 1152)<br>36.8% No (424 out of 1152) |
| **Men Mentee (n=721)** | 67% Yes (483 out of 721)<br>33% No (238 out of 721) |
| **Non-binary or did not disclose gender (n=23)** | 60.9% Yes (14 out of 23)<br>39.1% No (9 out of 23) |
| **Did not respond to this survey Question (n=215)** | 10.1% (215 out of 2114) |

**Table S27a. Please rate your feeling of happiness and satisfaction with your current research and position as a trainee mentee, by gender** on a Likert Scale (1 lowest-5 highest)

| **Theme** | **count** |
|---|---|
| **Total (n=1910)** | 10.7% rated 1 (lowest score) (205 out of 1910)<br>15.4% rated 2 (295 out of 1910)<br>19.4% rated 3 (370 out of 1910)<br>37.9% rated 4 (723 out of 1910)<br>16.6% rated 5 (highest score) (317 out of 1910) |
| **Women Mentee (n=1160)** | 10.8% rated 1 (lowest score) (125 out of 1160)<br>16.3% rated 2 (189 out of 1160)<br>20.8% rated 3 (241 out of 1160)<br>36.3% rated 4 (421 out of 1160)<br>15.9% rated 5 (highest score) (184 out of 1160) |
| **Men Mentee (n=724)** | 10.4% rated 1 (lowest score) (75 out of 724)<br>13.8% rated 2 (100 out of 724)<br>17.5% rated 3 (127 out of 724)<br>40.3% rated 4 (292 out of 724)<br>18% rated 5 (highest score) (130 out of 724) |
| **Non-binary or did not disclose gender (n=23)** | 13% rated 1 (lowest score) (3 out of 23)<br>21.7% rated 2 (5 out of 23)<br>8.7% rated 3 (2 out of 23) |

| | |
|---|---|
| | 43.5% rated 4 (10 out of 23)<br>13% rated 5 (highest score) (3 out of 23) |
| **Did not respond to this survey Question (n=204)** | 9.5% (204 out of 2114) |

| **Table S28. Do you (trainee mentee) feel optimistic about your future career?** | |
|---|---|
| **Theme** | **count** |
| **Total (n=1901)** | 68.6% Yes (1304 out of 1901)<br>31.4% No (597 out of 1901) |
| **Women Mentee (n=1152)** | 66.6% Yes (767 out of 1152)<br>33.4% No (385 out of 1152) |
| **Men Mentee (n=723)** | 71.6% Yes (518 out of 723)<br>28.4% No (205 out of 723) |
| **Non-binary or did not disclose gender (n=23)** | 78.3% Yes (18 out of 23)<br>21.7% No (5 out of 23) |
| **Did not respond to this survey Question (n=213)** | 10% (213 out of 2114) |

| **Table S28a Please rate your optimism about the future as a trainee mentee, analysis by gender.** on a Likert Scale (1 lowest-5 highest) | |
|---|---|
| **Theme** | **count** |
| **Total (n=1911)** | 8.5% rated 1 (lowest score) (163 out of 1911)<br>14.2% rated 2 (271 out of 1911)<br>21.2% rated 3 (406 out of 1911)<br>37.4% rated 4 (714 out of 1911)<br>18.7% rated 5 (highest score) (357 out of 1911) |
| **Women Mentee (n=1160)** | 9.3% rated 1 (lowest score) (108 out of 1160)<br>15% rated 2 (174 out of 1160)<br>22.2% rated 3 (257 out of 1160)<br>36.4% rated 4 (422 out of 1160)<br>17.2% rated 5 (highest score) (199 out of 1160) |
| **Men Mentee (n=725)** | 6.9% rated 1 (lowest score) (50 out of 725)<br>13.1% rated 2 (95 out of 725)<br>19.9% rated 3 (144 out of 725)<br>39% rated 4 (283 out of 725)<br>21.1% rated 5 (highest score) (153 out of 725) |
| **Non-binary or did not disclose gender (n=23)** | 13% rated 1 (lowest score) (3 out of 23)<br>8.7% rated 2 (2 out of 23)<br>17.4% rated 3 (4 out of 23)<br>39.1% rated 4 (9 out of 23)<br>21.7% rated 5 (highest score) (5 out of 23) |
| **Did not respond to this survey Question** | 9.6% (203 out of 2114) |

| Table S29. Has your (trainee mentee) Mentor discussed non-academic careers with you? by gender | |
|---|---|
| **Theme** | **count** |
| **All respondents (n=1915)** | 39.9% said Yes (764 out of 1915)<br>60.1% said No (1151 out of 1915) |
| **Women (n=1162)** | 38.5% said Yes (447 out of 1162)<br>61.5% said No (715 out of 1162) |
| **Men (n=726)** | 41.7% said Yes (303 out of 726)<br>58.3% said No (423 out of 726) |
| **Non-binary or did not disclose gender (n=24)** | 54.2% said Yes (13 out of 24)<br>45.8% said No (11 out of 24) |
| **Did not respond to this survey Question** | 9.4% (199 out of 2114) |

| Table S30. Does your (trainee mentee) mentor work in the same institution as you? | |
|---|---|
| **Theme** | **count** |
| **All respondents (n=2106)** | 92.3% said Yes (1943 out of 2106)<br>7.7% said No (163 out of 2106) |
| **Did not respond to this survey Question** | 0.4% (8 out of 2106) |

| Table S31. Does your (trainee mentee) mentor work in the same lab/group physical space as you? | |
|---|---|
| **Theme** | **count** |
| **All respondents (n=2103)** | 77% said Yes (1617 out of 2103)<br>23% said No (486 out of 2103) |
| **Did not respond to this survey Question** | 0.5% (11 out of 2114) |

| **Table S32. Mentor gender same as trainee mentee or not.** Responses included “Yes” or “No” only | |
|---|---|
| **Theme** | **count** |
| **Total (n=2104)** | 46% said Yes (968 out of 2104)<br>54% said No (1136 out of 2104) |
| **Women Mentee(n=1281)** | 30.8% said Yes (395 out of 1281)<br>69.2% said No (886 out of 1281) |
| **Men Mentee(n=797)** | 70.5% said Yes (562 out of 797)<br>29.5% said No (235 out of 797) |

| | |
|---|---|
| **Non-binary or did not disclose gender (n=23)** | 43.5% said Yes (10 out of 23)<br>56.5% said No (13 out of 23) |
| **Did not respond to this survey Question** | 0.5% (10 out of 2114) |

| **Table S33. Do you (trainee mentee) maintain contact with your former mentors? by gender** | |
|---|---|
| **Theme** | **count** |
| **Total (n=2086)** | 76% said Yes (1585 out of 2086)<br>24% said No (501 out of 2086) |
| **Women Mentee(n=1266)** | 75.9% said Yes (961 out of 1266)<br>24.1% said No (305 out of 1266) |
| **Men Mentee(n=794)** | 75.8% said Yes (602 out of 794)<br>24.2% said No (192 out of 794) |
| **Non-binary or did not disclose gender (n=23)** | 87% said Yes (20 out of 23)<br>13% said No (3 out of 23) |
| **Did not respond to this survey Question** | 1.3% (28 out of 2114) |

| **Table S34. Trainee mentee citizenship status in country of research by gender** | |
|---|---|
| **Theme** | **count** |
| **Total (n=2111)** | 61.8% said Yes (1304 out of 2111)<br>38.2% said No (807 out of 2111) |
| **Women (n=1282)** | 65.4% said Yes (839 out of 1282)<br>34.6% said No (443 out of 1282) |
| **Men (n=802)** | 55.6% said Yes (446 out of 802)<br>44.4% said No (356 out of 802) |
| **Non-binary or did not disclose (n=24)** | 75% said Yes (18 out of 24)<br>25% said No (6 out of 24) |
| **Did not respond to this survey Question (n=3)** | 0.1% (3 out of 2114) |

| **Table S35. Trainee mentee citizenship status in country of research by continent** | |
|---|---|
| **Theme** | **count** |
| **Total (n=2111)** | 61.8% said Yes (1304 out of 2111)<br>38.2% said No (807 out of 2111) |
| **North America (n=1091)** | 67.5% said Yes (736 out of 1091)<br>32.5% said No (355 out of 1091) |
| **Europe (n=600)** | 41.5% said Yes (249 out of 600) |

| | |
|---|---|
| | 58.5% said No (351 out of 600) |
| **Africa+Latin America+Asia+Oceania (n=417)** | 76.3% said Yes (318 out of 417)<br>23.7% said No (99 out of 417) |
| **Did not respond to this survey Question (n=3)** | 0.1% (3 out of 2114) |

| **Table S36. Characterization of trainee mentees by continent, gender and citizenship status** | |
|---|---|
| **Theme** | **count** |
| **North America<br>(n=1158)** | 37.6% Men (435 out of 1158)<br>62.4% Women (723 out of 1158) |
| **North America<br>Men (n=435)** | 59.5% National (259 out of 435)<br>40.5% International (176 out of 435) |
| **North America<br>Women (n=722)** | 69.4% National (501 out of 722)<br>30.6% International (221 out of 722) |
| **Europe<br>(n=594)** | 38% Men (226 out of 594)<br>62% Women (368 out of 594) |
| **Europe<br>Men (n=237)** | 35.4% National (84 out of 237)<br>64.6% International (153 out of 237) |
| **Europe<br>Women (n=368)** | 44.8% National (165 out of 368)<br>55.2% International (203 out of 368) |
| **Latin America+Africa<br>+Asia+Oceania (n=413)** | 41.4% Men (171 out of 413)<br>58.6% Women (242 out of 413) |
| **Latin America+Africa<br>+Asia+Oceania<br>Men (n=177)** | 68.4% National (121 out of 177)<br>31.6% International (56 out of 177) |
| **Latin America+Africa<br>+Asia+Oceania<br>Women (n=242)** | 81.4% National (197 out of 242)<br>18.6% International (45 out of 242) |
| **Did not respond to this survey Question (n=120)** | 5.7% (120 out of 2114) |

| **Table S37. Trainee mentees rating of the quality of the match between mentee and mentor. responses by gender and citizenship status** | |
|---|---|
| **Theme** | **count** |
| **Women<br>National<br>(n=840)** | 26.3% said Excellent (221 out of 840)<br>27.1% said Very Good (228 out of 840)<br>19.2% said Good (161 out of 840)<br>14.9% said Fair (125 out of 840)<br>12.5% said Poor (105 out of 840) |
| **Women<br>International** | 18.7% said Excellent (83 out of 443)<br>24.8% said Very Good (110 out of 443) |

| | |
|---|---|
| **(n=443)** | 21.9% said Good (97 out of 443)<br>14.9% said Fair (66 out of 443)<br>19.6% said Poor (87 out of 443) |
| **Men<br>National<br>(n=445)** | 25.6% said Excellent (114 out of 445)<br>29.9% said Very Good (133 out of 445)<br>18.7% said Good (83 out of 445)<br>13.7% said Fair (61 out of 445)<br>12.1% said Poor (54 out of 445) |
| **Men<br>International<br>(n=353)** | 21% said Excellent (74 out of 353)<br>25.8% said Very Good (91 out of 353)<br>21% said Good (74 out of 353)<br>15.3% said Fair (54 out of 353)<br>17% said Poor (60 out of 353) |

**Table S38. Do you (trainee mentee) find your interactions with your Mentor constructive? by mentee citizenship status and gender**

| Theme | count |
|---|---|
| **Women<br>National (n=832)** | 73% Yes (607 out of 832)<br>27% No (225 out of 832) |
| **Women<br>International (n=440)** | 66.1% Yes (291 out of 440)<br>33.9% No (149 out of 440) |
| **Men<br>National (n=445)** | 76.6% Yes (341 out of 445)<br>23.4% No (104 out of 445) |
| **Men<br>International (n=352)** | 67% Yes (236 out of 352)<br>33% No (116 out of 352) |

**Table S39. Do you (trainee mentee) feel happy and satisfied with your current research and position as a trainee mentee, analysis by mentee citizenship status and gender**

| Theme | count |
|---|---|
| **Women<br>National (n=765)** | 65.6% Yes (502 out of 765)<br>34.4% No (263 out of 765) |
| **Women<br>International (n=387)** | 58.4% Yes (226 out of 387)<br>41.6% No (161 out of 387) |
| **Men<br>National (n=407)** | 68.3% Yes (278 out of 407)<br>31.7% No (129 out of 407) |
| **Men<br>International (n=314)** | 65.3% Yes (205 out of 314)<br>34.7% No (109 out of 314) |

| **Table S40. Do you (trainee mentee) feel optimistic about your future career? Responses by gender and citizenship in country of research status** | |
|---|---|
| **Theme** | **count** |
| **Women**<br>**National (n=764)** | 69.6% Yes (532 out of 764)<br>30.4% No (232 out of 764) |
| **Women**<br>**International (n=387)** | 60.7% Yes (235 out of 387)<br>39.3% No (152 out of 387) |
| **Men**<br>**National (n=409)** | 74.3% Yes (304 out of 409)<br>25.7% No (105 out of 409) |
| **Men**<br>**International (n=314)** | 68.2% Yes (214 out of 314)<br>31.8% No (100 out of 314) |

| **Table S41. Overall, to what extent do you feel that your current mentor is meeting your expectations? by gender** on a Likert Scale (1 lowest-5 highest) | |
|---|---|
| **Theme** | **count** |
| **Total (n=2114)** | 14.8% rated 1 (lowest score) (313 out of 2114)<br>18.4% rated 2 (389 out of 2114)<br>18.3% rated 3 (386 out of 2114)<br>31.4% rated 4 (663 out of 2114)<br>17.1% rated 5 (highest score) (363 out of 2114) |
| **Women Mentee (n=1292)** | 15% rated 1 (lowest score) (194 out of 1292)<br>19% rated 2 (245 out of 1292)<br>19% rated 3 (246 out of 1292)<br>29.7% rated 4 (384 out of 1292)<br>17.3% rated 5 (highest score) (223 out of 1292) |
| **Men Mentee (n=802)** | 13.8% rated 1 (lowest score) (111 out of 802)<br>17.6% rated 2 (141 out of 802)<br>17.2% rated 3 (138 out of 802)<br>34.3% rated 4 (275 out of 802)<br>17.1% rated 5 (highest score) (137 out of 802) |
| **Non-binary or did not disclose gender (n=15)** | 40% rated 1 (lowest score) (6 out of 15)<br>13.3% rated 2 (2 out of 15)<br>6.7% rated 3 (1 out of 15)<br>26.7% rated 4 (4 out of 15)<br>13.3% rated 5 (highest score) (2 out of 15) |
| **Did not respond to this survey Question** | 0% (0 out of 2114) |

| **Table S42. Overall, to what extent do you feel that your current mentor is meeting your expectations? by citizenship status** on a Likert Scale (1 lowest-5 highest) | |
|---|---|
| **Theme** | **count** |
| **Total (n=2114)** | 14.8% rated 1 (lowest score) (313 out of 2114)<br>18.4% rated 2 (389 out of 2114) |

| | |
|---|---|
| | 18.3% rated 3 (386 out of 2114)<br>31.4% rated 4 (663 out of 2114)<br>17.1% rated 5 (highest score) (363 out of 2114) |
| **Non-Citizen (international) Mentee (n=807)** | 19.1% rated 1 (lowest score) (154 out of 807)<br>18.5% rated 2 (149 out of 807)<br>17.6% rated 3 (142 out of 807)<br>29.7% rated 4 (240 out of 807)<br>15.1% rated 5 (highest score) (122 out of 807) |
| **Citizen (national) Mentee (n=1304)** | 12.2% rated 1 (lowest score) (159 out of 1304)<br>18.4% rated 2 (240 out of 1304)<br>18.6% rated 3 (243 out of 1304)<br>32.4% rated 4 (422 out of 1304)<br>18.4% rated 5 (highest score) (240 out of 1304) |
| **Did not respond to this survey Question** | 0% (0 out of 2114) |

**Table S43. Overall, to what extent do you (the mentee) feel that your current mentor is meeting your expectations? by gender and citizenship status** on a Likert Scale (1 lowest-5 highest)

| Theme | count |
|---|---|
| **Non-Citizen (international) Women Mentee (n=443)** | 20.3% rated 1 (lowest score) (90 out of 443)<br>19.6% rated 2 (87 out of 443)<br>17.8% rated 3 (79 out of 443)<br>26.2% rated 4 (116 out of 443)<br>16% rated 5 (highest score) (71 out of 443) |
| **Non-Citizen (international) Men Mentee (n=356)** | 16.9% rated 1 (lowest score) (60 out of 356)<br>17.1% rated 2 (61 out of 356)<br>17.1% rated 3 (61 out of 356)<br>34.8% rated 4 (124 out of 356)<br>14% rated 5 (highest score) (50 out of 356) |
| **Citizen (national) Women Mentee (n=843)** | 12.2% rated 1 (lowest score) (103 out of 843)<br>18.5% rated 2 (156 out of 843)<br>19.9% rated 3 (168 out of 843)<br>31.3% rated 4 (264 out of 843)<br>18% rated 5 (highest score) (152 out of 843) |
| **Citizen (national) Men Mentee (n=446)** | 11.4% rated 1 (lowest score) (51 out of 446)<br>17.9% rated 2 (80 out of 446)<br>17.3% rated 3 (77 out of 446)<br>33.9% rated 4 (151 out of 446)<br>19.5% rated 5 (highest score) (87 out of 446) |

**Table S44. Mentor established a relationship based on trust with mentee by gender** on a Likert Scale (1 lowest-5 highest)

| Theme | count |
|---|---|
| **Total (n=2113)** | 11.7% rated 1 (lowest score) (247 out of 2113) |

| | |
|---|---|
| | 13.8% rated 2 (292 out of 2113)<br>13% rated 3 (274 out of 2113)<br>22.9% rated 4 (485 out of 2113)<br>38.6% rated 5 (highest score) (815 out of 2113) |
| **Women Mentee (n=1283)** | 12.2% rated 1 (lowest score) (157 out of 1283)<br>14% rated 2 (180 out of 1283)<br>13.2% rated 3 (169 out of 1283)<br>21.3% rated 4 (273 out of 1283)<br>39.3% rated 5 (highest score) (504 out of 1283) |
| **Men Mentee (n=801)** | 10.6% rated 1 (lowest score) (85 out of 801)<br>13.5% rated 2 (108 out of 801)<br>12.4% rated 3 (99 out of 801)<br>25.7% rated 4 (206 out of 801)<br>37.8% rated 5 (highest score) (303 out of 801) |
| **Non-binary or did not disclose gender (n=24)** | 16.7% rated 1 (lowest score) (4 out of 24)<br>8.3% rated 2 (2 out of 24)<br>20.8% rated 3 (5 out of 24)<br>20.8% rated 4 (5 out of 24)<br>33.3% rated 5 (highest score) (8 out of 24) |
| **Did not respond to this survey Question** | 0.05% (1 out of 2114) |

| **Table S45. Mentor established a relationship based on trust with the mentee. by citizenship status** on a Likert Scale (1 lowest-5 highest) | |
|---|---|
| **Theme** | **count** |
| **Total (n=2112)** | 11.7% rated 1 (lowest score) (247 out of 2112)<br>13.8% rated 2 (292 out of 2112)<br>13% rated 3 (274 out of 2112)<br>22.9% rated 4 (485 out of 2112)<br>38.6% rated 5 (highest score) (815 out of 2112) |
| **Non-Citizen (international) Mentee (n=806)** | 13% rated 1 (lowest score) (105 out of 806)<br>15.8% rated 2 (127 out of 806)<br>13.6% rated 3 (110 out of 806)<br>22% rated 4 (177 out of 806)<br>35.6% rated 5 (highest score) (287 out of 806) |
| **Citizen (national) Mentee (n=1304)** | 10.9% rated 1 (lowest score) (142 out of 1304)<br>12.6% rated 2 (164 out of 1304)<br>12.5% rated 3 (163 out of 1304)<br>23.5% rated 4 (307 out of 1304)<br>40.5% rated 5 (highest score) (528 out of 1304) |
| **Did not respond to this survey Question** | 0.09% (2 out of 2114) |

| **Table S46. Mentor established a relationship based on trust with the mentee. by citizenship status** on a Likert Scale (1 lowest-5 highest) | |
|---|---|
| **Theme** | **count** |
| **Non-Citizen (international) Women** | 13.8% rated 1 (lowest score) (61 out of 443) |

| | |
|---|---|
| **Mentee (n=443)** | 17.2% rated 2 (76 out of 443)<br>12.4% rated 3 (55 out of 443)<br>19.6% rated 4 (87 out of 443)<br>37% rated 5 (highest score) (164 out of 443) |
| **Non-Citizen (international) Men Mentee (n=355)** | 12.1% rated 1 (lowest score) (43 out of 355)<br>13.8% rated 2 (49 out of 355)<br>14.6% rated 3 (52 out of 355)<br>25.4% rated 4 (90 out of 355)<br>34.1% rated 5 (highest score) (121 out of 355) |
| **Citizen (national) Women Mentee (n=840)** | 11.4% rated 1 (lowest score) (96 out of 840)<br>12.4% rated 2 (104 out of 840)<br>13.6% rated 3 (114 out of 840)<br>22.1% rated 4 (186 out of 840)<br>40.5% rated 5 (highest score) (340 out of 840) |
| **Citizen (national) Men Mentee (n=446)** | 9.4% rated 1 (lowest score) (42 out of 446)<br>13.2% rated 2 (59 out of 446)<br>10.5% rated 3 (47 out of 446)<br>26% rated 4 (116 out of 446)<br>40.8% rated 5 (highest score) (182 out of 446) |

**Table S47. Mentor uses active listening (meaning identifying and accommodating different communication styles and employing strategies to improve communication with you)**
on a Likert Scale (1 lowest-5 highest)

| Theme | count |
|---|---|
| **Total (n=2110)** | 15.9% rated 1 (lowest score) (336 out of 2110)<br>18.4% rated 2 (389 out of 2110)<br>16.5% rated 3 (349 out of 2110)<br>25.3% rated 4 (533 out of 2110)<br>23.8% rated 5 (highest score) (503 out of 2110) |
| **Women Mentee (n=1281)** | 16.2% rated 1 (lowest score) (207 out of 1281)<br>18.6% rated 2 (238 out of 1281)<br>16.4% rated 3 (210 out of 1281)<br>24.2% rated 4 (310 out of 1281)<br>24.7% rated 5 (highest score) (316 out of 1281) |
| **Men Mentee (n=801)** | 15.4% rated 1 (lowest score) (123 out of 801)<br>18% rated 2 (144 out of 801)<br>17% rated 3 (136 out of 801)<br>27% rated 4 (216 out of 801)<br>22.7% rated 5 (highest score) (182 out of 801) |
| **Non-binary or did not disclose gender (n=24)** | 20.8% rated 1 (lowest score) (5 out of 24)<br>25% rated 2 (6 out of 24)<br>4.2% rated 3 (1 out of 24)<br>29.2% rated 4 (7 out of 24)<br>20.8% rated 5 (highest score) (5 out of 24) |
| **Did not respond to this survey Question** | 0.2% (4 out of 2114) |

| **Table S48. Mentor uses active listening (meaning identifying and accommodating different communication styles and employing strategies to improve communication with you) by citizenship status** on a Likert Scale (1 lowest-5 highest) | |
|---|---|
| **Theme** | **count** |
| **Total (n=2110)** | 15.9% rated 1 (lowest score) (336 out of 2110)<br>18.4% rated 2 (389 out of 2110)<br>16.5% rated 3 (349 out of 2110)<br>25.3% rated 4 (533 out of 2110)<br>23.8% rated 5 (highest score) (503 out of 2110) |
| **Non-Citizen (international) Mentee (n=805)** | 18% rated 1 (lowest score) (145 out of 805)<br>18.6% rated 2 (150 out of 805)<br>16.3% rated 3 (131 out of 805)<br>23% rated 4 (185 out of 805)<br>24.1% rated 5 (highest score) (194 out of 805) |
| **Citizen (national) Mentee (n=1303)** | 14.7% rated 1 (lowest score) (191 out of 1303)<br>18.3% rated 2 (239 out of 1303)<br>16.6% rated 3 (216 out of 1303)<br>26.7% rated 4 (348 out of 1303)<br>23.7% rated 5 (highest score) (309 out of 1303) |
| **Did not respond to this survey Question** | 0.2% (4 out of 2114) |

| **Table S49. Mentor uses active listening (meaning identifying and accommodating different communication styles and employing strategies to improve communication with you) by citizenship status** on a Likert Scale (1 lowest-5 highest) | |
|---|---|
| **Theme** | **count** |
| **Non-Citizen (international) Women Mentee (n=442)** | 18.8% rated 1 (lowest score) (83 out of 442)<br>18.1% rated 2 (80 out of 442)<br>16.1% rated 3 (71 out of 442)<br>20.8% rated 4 (92 out of 442)<br>26.2% rated 5 (highest score) (116 out of 442) |
| **Non-Citizen (international) Men Mentee (n=355)** | 16.6% rated 1 (lowest score) (59 out of 355)<br>19.4% rated 2 (69 out of 355)<br>16.3% rated 3 (58 out of 355)<br>25.9% rated 4 (92 out of 355)<br>21.7% rated 5 (highest score) (77 out of 355) |
| **Citizen (national) Women Mentee (n=839)** | 14.8% rated 1 (lowest score) (124 out of 839)<br>18.8% rated 2 (158 out of 839)<br>16.6% rated 3 (139 out of 839)<br>26% rated 4 (218 out of 839)<br>23.8% rated 5 (highest score) (200 out of 839) |
| **Citizen (national) Men Mentee (n=446)** | 14.3% rated 1 (lowest score) (64 out of 446)<br>16.8% rated 2 (75 out of 446)<br>17.5% rated 3 (78 out of 446)<br>27.8% rated 4 (124 out of 446)<br>23.5% rated 5 (highest score) (105 out of 446) |

| **Table S50. Mentor coordinates effectively with other mentors with whom you work**<br>on a Likert Scale (1 lowest-5 highest) | |
|---|---|
| **Theme** | **count** |
| **Total (n=2051)** | 16.8% rated 1 (lowest score) (344 out of 2051)<br>18.2% rated 2 (373 out of 2051)<br>23.3% rated 3 (478 out of 2051)<br>23.6% rated 4 (484 out of 2051)<br>18.1% rated 5 (highest score) (372 out of 2051) |
| **Women Mentee (n=1246)** | 17.7% rated 1 (lowest score) (220 out of 1246)<br>17.8% rated 2 (222 out of 1246)<br>22.5% rated 3 (280 out of 1246)<br>23.8% rated 4 (297 out of 1246)<br>18.2% rated 5 (highest score) (227 out of 1246) |
| **Men Mentee (n=777)** | 14.9% rated 1 (lowest score) (116 out of 777)<br>18.8% rated 2 (146 out of 777)<br>24.6% rated 3 (191 out of 777)<br>23.3% rated 4 (181 out of 777)<br>18.4% rated 5 (highest score) (143 out of 777) |
| **Non-binary or did not disclose gender (n=24)** | 25% rated 1 (lowest score) (6 out of 24)<br>20.8% rated 2 (5 out of 24)<br>20.8% rated 3 (5 out of 24)<br>25% rated 4 (6 out of 24)<br>8.3% rated 5 (highest score) (2 out of 24) |
| **Did not respond to this survey Question** | 3% (63 out of 2114) |

| **Table S51. Mentor coordinates effectively with other mentors with whom you work**<br>on a Likert Scale (1 lowest-5 highest) | |
|---|---|
| **Theme** | **Count** |
| **Total (n=2051)** | 16.8% rated 1 (lowest score) (344 out of 2051)<br>18.2% rated 2 (373 out of 2051)<br>23.3% rated 3 (478 out of 2051)<br>23.6% rated 4 (484 out of 2051)<br>18.1% rated 5 (highest score) (372 out of 2051) |
| **Non-Citizen (international) Mentee (n=779)** | 19.9% rated 1 (lowest score) (155 out of 779)<br>19% rated 2 (148 out of 779)<br>22.6% rated 3 (176 out of 779)<br>22% rated 4 (171 out of 779)<br>16.6% rated 5 (highest score) (129 out of 779) |
| **Citizen (national) Mentee (n=1270)** | 14.8% rated 1 (lowest score) (188 out of 1270)<br>17.7% rated 2 (225 out of 1270)<br>23.7% rated 3 (301 out of 1270)<br>24.6% rated 4 (313 out of 1270)<br>19.1% rated 5 (highest score) (243 out of 1270) |
| **Did not respond to this survey Question** | 3% (63 out of 2114) |

| **Table S52. Mentor coordinates effectively with other mentors with whom you work**<br>on a Likert Scale (1 lowest-5 highest) | |
|---|---|
| **Theme** | **count** |
| **Non-Citizen (international) Women Mentee (n=430)** | 21.4% rated 1 (lowest score) (92 out of 430)<br>18.4% rated 2 (79 out of 430)<br>20.5% rated 3 (88 out of 430)<br>23.5% rated 4 (101 out of 430)<br>16.3% rated 5 (highest score) (70 out of 430) |
| **Non-Citizen (international) Men Mentee (n=341)** | 17.6% rated 1 (lowest score) (60 out of 341)<br>19.6% rated 2 (67 out of 341)<br>25.2% rated 3 (86 out of 341)<br>20.2% rated 4 (69 out of 341)<br>17.3% rated 5 (highest score) (59 out of 341) |
| **Citizen (national) Women Mentee (n=816)** | 15.7% rated 1 (lowest score) (128 out of 816)<br>17.5% rated 2 (143 out of 816)<br>23.5% rated 3 (192 out of 816)<br>24% rated 4 (196 out of 816)<br>19.2% rated 5 (highest score) (157 out of 816) |
| **Citizen (national) Men Mentee (n=436)** | 12.8% rated 1 (lowest score) (56 out of 436)<br>18.1% rated 2 (79 out of 436)<br>24.1% rated 3 (105 out of 436)<br>25.7% rated 4 (112 out of 436)<br>19.3% rated 5 (highest score) (84 out of 436) |

| **Table S53. Mentor works with you to set clear expectations of the mentoring relationship**<br>on a Likert Scale (1 lowest-5 highest) | |
|---|---|
| **Theme** | **Count** |
| **Total (n=2096)** | 21.3% rated 1 (lowest score) (446 out of 2096)<br>20.3% rated 2 (426 out of 2096)<br>18.4% rated 3 (386 out of 2096)<br>23.3% rated 4 (488 out of 2096)<br>16.7% rated 5 (highest score) (350 out of 2096) |
| **Women Mentee (n=1272)** | 22% rated 1 (lowest score) (280 out of 1272)<br>19.9% rated 2 (253 out of 1272)<br>19% rated 3 (242 out of 1272)<br>22.1% rated 4 (281 out of 1272)<br>17% rated 5 (highest score) (216 out of 1272) |
| **Men Mentee (n=796)** | 20.1% rated 1 (lowest score) (160 out of 796)<br>20.9% rated 2 (166 out of 796)<br>17.1% rated 3 (136 out of 796)<br>25.5% rated 4 (203 out of 796)<br>16.5% rated 5 (highest score) (131 out of 796) |
| **Non-binary or did not disclose gender (n=24)** | 20.8% rated 1 (lowest score) (5 out of 24)<br>25% rated 2 (6 out of 24)<br>25% rated 3 (6 out of 24)<br>16.7% rated 4 (4 out of 24) |

| | |
|---|---|
| | 12.5% rated 5 (highest score) (3 out of 24) |
| **Did not respond to this survey Question** | 0.85% (18 out of 2114) |

| **Table S54. Mentor works with you to set clear expectations of the mentoring relationship by citizenship status** on a Likert Scale (1 lowest-5 highest) | |
|---|---|
| **Theme** | **Count** |
| **Total (n=2096)** | 21.3% rated 1 (lowest score) (446 out of 2096)<br>20.3% rated 2 (426 out of 2096)<br>18.4% rated 3 (386 out of 2096)<br>23.3% rated 4 (488 out of 2096)<br>16.7% rated 5 (highest score) (350 out of 2096) |
| **Non-Citizen (international) Mentee (n=799)** | 24.7% rated 1 (lowest score) (197 out of 799)<br>19.3% rated 2 (154 out of 799)<br>16.8% rated 3 (134 out of 799)<br>21.7% rated 4 (173 out of 799)<br>17.6% rated 5 (highest score) (141 out of 799) |
| **Citizen (national) Mentee (n=1295)** | 19.2% rated 1 (lowest score) (249 out of 1295)<br>20.9% rated 2 (271 out of 1295)<br>19.4% rated 3 (251 out of 1295)<br>24.3% rated 4 (315 out of 1295)<br>16.1% rated 5 (highest score) (209 out of 1295) |
| **Did not respond to this survey Question** | 0.85% (18 out of 2114) |

| **Table S55. Mentor works with you to set clear expectations of the mentoring relationship by citizenship status** on a Likert Scale (1 lowest-5 highest) | |
|---|---|
| **Theme** | **count** |
| **Non-Citizen (international) Women Mentee (n=438)** | 26.9% rated 1 (lowest score) (118 out of 438)<br>18.3% rated 2 (80 out of 438)<br>16.9% rated 3 (74 out of 438)<br>20.3% rated 4 (89 out of 438)<br>17.6% rated 5 (highest score) (77 out of 438) |
| **Non-Citizen (international) Men Mentee (n=353)** | 21.8% rated 1 (lowest score) (77 out of 353)<br>20.4% rated 2 (72 out of 353)<br>16.1% rated 3 (57 out of 353)<br>23.5% rated 4 (83 out of 353)<br>18.1% rated 5 (highest score) (64 out of 353) |
| **Citizen (national) Women Mentee (n=834)** | 19.4% rated 1 (lowest score) (162 out of 834)<br>20.7% rated 2 (173 out of 834)<br>20.1% rated 3 (168 out of 834)<br>23% rated 4 (192 out of 834)<br>16.7% rated 5 (highest score) (139 out of 834) |
| **Citizen (national) Men Mentee (n=443)** | 18.7% rated 1 (lowest score) (83 out of 443)<br>21.2% rated 2 (94 out of 443) |

| | |
|---|---|
| | 17.8% rated 3 (79 out of 443)<br>27.1% rated 4 (120 out of 443)<br>15.1% rated 5 (highest score) (67 out of 443) |

| **Table S56. Mentor provides you with respectful, positive affirmation and constructive feedback**<br>on a Likert Scale (1 lowest-5 highest) | |
|---|---|
| **Theme** | **Count** |
| **Total (n=2089)** | 15.2% rated 1 (lowest score) (317 out of 2089)<br>14.1% rated 2 (295 out of 2089)<br>15.3% rated 3 (319 out of 2089)<br>24.2% rated 4 (506 out of 2089)<br>31.2% rated 5 (highest score) (652 out of 2089) |
| **Women Mentee (n=1283)** | 16.4% rated 1 (lowest score) (211 out of 1283)<br>14.3% rated 2 (183 out of 1283)<br>14.6% rated 3 (187 out of 1283)<br>23.5% rated 4 (302 out of 1283)<br>31.2% rated 5 (highest score) (400 out of 1283) |
| **Men Mentee (n=802)** | 13% rated 1 (lowest score) (104 out of 802)<br>13.8% rated 2 (111 out of 802)<br>16.3% rated 3 (131 out of 802)<br>25.4% rated 4 (204 out of 802)<br>31.4% rated 5 (highest score) (252 out of 802) |
| **Non-binary or did not disclose gender (n=24)** | 25% rated 1 (lowest score) (6 out of 24)<br>16.7% rated 2 (4 out of 24)<br>16.7% rated 3 (4 out of 24)<br>20.8% rated 4 (5 out of 24)<br>20.8% rated 5 (highest score) (5 out of 24) |
| **Did not respond to this survey Question** | 1.2% (25 out of 2114) |

| **Table S57. Mentor provides you with respectful, positive affirmation and constructive feedback**<br>**by citizenship status** on a Likert Scale (1 lowest-5 highest) | |
|---|---|
| **Theme** | **Count** |
| **Total (n=2113)** | 15.3% rated 1 (lowest score) (323 out of 2113)<br>14.2% rated 2 (299 out of 2113)<br>15.3% rated 3 (323 out of 2113)<br>24.2% rated 4 (511 out of 2113)<br>31.1% rated 5 (highest score) (657 out of 2113) |
| **Non-Citizen (international) Mentee (n=807)** | 17.2% rated 1 (lowest score) (139 out of 807)<br>14.7% rated 2 (119 out of 807)<br>15.1% rated 3 (122 out of 807)<br>22.1% rated 4 (178 out of 807)<br>30.9% rated 5 (highest score) (249 out of 807) |

| | |
|---|---|
| **Citizen (national) Mentee (n=1304)** | 14.1% rated 1 (lowest score) (184 out of 1304)<br>13.8% rated 2 (180 out of 1304)<br>15.3% rated 3 (200 out of 1304)<br>25.5% rated 4 (332 out of 1304)<br>31.3% rated 5 (highest score) (408 out of 1304) |
| **Did not respond to this survey Question** | 0.04% (1 out of 2114) |

| **Table S58. Mentor provides you with respectful, positive affirmation and constructive feedback by citizenship status** on a Likert Scale (1 lowest-5 highest) | |
|---|---|
| **Theme** | **count** |
| **Non-Citizen (international) Women Mentee (n=443)** | 18.7% rated 1 (lowest score) (83 out of 443)<br>14% rated 2 (62 out of 443)<br>15.8% rated 3 (70 out of 443)<br>21.7% rated 4 (96 out of 443)<br>29.8% rated 5 (highest score) (132 out of 443) |
| **Non-Citizen (international) Men Mentee (n=356)** | 14.9% rated 1 (lowest score) (53 out of 356)<br>15.7% rated 2 (56 out of 356)<br>14.3% rated 3 (51 out of 356)<br>22.5% rated 4 (80 out of 356)<br>32.6% rated 5 (highest score) (116 out of 356) |
| **Citizen (national) Women Mentee (n=840)** | 15.2% rated 1 (lowest score) (128 out of 840)<br>14.4% rated 2 (121 out of 840)<br>13.9% rated 3 (117 out of 840)<br>24.5% rated 4 (206 out of 840)<br>31.9% rated 5 (highest score) (268 out of 840) |
| **Citizen (national) Men Mentee (n=446)** | 11.4% rated 1 (lowest score) (51 out of 446)<br>12.3% rated 2 (55 out of 446)<br>17.9% rated 3 (80 out of 446)<br>27.8% rated 4 (124 out of 446)<br>30.5% rated 5 (highest score) (136 out of 446) |

| **Table S59. Mentor challenges you (trainee mentee) during your career. Analysis by gender** on a Likert Scale (1 lowest-5 highest) | |
|---|---|
| **Theme** | **Count** |
| **Total (n=2094)** | 8.1% rated 1 (lowest score) (170 out of 2094)<br>12.5% rated 2 (262 out of 2094)<br>18.7% rated 3 (392 out of 2094)<br>33.1% rated 4 (693 out of 2094)<br>27.6% rated 5 (highest score) (577 out of 2094) |
| **Women Mentee (n=1271)** | 8.3% rated 1 (lowest score) (105 out of 1271)<br>12.1% rated 2 (154 out of 1271)<br>19.1% rated 3 (243 out of 1271)<br>31.8% rated 4 (404 out of 1271)<br>28.7% rated 5 (highest score) (365 out of 1271) |

| | |
|---|---|
| **Men Mentee (n=795)** | 7.9% rated 1 (lowest score) (63 out of 795)<br>12.8% rated 2 (102 out of 795)<br>18.4% rated 3 (146 out of 795)<br>35.2% rated 4 (280 out of 795)<br>25.7% rated 5 (highest score) (204 out of 795) |
| **Non-binary or did not disclose gender (n=24)** | 8.3% rated 1 (lowest score) (2 out of 24)<br>20.8% rated 2 (5 out of 24)<br>8.3% rated 3 (2 out of 24)<br>33.3% rated 4 (8 out of 24)<br>29.2% rated 5 (highest score) (7 out of 24) |
| **Did not respond to this survey Question** | 0.9% (20 out of 2114) |

| **Table S60. Mentor challenges you (trainee mentee) during your career. by gender** on a Likert Scale (1 lowest-5 highest) | |
|---|---|
| **Theme** | **Count** |
| **Total (n=2094)** | 8.1% rated 1 (lowest score) (170 out of 2094)<br>12.5% rated 2 (262 out of 2094)<br>18.7% rated 3 (392 out of 2094)<br>33.1% rated 4 (693 out of 2094)<br>27.6% rated 5 (highest score) (577 out of 2094) |
| **Non-Citizen (international) Mentee (n=796)** | 9.4% rated 1 (lowest score) (75 out of 796)<br>13.6% rated 2 (108 out of 796)<br>19.1% rated 3 (152 out of 796)<br>33.8% rated 4 (269 out of 796)<br>24.1% rated 5 (highest score) (192 out of 796) |
| **Citizen (national) Mentee (n=1296)** | 7.3% rated 1 (lowest score) (94 out of 1296)<br>11.9% rated 2 (154 out of 1296)<br>18.4% rated 3 (239 out of 1296)<br>32.7% rated 4 (424 out of 1296)<br>29.7% rated 5 (highest score) (385 out of 1296) |
| **Did not respond to this survey Question** | 0.9% (20 out of 2114) |

| **Table S61. Mentor challenges you (trainee mentee) during your career. by gender** on a Likert Scale (1 lowest-5 highest) | |
|---|---|
| **Theme** | **count** |
| **Non-Citizen (international) Women Mentee (n=437)** | 10.1% rated 1 (lowest score) (44 out of 437)<br>13.3% rated 2 (58 out of 437)<br>20.4% rated 3 (89 out of 437)<br>32% rated 4 (140 out of 437)<br>24.3% rated 5 (highest score) (106 out of 437) |
| **Non-Citizen (international) Men Mentee (n=351)** | 8.8% rated 1 (lowest score) (31 out of 351)<br>13.1% rated 2 (46 out of 351)<br>17.9% rated 3 (63 out of 351) |

| | |
|---|---|
| | 36.5% rated 4 (128 out of 351)<br>23.6% rated 5 (highest score) (83 out of 351) |
| **Citizen (national) Women Mentee (n=834)** | 7.3% rated 1 (lowest score) (61 out of 834)<br>11.5% rated 2 (96 out of 834)<br>18.5% rated 3 (154 out of 834)<br>31.7% rated 4 (264 out of 834)<br>31.1% rated 5 (highest score) (259 out of 834) |
| **Citizen (national) Men Mentee (n=444)** | 7.2% rated 1 (lowest score) (32 out of 444)<br>12.6% rated 2 (56 out of 444)<br>18.7% rated 3 (83 out of 444)<br>34.2% rated 4 (152 out of 444)<br>27.3% rated 5 (highest score) (121 out of 444) |

**Table S62. Mentor aligns his/her expectations with your own.**
on Likert Scale (1 lowest-5 highest)

| Theme | Count |
|---|---|
| **Total (n=2095)** | 14.1% rated 1 (lowest score) (296 out of 2095)<br>19.3% rated 2 (404 out of 2095)<br>19.4% rated 3 (406 out of 2095)<br>28.1% rated 4 (589 out of 2095)<br>19.1% rated 5 (highest score) (400 out of 2095) |
| **Women Mentee (n=1272)** | 14.9% rated 1 (lowest score) (190 out of 1272)<br>20.4% rated 2 (259 out of 1272)<br>17.8% rated 3 (227 out of 1272)<br>27.4% rated 4 (349 out of 1272)<br>19.4% rated 5 (highest score) (247 out of 1272) |
| **Men Mentee (n=795)** | 12.6% rated 1 (lowest score) (100 out of 795)<br>17.6% rated 2 (140 out of 795)<br>21.8% rated 3 (173 out of 795)<br>29.3% rated 4 (233 out of 795)<br>18.7% rated 5 (highest score) (149 out of 795) |
| **Non-binary or did not disclose gender (n=24)** | 20.8% rated 1 (lowest score) (5 out of 24)<br>20.8% rated 2 (5 out of 24)<br>12.5% rated 3 (3 out of 24)<br>29.2% rated 4 (7 out of 24)<br>16.7% rated 5 (highest score) (4 out of 24) |
| **Did not respond to this survey Question** | 0.9% (19 out of 2114) |

**Table S63. Mentor aligns his/her expectations with your own.**
on Likert Scale (1 lowest-5 highest)

| Theme | Count |
|---|---|
| **Total (n=2095)** | 14.1% rated 1 (lowest score) (296 out of 2095)<br>19.3% rated 2 (404 out of 2095)<br>19.4% rated 3 (406 out of 2095) |

| | |
|---|---|
| | 28.1% rated 4 (589 out of 2095)<br>19.1% rated 5 (highest score) (400 out of 2095) |
| **Non-Citizen (international) Mentee (n=799)** | 16% rated 1 (lowest score) (128 out of 799)<br>19.9% rated 2 (159 out of 799)<br>18.5% rated 3 (148 out of 799)<br>27.8% rated 4 (222 out of 799)<br>17.8% rated 5 (highest score) (142 out of 799) |
| **Citizen (national) Mentee (n=1294)** | 13% rated 1 (lowest score) (168 out of 1294)<br>18.9% rated 2 (245 out of 1294)<br>19.8% rated 3 (256 out of 1294)<br>28.4% rated 4 (367 out of 1294)<br>19.9% rated 5 (highest score) (258 out of 1294) |
| **Did not respond to this survey Question** | 0.9% (19 out of 2114) |

**Table S64. Mentor aligns his/her expectations with your own.**
on Likert Scale (1 lowest-5 highest)

| Theme | count |
|---|---|
| **Non-Citizen (international) Women Mentee (n=438)** | 17.6% rated 1 (lowest score) (77 out of 438)<br>21.9% rated 2 (96 out of 438)<br>15.5% rated 3 (68 out of 438)<br>26.9% rated 4 (118 out of 438)<br>18% rated 5 (highest score) (79 out of 438) |
| **Non-Citizen (international) Men Mentee (n=353)** | 14.2% rated 1 (lowest score) (50 out of 353)<br>17% rated 2 (60 out of 353)<br>22.1% rated 3 (78 out of 353)<br>28.9% rated 4 (102 out of 353)<br>17.8% rated 5 (highest score) (63 out of 353) |
| **Citizen (national) Women Mentee (n=834)** | 13.5% rated 1 (lowest score) (113 out of 834)<br>19.5% rated 2 (163 out of 834)<br>19.1% rated 3 (159 out of 834)<br>27.7% rated 4 (231 out of 834)<br>20.1% rated 5 (highest score) (168 out of 834) |
| **Citizen (national) Men Mentee (n=442)** | 11.3% rated 1 (lowest score) (50 out of 442)<br>18.1% rated 2 (80 out of 442)<br>21.5% rated 3 (95 out of 442)<br>29.6% rated 4 (131 out of 442)<br>19.5% rated 5 (highest score) (86 out of 442) |

**Table S65. Mentor considers how personal and professional differences may impact expectations.** on a Likert Scale (1 lowest-5 highest)

| Theme | Count |
|---|---|
| **Total (n=2088)** | 16.7% rated 1 (lowest score) (348 out of 2088)<br>16.5% rated 2 (345 out of 2088)<br>21.3% rated 3 (445 out of 2088)<br>24.1% rated 4 (504 out of 2088) |

| | |
|---|---|
| | 21.4% rated 5 (highest score) (446 out of 2088) |
| **Women Mentee (n=1268)** | 18.1% rated 1 (lowest score) (229 out of 1268)<br>17.3% rated 2 (219 out of 1268)<br>20.3% rated 3 (257 out of 1268)<br>23.3% rated 4 (295 out of 1268)<br>21.1% rated 5 (highest score) (268 out of 1268) |
| **Men Mentee (n=792)** | 14.1% rated 1 (lowest score) (112 out of 792)<br>14.8% rated 2 (117 out of 792)<br>23.2% rated 3 (184 out of 792)<br>26% rated 4 (206 out of 792)<br>21.8% rated 5 (highest score) (173 out of 792) |
| **Non-binary or did not disclose gender (n=24)** | 25% rated 1 (lowest score) (6 out of 24)<br>29.2% rated 2 (7 out of 24)<br>12.5% rated 3 (3 out of 24)<br>12.5% rated 4 (3 out of 24)<br>20.8% rated 5 (highest score) (5 out of 24) |
| **Did not respond to this survey Question** | 1.2% (26 out of 2114) |

| **Table S66. Mentor considers how personal and professional differences may impact expectations. by citizenship status.** on a Likert Scale (1 lowest-5 highest) | |
|---|---|
| **Theme** | **Count** |
| **Total (n=2094)** | 16.7% rated 1 (lowest score) (348 out of 2094)<br>16.5% rated 2 (345 out of 2094)<br>21.3% rated 3 (445 out of 2094)<br>24.1% rated 4 (504 out of 2094)<br>21.4% rated 5 (highest score) (446 out of 2094) |
| **Non-Citizen (international) Mentee (n=803)** | 17.4% rated 1 (lowest score) (140 out of 803)<br>16.7% rated 2 (134 out of 803)<br>21.8% rated 3 (175 out of 803)<br>24.5% rated 4 (197 out of 803)<br>19.6% rated 5 (highest score) (157 out of 803) |
| **Citizen (national) Mentee (n=1289)** | 16.1% rated 1 (lowest score) (208 out of 1289)<br>16.3% rated 2 (210 out of 1289)<br>20.9% rated 3 (269 out of 1289)<br>24.3% rated 4 (313 out of 1289)<br>22.4% rated 5 (highest score) (289 out of 1289) |
| **Did not respond to this survey Question** | 0.95% (20 out of 2114) |

| **Table S67. Mentor considers how personal and professional differences may impact expectations. by citizenship status.** on a Likert Scale (1 lowest-5 highest) | |
|---|---|
| **Theme** | **count** |
| **Non-Citizen (international) Women Mentee** | 18.9% rated 1 (lowest score) (83 out of 438)<br>17.6% rated 2 (77 out of 438) |

| | |
|---|---|
| **(n=438)** | 21.2% rated 3 (93 out of 438)<br>22.8% rated 4 (100 out of 438)<br>19.4% rated 5 (highest score) (85 out of 438) |
| **Non-Citizen (international) Men Mentee (n=353)** | 15.6% rated 1 (lowest score) (55 out of 353)<br>15.3% rated 2 (54 out of 53)<br>22.7% rated 3 (80 out of 353)<br>26.1% rated 4 (92 out of 353)<br>20.4% rated 5 (highest score) (72 out of 353) |
| **Citizen (national) Women Mentee (n=830)** | 17.6% rated 1 (lowest score) (146 out of 830)<br>17.1% rated 2 (142 out of 830)<br>19.8% rated 3 (164 out of 830)<br>23.5% rated 4 (195 out of 830)<br>22% rated 5 (highest score) (183 out of 830) |
| **Citizen (national) Men Mentee (n=441)** | 12.9% rated 1 (lowest score) (57 out of 441)<br>14.3% rated 2 (63 out of 441)<br>23.6% rated 3 (104 out of 441)<br>26.3% rated 4 (116 out of 441)<br>22.9% rated 5 (highest score) (101 out of 441) |

**Table S68. Mentor Works with you to set research goals. By gender**
on a Likert Scale (1 lowest-5 highest)

| Theme | Count |
|---|---|
| **Total (n=2109)** | 13.2% rated 1 (lowest score) (279 out of 2109)<br>16.5% rated 2 (348 out of 2109)<br>16.1% rated 3 (340 out of 2109)<br>26.9% rated 4 (568 out of 2109)<br>27.2% rated 5 (highest score) (574 out of 2109) |
| **Women Mentee (n=1282)** | 13.4% rated 1 (lowest score) (172 out of 1282)<br>16.7% rated 2 (214 out of 1282)<br>15.7% rated 3 (201 out of 1282)<br>27.1% rated 4 (347 out of 1282)<br>27.1% rated 5 (highest score) (348 out of 1282) |
| **Men Mentee (n=799)** | 13% rated 1 (lowest score) (104 out of 799)<br>15.8% rated 2 (126 out of 799)<br>16.5% rated 3 (132 out of 799)<br>27% rated 4 (216 out of 799)<br>27.7% rated 5 (highest score) (221 out of 799) |
| **Non-binary or did not disclose gender (n=24)** | 8.3% rated 1 (lowest score) (2 out of 24)<br>29.2% rated 2 (7 out of 24)<br>25% rated 3 (6 out of 24)<br>16.7% rated 4 (4 out of 24)<br>20.8% rated 5 (highest score) (5 out of 24) |
| **Did not respond to this survey Question** | 0.2% (5 out of 2114) |

| **Table S69. Mentor Works with you to set research goals. By citizenship status**<br>on a Likert Scale (1 lowest-5 highest) | |
|---|---|
| **Theme** | **Count** |
| **Total (n=2109)** | 13.2% rated 1 (lowest score) (279 out of 2109)<br>16.5% rated 2 (348 out of 2109)<br>16.1% rated 3 (340 out of 2109)<br>26.9% rated 4 (568 out of 2109)<br>27.2% rated 5 (highest score) (574 out of 2109) |
| **Non-Citizen (international) Mentee (n=804)** | 16% rated 1 (lowest score) (129 out of 804)<br>17.3% rated 2 (139 out of 804)<br>15.3% rated 3 (123 out of 804)<br>27.6% rated 4 (222 out of 804)<br>23.8% rated 5 (highest score) (191 out of 804) |
| **Citizen (national) Mentee (n=1303)** | 11.5% rated 1 (lowest score) (150 out of 1303)<br>16% rated 2 (209 out of 1303)<br>16.6% rated 3 (216 out of 1303)<br>26.5% rated 4 (345 out of 1303)<br>29.4% rated 5 (highest score) (383 out of 1303) |
| **Did not respond to this survey Question** | 0.2% (5 out of 2114) |

| **Table S70. Mentor Works with you to set research goals. Analysis by citizenship status**<br>on a Likert Scale (1 lowest-5 highest) | |
|---|---|
| **Theme** | **count** |
| **Non-Citizen (international) Women Mentee (n=442)** | 16.5% rated 1 (lowest score) (73 out of 442)<br>18.6% rated 2 (82 out of 442)<br>14.9% rated 3 (66 out of 442)<br>26.5% rated 4 (117 out of 442)<br>23.5% rated 5 (highest score) (104 out of 442) |
| **Non-Citizen (international) Men Mentee (n=354)** | 15.3% rated 1 (lowest score) (54 out of 354)<br>15.5% rated 2 (55 out of 354)<br>15.5% rated 3 (55 out of 354)<br>29.4% rated 4 (104 out of 354)<br>24.3% rated 5 (highest score) (86 out of 354) |
| **Citizen (national) Women Mentee (n=840)** | 11.8% rated 1 (lowest score) (99 out of 840)<br>15.7% rated 2 (132 out of 840)<br>16.1% rated 3 (135 out of 840)<br>27.4% rated 4 (230 out of 840)<br>29% rated 5 (highest score) (244 out of 840) |
| **Citizen (national) Men Mentee (n=445)** | 11.2% rated 1 (lowest score) (50 out of 445)<br>16% rated 2 (71 out of 445)<br>17.3% rated 3 (77 out of 445)<br>25.2% rated 4 (112 out of 445)<br>30.3% rated 5 (highest score) (135 out of 445) |

| Table S71. Mentor ensures a working environment free from discrimination and harassment on a Likert Scale (1 lowest-5 highest) | |
|---|---|
| **Theme** | **Count** |
| **Total (n=2105)** | 12.5% rated 1 (lowest score) (263 out of 2105)<br>10% rated 2 (211 out of 2105)<br>12.6% rated 3 (265 out of 2105)<br>21% rated 4 (441 out of 2105)<br>43.9% rated 5 (highest score) (925 out of 2105) |
| **Women Mentee (n=1278)** | 13.1% rated 1 (lowest score) (168 out of 1278)<br>10.1% rated 2 (129 out of 1278)<br>13% rated 3 (166 out of 1278)<br>21.2% rated 4 (271 out of 1278)<br>42.6% rated 5 (highest score) (544 out of 1278) |
| **Men Mentee (n=799)** | 11.4% rated 1 (lowest score) (91 out of 799)<br>9.8% rated 2 (78 out of 799)<br>12% rated 3 (96 out of 799)<br>20.9% rated 4 (167 out of 799)<br>45.9% rated 5 (highest score) (367 out of 799) |
| **Non-binary or did not disclose gender (n=24)** | 12.5% rated 1 (lowest score) (3 out of 24)<br>16.7% rated 2 (4 out of 24)<br>8.3% rated 3 (2 out of 24)<br>12.5% rated 4 (3 out of 24)<br>50% rated 5 (highest score) (12 out of 24) |
| **Did not respond to this survey Question** | 0.4% (9 out of 2114) |

| Table S72. Mentor ensures a working environment free from discrimination and harassment on a Likert Scale (1 lowest-5 highest) | |
|---|---|
| **Theme** | **Count** |
| **Total (n=2105)** | 12.5% rated 1 (lowest score) (263 out of 2105)<br>10% rated 2 (211 out of 2105)<br>12.6% rated 3 (265 out of 2105)<br>21% rated 4 (441 out of 2105)<br>43.9% rated 5 (highest score) (925 out of 2105) |
| **Non-Citizen (international) Mentee (n=803)** | 14.4% rated 1 (lowest score) (116 out of 803)<br>9.8% rated 2 (79 out of 803)<br>12.3% rated 3 (99 out of 803)<br>21.4% rated 4 (172 out of 803)<br>42% rated 5 (highest score) (337 out of 803) |
| **Citizen (national) Mentee (n=1300)** | 11.2% rated 1 (lowest score) (146 out of 1300)<br>10.2% rated 2 (132 out of 1300)<br>12.7% rated 3 (165 out of 1300)<br>20.7% rated 4 (269 out of 1300)<br>45.2% rated 5 (highest score) (588 out of 1300) |
| **Did not respond to this survey Question** | 0.4% (9 out of 2114) |

| **Table S73. Mentor ensures a working environment free from discrimination and harassment** on a Likert Scale (1 lowest-5 highest) | |
|---|---|
| **Theme** | **count** |
| **Non-Citizen (international) Women Mentee (n=440)** | 16.6% rated 1 (lowest score) (73 out of 440)<br>9.3% rated 2 (41 out of 440)<br>11.1% rated 3 (49 out of 440)<br>21.8% rated 4 (96 out of 440)<br>41.1% rated 5 (highest score) (181 out of 440) |
| **Non-Citizen (international) Men Mentee (n=353)** | 11.9% rated 1 (lowest score) (42 out of 353)<br>10.5% rated 2 (37 out of 353)<br>13.9% rated 3 (49 out of 353)<br>21% rated 4 (74 out of 353)<br>42.8% rated 5 (highest score) (151 out of 353) |
| **Citizen (national) Women Mentee (n=838)** | 11.3% rated 1 (lowest score) (95 out of 838)<br>10.5% rated 2 (88 out of 838)<br>14% rated 3 (117 out of 838)<br>20.9% rated 4 (175 out of 838)<br>43.3% rated 5 (highest score) (363 out of 838) |
| **Citizen (national) Men Mentee (n=446)** | 11% rated 1 (lowest score) (49 out of 446)<br>9.2% rated 2 (41 out of 446)<br>10.5% rated 3 (47 out of 446)<br>20.9% rated 4 (93 out of 446)<br>48.4% rated 5 (highest score) (216 out of 446) |

| **Table S74. Mentor accurately estimates your level of scientific knowledge.** on a Likert Scale (1 lowest-5 highest) | |
|---|---|
| **Theme** | **Count** |
| **Total (n=2090)** | 8.1% rated 1 (lowest score) (169 out of 2090)<br>13.5% rated 2 (283 out of 2090)<br>19.4% rated 3 (406 out of 2090)<br>33% rated 4 (690 out of 2090)<br>25.9% rated 5 (highest score) (542 out of 2090) |
| **Women Mentee (n=1267)** | 8.9% rated 1 (lowest score) (113 out of 1267)<br>12.9% rated 2 (164 out of 1267)<br>20.5% rated 3 (260 out of 1267)<br>33.4% rated 4 (423 out of 1267)<br>24.3% rated 5 (highest score) (307 out of 1267) |
| **Men Mentee (n=795)** | 6.7% rated 1 (lowest score) (53 out of 795)<br>14.3% rated 2 (114 out of 795)<br>17.7% rated 3 (141 out of 795)<br>32.6% rated 4 (259 out of 795)<br>28.7% rated 5 (highest score) (228 out of 795) |
| **Non-binary or did not disclose gender (n=24)** | 8.3% rated 1 (lowest score) (2 out of 24)<br>16.7% rated 2 (4 out of 24)<br>12.5% rated 3 (3 out of 24)<br>33.3% rated 4 (8 out of 24) |

| | |
|---|---|
| | 29.2% rated 5 (highest score) (7 out of 24) |
| **Did not respond to this survey Question** | 1.3% (24 out of 2114) |

| **Table S75. Mentor accurately estimates your level of scientific knowledge.** on a Likert Scale (1 lowest-5 highest) | |
|---|---|
| **Theme** | **Count** |
| **Total (n=2090)** | 8.1% rated 1 (lowest score) (169 out of 2090)<br>13.5% rated 2 (283 out of 2090)<br>19.4% rated 3 (406 out of 2090)<br>33% rated 4 (690 out of 2090)<br>25.9% rated 5 (highest score) (542 out of 2090) |
| **Non-Citizen (international) Mentee (n=1292)** | 6.7% rated 1 (lowest score) (86 out of 1292)<br>12.3% rated 2 (159 out of 1292)<br>21% rated 3 (271 out of 1292)<br>33.7% rated 4 (436 out of 1292)<br>26.3% rated 5 (highest score) (340 out of 1292) |
| **Citizen (national) Mentee (n=796)** | 10.4% rated 1 (lowest score) (83 out of 796)<br>15.6% rated 2 (124 out of 796)<br>16.7% rated 3 (133 out of 796)<br>31.9% rated 4 (254 out of 796)<br>25.4% rated 5 (highest score) (202 out of 796) |
| **Did not respond to this survey Question** | 1.3% (24 out of 2114) |

| **Table S76. Mentor accurately estimates your level of scientific knowledge.** on a Likert Scale (1 lowest-5 highest) | |
|---|---|
| **Theme** | **count** |
| **Non-Citizen (international) Women Mentee (n=436)** | 12.4% rated 1 (lowest score) (54 out of 436)<br>15.1% rated 2 (66 out of 436)<br>15.6% rated 3 (68 out of 436)<br>31.4% rated 4 (137 out of 436)<br>25.5% rated 5 (highest score) (111 out of 436) |
| **Non-Citizen (international) Men Mentee (n=352)** | 7.7% rated 1 (lowest score) (27 out of 352)<br>16.2% rated 2 (57 out of 352)<br>18.2% rated 3 (64 out of 352)<br>33% rated 4 (116 out of 352)<br>25% rated 5 (highest score) (88 out of 352) |
| **Citizen (national) Women Mentee (n=831)** | 7.1% rated 1 (lowest score) (59 out of 831)<br>11.8% rated 2 (98 out of 831)<br>23.1% rated 3 (192 out of 831)<br>34.4% rated 4 (286 out of 831)<br>23.6% rated 5 (highest score) (196 out of 831) |
| **Citizen (national) Men Mentee** | 5.9% rated 1 (lowest score) (26 out of 443) |

| | |
|---|---|
| **(n=443)** | 12.9% rated 2 (57 out of 443)<br>17.4% rated 3 (77 out of 443)<br>32.3% rated 4 (143 out of 443)<br>31.6% rated 5 (highest score) (140 out of 443) |

| **Table S77. Mentor accurately estimates mentee ability to conduct research.** on a Likert Scale (1 lowest-5 highest) | |
|---|---|
| **Theme** | **Count** |
| **Total (n=2092)** | 8.1% rated 1 (lowest score) (170 out of 2092)<br>12.2% rated 2 (225 out of 2092)<br>16.8% rated 3 (352 out of 2092)<br>31.9% rated 4 (668 out of 2092)<br>30.9% rated 5 (highest score) (647 out of 2092) |
| **Women Mentee (n=1271)** | 8.8% rated 1 (lowest score) (112 out of 1271)<br>11.1% rated 2 (141 out of 1271)<br>17.3% rated 3 (220 out of 1271)<br>32.5% rated 4 (413 out of 1271)<br>30.3% rated 5 (highest score) (385 out of 1271) |
| **Men Mentee (n=794)** | 7.1% rated 1 (lowest score) (56 out of 794)<br>13.6% rated 2 (108 out of 794)<br>16% rated 3 (127 out of 794)<br>31.6% rated 4 (251 out of 794)<br>31.7% rated 5 (highest score) (252 out of 794) |
| **Non-binary or did not disclose gender (n=23)** | 4.3% rated 1 (lowest score) (1 out of 23)<br>21.7% rated 2 (5 out of 23)<br>13% rated 3 (3 out of 23)<br>17.4% rated 4 (4 out of 23)<br>43.5% rated 5 (highest score) (10 out of 23) |
| **Did not respond to this survey Question** | 1% (22 out of 2114) |

| **Table S78. Mentor accurately estimates mentee ability to conduct research.** on a Likert Scale (1 lowest-5 highest) | |
|---|---|
| **Theme** | **Count** |
| **Total (n=2092)** | 8.1% rated 1 (lowest score) (170 out of 2092)<br>12.2% rated 2 (225 out of 2092)<br>16.8% rated 3 (352 out of 2092)<br>31.9% rated 4 (668 out of 2092)<br>30.9% rated 5 (highest score) (647 out of 2092) |
| **Non-Citizen (international) Mentee (n=796)** | 9.9% rated 1 (lowest score) (79 out of 796)<br>14.1% rated 2 (112 out of 796)<br>15.1% rated 3 (120 out of 796)<br>31.2% rated 4 (248 out of 796)<br>29.8% rated 5 (highest score) (237 out of 796) |
| **Citizen (national) Mentee (n=1294)** | 7% rated 1 (lowest score) (91 out of 1294)<br>11.1% rated 2 (143 out of 1294) |

| | |
|---|---|
| | 17.9% rated 3 (231 out of 1294)<br>32.4% rated 4 (419 out of 1294)<br>31.7% rated 5 (highest score) (410 out of 1294) |
| **Did not respond to this survey Question** | 1% (22 out of 2114) |

| **Table S79. Mentor accurately estimates mentee ability to conduct research.**<br>on a Likert Scale (1 lowest-5 highest) | |
|---|---|
| **Theme** | **count** |
| **Non-Citizen (international) Women Mentee (n=437)** | 11.4% rated 1 (lowest score) (50 out of 437)<br>12.4% rated 2 (54 out of 437)<br>16% rated 3 (70 out of 437)<br>29.7% rated 4 (130 out of 437)<br>30.4% rated 5 (highest score) (133 out of 437) |
| **Non-Citizen (international) Men Mentee (n=351)** | 8% rated 1 (lowest score) (28 out of 351)<br>16% rated 2 (56 out of 351)<br>13.7% rated 3 (48 out of 351)<br>33.3% rated 4 (117 out of 351)<br>29.1% rated 5 (highest score) (102 out of 351) |
| **Citizen (national) Women Mentee (n=834)** | 7.4% rated 1 (lowest score) (62 out of 834)<br>10.4% rated 2 (87 out of 834)<br>18% rated 3 (150 out of 834)<br>33.9% rated 4 (283 out of 834)<br>30.2% rated 5 (highest score) (252 out of 834) |
| **Citizen (national) Men Mentee (n=443)** | 6.3% rated 1 (lowest score) (28 out of 443)<br>11.7% rated 2 (52 out of 443)<br>17.8% rated 3 (79 out of 443)<br>30.2% rated 4 (134 out of 443)<br>33.9% rated 5 (highest score) (150 out of 443) |

| **Table S80. Mentor employs strategies to enhance your understanding of the research.**<br>on a Likert Scale (1 lowest-5 highest) | |
|---|---|
| **Theme** | **Count** |
| **Total (n=2089)** | 14.4% rated 1 (lowest score) (301 out of 2089)<br>18.6% rated 2 (388 out of 2089)<br>20.6% rated 3 (431 out of 2089)<br>25.5% rated 4 (533 out of 2089)<br>20.9% rated 5 (highest score) (436 out of 2089) |
| **Women Mentee (n=1269)** | 14.7% rated 1 (lowest score) (187 out of 1269)<br>17.3% rated 2 (220 out of 1269)<br>21.7% rated 3 (276 out of 1269)<br>25.1% rated 4 (318 out of 1269)<br>21.1% rated 5 (highest score) (268 out of 1269) |
| **Men Mentee (n=792)** | 13.5% rated 1 (lowest score) (107 out of 792)<br>20.6% rated 2 (163 out of 792)<br>18.8% rated 3 (149 out of 792)<br>26.6% rated 4 (211 out of 792) |

| | |
|---|---|
| | 20.5% rated 5 (highest score) (162 out of 792) |
| **Non-binary or did not disclose gender (n=24)** | 25% rated 1 (lowest score) (6 out of 24)<br>12.5% rated 2 (3 out of 24)<br>20.8% rated 3 (5 out of 24)<br>16.7% rated 4 (4 out of 24)<br>25% rated 5 (highest score) (6 out of 24) |
| **Did not respond to this survey Question** | 1.2% (25 out of 2114) |

| **Table S81. Mentor employs strategies to enhance your understanding of the research.** on a Likert Scale (1 lowest-5 highest) | |
|---|---|
| **Theme** | **Count** |
| **Total (n=2091)** | 13.3% rated 1 (lowest score) (301 out of 2091)<br>18.6% rated 2 (388 out of 2091)<br>20% rated 3 (433 out of 2091)<br>26.2% rated 4 (533 out of 2091)<br>21.9% rated 5 (highest score) (436 out of 2091) |
| **Non-Citizen (international) Mentee (n=796)** | 16.2% rated 1 (lowest score) (129 out of 796)<br>18.5% rated 2 (147 out of 796)<br>21.6% rated 3 (172 out of 796)<br>24.5% rated 4 (195 out of 796)<br>19.2% rated 5 (highest score) (153 out of 796) |
| **Citizen (national) Mentee (n=1291)** | 13.3% rated 1 (lowest score) (172 out of 1291)<br>18.6% rated 2 (240 out of 1291)<br>20% rated 3 (258 out of 1291)<br>26.2% rated 4 (338 out of 1291)<br>21.9% rated 5 (highest score) (283 out of 1291) |
| **Did not respond to this survey Question** | 1.1% (23 out of 2114) |

| **Table S82. Mentor employs strategies to enhance your understanding of the research.** on a Likert Scale (1 lowest-5 highest) | |
|---|---|
| **Theme** | **count** |
| **Non-Citizen (international) Women Mentee (n=437)** | 16.9% rated 1 (lowest score) (74 out of 437)<br>15.6% rated 2 (68 out of 437)<br>23.3% rated 3 (102 out of 437)<br>24.3% rated 4 (106 out of 437)<br>19.9% rated 5 (highest score) (87 out of 437) |
| **Non-Citizen (international) Men Mentee (n=351)** | 15.1% rated 1 (lowest score) (53 out of 351)<br>21.9% rated 2 (77 out of 351)<br>19.1% rated 3 (67 out of 351)<br>25.1% rated 4 (88 out of 351)<br>18.8% rated 5 (highest score) (66 out of 351) |
| **Citizen (national) Women Mentee (n=832)** | 13.6% rated 1 (lowest score) (113 out of 832)<br>18.3% rated 2 (152 out of 832)<br>20.9% rated 3 (174 out of 832)<br>25.5% rated 4 (212 out of 832) |

| | |
|---|---|
| | 21.8% rated 5 (highest score) (181 out of 832) |
| **Citizen (national) Men Mentee (n=441)** | 12.2% rated 1 (lowest score) (54 out of 441)<br>19.5% rated 2 (86 out of 441)<br>18.6% rated 3 (82 out of 441)<br>27.9% rated 4 (123 out of 441)<br>21.8% rated 5 (highest score) (96 out of 441) |

| **Table S83. Mentor motivates the mentee to build confidence in themselves as a scientist.** on a Likert Scale (1 lowest-5 highest) | |
|---|---|
| **Theme** | **Count** |
| **Total (n=2091)** | 20.1% rated 1 (lowest score) (420 out of 2091)<br>14.8% rated 2 (310 out of 2091)<br>15.4% rated 3 (323 out of 2091)<br>19.9% rated 4 (417 out of 2091)<br>29.7% rated 5 (highest score) (621 out of 2091) |
| **Women Mentee (n=1269)** | 20.8% rated 1 (lowest score) (264 out of 1269)<br>14.7% rated 2 (186 out of 1269)<br>14.9% rated 3 (189 out of 1269)<br>19.9% rated 4 (252 out of 1269)<br>29.8% rated 5 (highest score) (378 out of 1269) |
| **Men Mentee (n=795)** | 18.6% rated 1 (lowest score) (148 out of 795)<br>155% rated 2 (119 out of 795)<br>16.4% rated 3 (130 out of 795)<br>20.4% rated 4 (162 out of 795)<br>29.7% rated 5 (highest score) (236 out of 795) |
| **Non-binary or did not disclose gender (n=23)** | 30.4% rated 1 (lowest score) (7 out of 23)<br>13% rated 2 (3 out of 23)<br>13% rated 3 (3 out of 23)<br>13% rated 4 (3 out of 23)<br>30.4% rated 5 (highest score) (7 out of 23) |
| **Did not respond to this survey Question** | 1.1% (23 out of 2114) |

| **Table S84. Mentor motivates the mentee to build confidence in themselves as a scientist.** on a Likert Scale (1 lowest-5 highest) | |
|---|---|
| **Theme** | **Count** |
| **Total (n=2091)** | 20.1% rated 1 (lowest score) (420 out of 2092)<br>14.8% rated 2 (310 out of 2092)<br>15.4% rated 3 (323 out of 2092)<br>19.9% rated 4 (417 out of 2092)<br>29.7% rated 5 (highest score) (622 out of 2092) |
| **Non-Citizen (international) Mentee (n=797)** | 23.6% rated 1 (lowest score) (188 out of 797)<br>15.4% rated 2 (123 out of 797)<br>14.6% rated 3 (116 out of 797)<br>19.1% rated 4 (152 out of 797)<br>27.4% rated 5 (highest score) (218 out of 797) |

| | |
|---|---|
| **Citizen (national) Mentee (n=1293)** | 17.9% rated 1 (lowest score) (231 out of 1293)<br>14.5% rated 2 (187 out of 1293)<br>15.9% rated 3 (206 out of 1293)<br>20.5% rated 4 (265 out of 1293)<br>31.2% rated 5 (highest score) (404 out of 1293) |
| **Did not respond to this survey Question** | 1.1% (22 out of 2114) |

| **Table S85. Mentor motivates the mentee to build confidence in themselves as a scientist.**<br>on a Likert Scale (1 lowest-5 highest) | |
|---|---|
| **Theme** | **count** |
| **Non-Citizen (international) Women Mentee (n=438)** | 24.9% rated 1 (lowest score) (109 out of 438)<br>14.4% rated 2 (63 out of 438)<br>13.2% rated 3 (58 out of 438)<br>20.8% rated 4 (91 out of 438)<br>26.7% rated 5 (highest score) (117 out of 438) |
| **Non-Citizen (international) Men Mentee (n=351)** | 21.7% rated 1 (lowest score) (76 out of 351)<br>16.5% rated 2 (58 out of 351)<br>16% rated 3 (56 out of 351)<br>17.4% rated 4 (61 out of 351)<br>28.5% rated 5 (highest score) (100 out of 351) |
| **Citizen (national) Women Mentee (n=831)** | 18.7% rated 1 (lowest score) (155 out of 831)<br>14.8% rated 2 (123 out of 831)<br>15.8% rated 3 (131 out of 831)<br>19.4% rated 4 (161 out of 831)<br>31.4% rated 5 (highest score) (261 out of 831) |
| **Citizen (national) Men Mentee (n=444)** | 16.2% rated 1 (lowest score) (72 out of 444)<br>13.7% rated 2 (61 out of 444)<br>16.7% rated 3 (74 out of 444)<br>22.7% rated 4 (101 out of 444)<br>30.6% rated 5 (highest score) (136 out of 444) |

| **Table S86. Mentor stimulates mentee creativity.**<br>on a Likert Scale (1 lowest-5 highest) | |
|---|---|
| **Theme** | **Count** |
| **Total (n=2092)** | 16.6% rated 1 (lowest score) (347 out of 2092)<br>17.1% rated 2 (358 out of 2092)<br>18.8% rated 3 (394 out of 2092)<br>22.8% rated 4 (476 out of 2092)<br>24.7% rated 5 (highest score) (517 out of 2092) |
| **Women Mentee (n=1270)** | 17% rated 1 (lowest score) (216 out of 1270)<br>16.1% rated 2 (205 out of 1270)<br>19.8% rated 3 (252 out of 1270)<br>22.9% rated 4 (291 out of 1270)<br>24.1% rated 5 (highest score) (306 out of 1270) |
| **Men Mentee (n=794)** | 15.4% rated 1 (lowest score) (122 out of 794)<br>18.5% rated 2 (147 out of 794) |

| | |
|---|---|
| | 17.5% rated 3 (139 out of 794)<br>22.8% rated 4 (181 out of 794)<br>25.8% rated 5 (highest score) (205 out of 794) |
| **Non-binary or did not disclose gender (n=24)** | 29.2% rated 1 (lowest score) (7 out of 24)<br>20.8% rated 2 (5 out of 24)<br>8.3% rated 3 (2 out of 24)<br>16.7% rated 4 (4 out of 24)<br>25% rated 5 (highest score) (6 out of 24) |
| **Did not respond to this survey Question** | 1% (22 out of 2114) |

**Table S87. Mentor stimulates mentee creativity.**
on a Likert Scale (1 lowest-5 highest)

| Theme | Count |
|---|---|
| **Total (n=2092)** | 16.6% rated 1 (lowest score) (347 out of 2092)<br>17.1% rated 2 (358 out of 2092)<br>18.8% rated 3 (394 out of 2092)<br>22.8% rated 4 (476 out of 2092)<br>24.7% rated 5 (highest score) (517 out of 2092) |
| **Non-Citizen (international) Mentee (n=796)** | 20% rated 1 (lowest score) (159 out of 796)<br>18.7% rated 2 (149 out of 796)<br>18.8% rated 3 (150 out of 796)<br>19.6% rated 4 (156 out of 796)<br>22.9% rated 5 (highest score) (182 out of 796) |
| **Citizen (national) Mentee (n=1294)** | 14.5% rated 1 (lowest score) (188 out of 1294)<br>16.2% rated 2 (209 out of 1294)<br>18.8% rated 3 (243 out of 1294)<br>24.7% rated 4 (319 out of 1294)<br>25.9% rated 5 (highest score) (335 out of 1294) |
| **Did not respond to this survey Question** | 1% (22 out of 2114) |

**Table S88. Mentor stimulates mentee creativity.**
on a Likert Scale (1 lowest-5 highest)

| Theme | count |
|---|---|
| **Non-Citizen (international) Women Mentee (n=437)** | 21.1% rated 1 (lowest score) (92 out of 437)<br>17.2% rated 2 (75 out of 437)<br>20.4% rated 3 (89 out of 437)<br>18.3% rated 4 (80 out of 437)<br>23.1% rated 5 (highest score) (101 out of 437) |
| **Non-Citizen (international) Men Mentee (n=351)** | 17.9% rated 1 (lowest score) (63 out of 351)<br>20.8% rated 2 (73 out of 351)<br>17.1% rated 3 (60 out of 351)<br>21.4% rated 4 (75 out of 351)<br>22.8% rated 5 (highest score) (80 out of 351) |
| **Citizen (national) Women Mentee (n=833)** | 14.9% rated 1 (lowest score) (124 out of 833)<br>15.6% rated 2 (130 out of 833) |

| | |
|---|---|
| | 19.6% rated 3 (163 out of 833)<br>25.3% rated 4 (211 out of 833)<br>24.6% rated 5 (highest score) (205 out of 833) |
| **Citizen (national) Men Mentee (n=443)** | 13.3% rated 1 (lowest score) (59 out of 443)<br>16.7% rated 2 (74 out of 443)<br>17.8% rated 3 (79 out of 443)<br>23.9% rated 4 (106 out of 443)<br>28.2% rated 5 (highest score) (125 out of 443) |

**Table S89. Mentor acknowledges mentee professional contributions. Analysis by mentee gender.** on a Likert Scale (1 lowest-5 highest)

| Theme | Count |
|---|---|
| **Total (n=2109)** | 13.1% rated 1 (lowest score) (276 out of 2109)<br>11.3% rated 2 (238 out of 2109)<br>15.6% rated 3 (328 out of 2109)<br>24% rated 4 (506 out of 2109)<br>36.1% rated 5 (highest score) (761 out of 2109) |
| **Women Mentee (n=1280)** | 14.4% rated 1 (lowest score) (184 out of 1280)<br>11.6% rated 2 (149 out of 1280)<br>15.4% rated 3 (197 out of 1280)<br>23.2% rated 4 (297 out of 1280)<br>35.4% rated 5 (highest score) (453 out of 1280) |
| **Men Mentee (n=801)** | 11% rated 1 (lowest score) (88 out of 801)<br>10.6% rated 2 (85 out of 801)<br>15.6% rated 3 (125 out of 801)<br>25.1% rated 4 (201 out of 801)<br>37.7% rated 5 (highest score) (302 out of 801) |
| **Non-binary or did not disclose gender (n=24)** | 12.5% rated 1 (lowest score) (3 out of 24)<br>12.5% rated 2 (3 out of 24)<br>16.7% rated 3 (4 out of 24)<br>33.3% rated 4 (8 out of 24)<br>25% rated 5 (highest score) (6 out of 24) |
| **Did not respond to this survey Question** | 0.2% (5 out of 2114) |

**Table S90. Mentor acknowledges mentee professional contributions by mentee citizenship status.** on a Likert Scale (1 lowest-5 highest)

| Theme | Count |
|---|---|
| **Total (n=2109)** | 13.1% rated 1 (lowest score) (276 out of 2109)<br>11.3% rated 2 (238 out of 2109)<br>15.6% rated 3 (328 out of 2109)<br>24% rated 4 (506 out of 2109)<br>36.1% rated 5 (highest score) (761 out of 2109) |
| **Non-Citizen (international) Mentee (n=807)** | 15.5% rated 1 (lowest score) (125 out of 807)<br>10.4% rated 2 (84 out of 807)<br>16.2% rated 3 (131 out of 807) |

| | |
|---|---|
| | 24.5% rated 4 (198 out of 807)<br>33.3% rated 5 (highest score) (269 out of 807) |
| **Citizen (national) Mentee (n=1300)** | 11.6% rated 1 (lowest score) (151 out of 1300)<br>11.8% rated 2 (153 out of 1300)<br>15.1% rated 3 (196 out of 1300)<br>23.7% rated 4 (308 out of 1300)<br>37.8% rated 5 (highest score) (492 out of 1300) |
| **Did not respond to this survey Question** | 0.2% (5 out of 2114) |

**Table S91. Mentor acknowledges mentee professional contributions by mentee citizenship status.** on a Likert Scale (1 lowest-5 highest)

| Theme | count |
|---|---|
| **Non-Citizen (international) Women Mentee (n=443)** | 17.6% rated 1 (lowest score) (78 out of 443)<br>10.4% rated 2 (46 out of 443)<br>14.2% rated 3 (63 out of 443)<br>23.9% rated 4 (106 out of 443)<br>33.9% rated 5 (highest score) (150 out of 443) |
| **Non-Citizen (international) Men Mentee (n=356)** | 12.9% rated 1 (lowest score) (46 out of 356)<br>10.1% rated 2 (36 out of 356)<br>18.8% rated 3 (67 out of 356)<br>25.3% rated 4 (90 out of 356)<br>32.9% rated 5 (highest score) (117 out of 356) |
| **Citizen (national) Women Mentee (n=837)** | 12.7% rated 1 (lowest score) (106 out of 837)<br>12.3% rated 2 (103 out of 837)<br>16% rated 3 (134 out of 837)<br>22.8% rated 4 (191 out of 837)<br>36.2% rated 5 (highest score) (303 out of 837) |
| **Citizen (national) Men Mentee (n=445)** | 9.4% rated 1 (lowest score) (42 out of 445)<br>11% rated 2 (49 out of 445)<br>13% rated 3 (58 out of 445)<br>24.9% rated 4 (111 out of 445)<br>41.6% rated 5 (highest score) (185 out of 445) |

**Table S92. Mentor support of mentee Professional Development by supporting mentee career Plan? by mentee gender** on a Likert Scale (1 lowest-5 highest)

| Theme | Count |
|---|---|
| **Total (n=2090)** | 13.5% rated 1 (lowest score) (282 out of 2090)<br>12.4% rated 2 (260 out of 2090)<br>17.5% rated 3 (365 out of 2090)<br>22% rated 4 (460 out of 2090)<br>34.6% rated 5 (highest score) (723 out of 2090) |
| **Women Mentee (n=1269)** | 14.9% rated 1 (lowest score) (189 out of 1269)<br>13.1% rated 2 (166 out of 1269)<br>16.4% rated 3 (208 out of 1269) |

| | |
|---|---|
| | 22.9% rated 4 (291 out of 1269)<br>32.7% rated 5 (highest score) (415 out of 1269) |
| **Men Mentee (n=793)** | 11% rated 1 (lowest score) (87 out of 793)<br>11.6% rated 2 (92 out of 793)<br>19.2% rated 3 (152 out of 793)<br>20.4% rated 4 (162 out of 793)<br>37.8% rated 5 (highest score) (300 out of 793) |
| **Non-binary or did not disclose gender (n=24)** | 20.8% rated 1 (lowest score) (5 out of 24)<br>8.3% rated 2 (2 out of 24)<br>8.3% rated 3 (2 out of 24)<br>29.2% rated 4 (7 out of 24)<br>33.3% rated 5 (highest score) (8 out of 24) |
| **Did not respond to this survey Question** | 1.1% (24 out of 2114) |

| **Table S93. Mentor support of mentee Professional Development by supporting mentee career<br>Plan? by mentee citizenship status** on a Likert Scale (1 lowest-5 highest) | |
|---|---|
| **Theme** | **Count** |
| **Total (n=2090)** | 13.5% rated 1 (lowest score) (282 out of 2090)<br>12.4% rated 2 (260 out of 2090)<br>17.5% rated 3 (365 out of 2090)<br>22% rated 4 (460 out of 2090)<br>34.6% rated 5 (highest score) (723 out of 2090) |
| **Non-Citizen (international) Mentee (n=798)** | 15.7% rated 1 (lowest score) (125 out of 798)<br>14.7% rated 2 (117 out of 798)<br>16.2% rated 3 (129 out of 798)<br>21.9% rated 4 (175 out of 798)<br>31.6% rated 5 (highest score) (252 out of 798) |
| **Citizen (national) Mentee (n=1290)** | 12.1% rated 1 (lowest score) (156 out of 1290)<br>11.1% rated 2 (143 out of 1290)<br>18.2% rated 3 (235 out of 1290)<br>22.1% rated 4 (471 out of 1290)<br>36.5% rated 5 (highest score) (out of 1290) |
| **Did not respond to this survey Question** | 1.1% (24 out of 2114) |

| **Table S94. Mentor support of mentee Professional Development by supporting mentee career<br>Plan? by mentee citizenship status** on a Likert Scale (1 lowest-5 highest) | |
|---|---|
| **Theme** | **count** |
| **Non-Citizen (international) Women Mentee (n=438)** | 17.1% rated 1 (lowest score) (75 out of 438)<br>18% rated 2 (79 out of 438)<br>11.4% rated 3 (50 out of 438)<br>24% rated 4 (105 out of 438)<br>29.5% rated 5 (highest score) (129 out of 438) |
| **Non-Citizen (international) Men** | 13.6% rated 1 (lowest score) (48 out of 352) |

| | |
|---|---|
| **Mentee (n=352)** | 10.5% rated 2 (37 out of 352)<br>21.9% rated 3 (77 out of 352)<br>19.6% rated 4 (69 out of 352)<br>34.4% rated 5 (highest score) (121 out of 352) |
| **Citizen (national) Women Mentee (n=831)** | 13.7% rated 1 (lowest score) (114 out of 831)<br>10.5% rated 2 (87 out of 831)<br>19% rated 3 (158 out of 831)<br>22.4% rated 4 (186 out of 831)<br>34.4% rated 5 (highest score) (286 out of 831) |
| **Citizen (national) Men Mentee (n=441)** | 8.8% rated 1 (lowest score) (39 out of 441)<br>12.5% rated 2 (55 out of 441)<br>17% rated 3 (75 out of 441)<br>21.1% rated 4 (93 out of 441)<br>40.6% rated 5 (highest score) (179 out of 441) |

**Table S95. Mentor helps the mentee develop strategies to meet career goals.**
on a Likert Scale (1 lowest-5 highest)

| Theme | Count |
|---|---|
| **Total (n=2091)** | 21.2% rated 1 (lowest score) (443 out of 2091)<br>15.9% rated 2 (333 out of 2091)<br>19.4% rated 3 (406 out of 2091)<br>21.9% rated 4 (457 out of 2091)<br>21.6% rated 5 (highest score) (452 out of 2091) |
| **Women Mentee (n=1270)** | 23.6% rated 1 (lowest score) (300 out of 1270)<br>15.3% rated 2 (194 out of 1270)<br>19.8% rated 3 (251 out of 1270)<br>20.4% rated 4 (259 out of 1270)<br>20.9% rated 5 (highest score) (266 out of 1270) |
| **Men Mentee (n=793)** | 17.2% rated 1 (lowest score) (136 out of 793)<br>16.6% rated 2 (132 out of 793)<br>19.2% rated 3 (152 out of 793)<br>24.2% rated 4 (192 out of 793)<br>22.8% rated 5 (highest score) (181 out of 793) |
| **Non-binary or did not disclose gender (n=24)** | 25% rated 1 (lowest score) (6 out of 24)<br>20.8% rated 2 (5 out of 24)<br>8.3% rated 3 (2 out of 24)<br>25% rated 4 (6 out of 24)<br>20.8% rated 5 (highest score) (5 out of 24) |
| **Did not respond to this survey Question** | 1.1% (23 out of 2114) |

**Table S96. Mentor helps the mentee develop strategies to meet career goals.**
on a Likert Scale (1 lowest-5 highest)

| Theme | Count |
|---|---|
| **Total (n=2091)** | 21.2% rated 1 (lowest score) (443 out of 2091) |

| | |
|---|---|
| | 15.9% rated 2 (333 out of 2091)<br>19.4% rated 3 (406 out of 2091)<br>21.9% rated 4 (457 out of 2091)<br>21.6% rated 5 (highest score) (452 out of 2091) |
| **Non-Citizen (international) Mentee (n=799)** | 23.7% rated 1 (lowest score) (189 out of 799)<br>17.6% rated 2 (141 out of 799)<br>16.6% rated 3 (133 out of 799)<br>22.5% rated 4 (180 out of 799)<br>19.5% rated 5 (highest score) (156 out of 799) |
| **Citizen (national) Mentee (n=1290)** | 19.6% rated 1 (lowest score) (253 out of 1290)<br>14.9% rated 2 (192 out of 1290)<br>21.1% rated 3 (272 out of 1290)<br>21.5% rated 4 (277 out of 1290)<br>22.9% rated 5 (highest score) (296 out of 1290) |
| **Did not respond to this survey Question** | 1.1% (23 out of 2114) |

**Table S97. Mentor helps the mentee develop strategies to meet career goals.**
on a Likert Scale (1 lowest-5 highest)

| Theme | count |
|---|---|
| **Non-Citizen (international) Women Mentee (n=438)** | 29.2% rated 1 (lowest score) (128 out of 438)<br>16% rated 2 (70 out of 438)<br>15.5% rated 3 (68 out of 438)<br>20.1% rated 4 (88 out of 438)<br>19.2% rated 5 (highest score) (84 out of 438) |
| **Non-Citizen (international) Men Mentee (n=353)** | 16.4% rated 1 (lowest score) (58 out of 353)<br>19.3% rated 2 (68 out of 353)<br>18.4% rated 3 (65 out of 353)<br>25.8% rated 4 (91 out of 353)<br>20.1% rated 5 (highest score) (71 out of 353) |
| **Citizen (national) Women Mentee (n=832)** | 20.7% rated 1 (lowest score) (172 out of 832)<br>14.9% rated 2 (124 out of 832)<br>22% rated 3 (183 out of 832)<br>20.6% rated 4 (171 out of 832)<br>21.9% rated 5 (highest score) (182 out of 832) |
| **Citizen (national) Men Mentee (n=440)** | 17.7% rated 1 (lowest score) (78 out of 440)<br>14.5% rated 2 (64 out of 440)<br>19.8% rated 3 (87 out of 440)<br>23% rated 4 (101 out of 440)<br>25% rated 5 (highest score) (110 out of 440) |

**Table S98. Mentor negotiates a path to professional independence with mentee.**
on a Likert Scale (1 lowest-5 highest)

| Theme | Count |
|---|---|
| **Total (n=2082)** | 21.4% rated 1 (lowest score) (445 out of 2082) |

| | |
|---|---|
| | 16.2% rated 2 (338 out of 2082)<br>21% rated 3 (437 out of 2082)<br>21.8% rated 4 (453 out of 2082)<br>19.6% rated 5 (highest score) (409 out of 2082) |
| **Women Mentee (n=1261)** | 24% rated 1 (lowest score) (296 out of 1261)<br>15.9% rated 2 (196 out of 1261)<br>21.5% rated 3 (265 out of 1261)<br>21.5% rated 4 (265 out of 1261)<br>19.4% rated 5 (highest score) (239 out of 1261) |
| **Men Mentee (n=793)** | 17.7% rated 1 (lowest score) (140 out of 793)<br>17.4% rated 2 (138 out of 793)<br>21.1% rated 3 (167 out of 793)<br>23.2% rated 4 (184 out of 793)<br>20.7% rated 5 (highest score) (164 out of 793) |
| **Non-binary or did not disclose gender (n=24)** | 29.2% rated 1 (lowest score) (7 out of 24)<br>12.5% rated 2 (3 out of 24)<br>16.7% rated 3 (4 out of 24)<br>16.7% rated 4 (4 out of 24)<br>25% rated 5 (highest score) (6 out of 24) |
| **Did not respond to this survey Question** | 1.5% (32 out of 2114) |

**Table S99. Mentor negotiates a path to professional independence with mentee.**
on a Likert Scale (1 lowest-5 highest)

| Theme | Count |
|---|---|
| **Total (n=2082)** | 21.4% rated 1 (lowest score) (445 out of 2082)<br>16.2% rated 2 (338 out of 2082)<br>21% rated 3 (437 out of 2082)<br>21.8% rated 4 (453 out of 2082)<br>19.6% rated 5 (highest score) (409 out of 2082) |
| **Non-Citizen (international) Mentee (n=795)** | 24.3% rated 1 (lowest score) (193 out of 795)<br>16.9% rated 2 (134 out of 795)<br>19.7% rated 3 (157 out of 795)<br>20.9% rated 4 (166 out of 795)<br>18.2% rated 5 (highest score) (145 out of 795) |
| **Citizen (national) Mentee (n=1285)** | 19.5% rated 1 (lowest score) (251 out of 1285)<br>15.9% rated 2 (204 out of 1285)<br>21.7% rated 3 (279 out of 1285)<br>22.3% rated 4 (287 out of 1285)<br>20.5% rated 5 (highest score) (264 out of 1285) |
| **Did not respond to this survey Question** | 1.5% (32 out of 2114) |

**Table S100. Mentor negotiates a path to professional independence score with mentee.**
on a Likert Scale (1 lowest-5 highest)

| Theme | count |
|---|---|
| **Non-Citizen (international) Women** | 28.5% rated 1 (lowest score) (124 out of 435) |

| | |
|---|---|
| **Mentee (n=435)** | 16.3% rated 2 (71 out of 435)<br>18.4% rated 3 (80 out of 435)<br>18.6% rated 4 (81 out of 435)<br>18.2% rated 5 (highest score) (79 out of 435) |
| **Non-Citizen (international) Men Mentee (n=332)** | 19.6% rated 1 (lowest score) (65 out of 332)<br>18.7% rated 2 (62 out of 332)<br>23.2% rated 3 (77 out of 332)<br>19% rated 4 (63 out of 332)<br>19.6% rated 5 (highest score) (65 out of 332) |
| **Citizen (national) Women Mentee (n=826)** | 20.8% rated 1 (lowest score) (172 out of 826)<br>15.1% rated 2 (125 out of 826)<br>22.4% rated 3 (185 out of 826)<br>22.3% rated 4 (184 out of 826)<br>19.4% rated 5 (highest score) (160 out of 826) |
| **Citizen (national) Men Mentee (n=430)** | 17.4% rated 1 (lowest score) (75 out of 430)<br>17.7% rated 2 (76 out of 430)<br>20.9% rated 3 (90 out of 430)<br>23.5% rated 4 (101 out of 430)<br>20.5% rated 5 (highest score) (88 out of 430) |

**Table S101. Mentor takes into account the biases and prejudices they bring to the mentoring relationship.** on a Likert Scale (1 lowest-5 highest)

| Theme | Count |
|---|---|
| **Total (n=2097)** | 20.9% rated 1 (lowest score) (439 out of 2097)<br>18.8% rated 2 (395 out of 2097)<br>27.4% rated 3 (575 out of 2097)<br>18.8% rated 4 (395 out of 2097)<br>14% rated 5 (highest score) (293 out of 2097) |
| **Women Mentee (n=1272)** | 23% rated 1 (lowest score) (292 out of 1272)<br>19.3% rated 2 (245 out of 1272)<br>27.3% rated 3 (347 out of 1272)<br>17% rated 4 (216 out of 1272)<br>13.5% rated 5 (highest score) (172 out of 1272) |
| **Men Mentee (n=797)** | 17.3% rated 1 (lowest score) (138 out of 797)<br>17.9% rated 2 (143 out of 797)<br>28% rated 3 (223 out of 797)<br>22.2% rated 4 (177 out of 797)<br>14.6% rated 5 (highest score) (116 out of 797) |
| **Non-binary or did not disclose gender (n=24)** | 29.2% rated 1 (lowest score) (7 out of 24)<br>25% rated 2 (6 out of 24)<br>16.7% rated 3 (4 out of 24)<br>8.3% rated 4 (2 out of 24)<br>20.8% rated 5 (highest score) (5 out of 24) |
| **Did not respond to this survey Question** | 0.8% (17 out of 2114) |

| **Table S102. Mentor takes into account the biases and prejudices they bring to the mentoring relationship.** on a Likert Scale (1 lowest-5 highest) | |
|---|---|
| **Theme** | **Count** |
| **Total (n=2097)** | 20.9% rated 1 (lowest score) (439 out of 2097)<br>18.8% rated 2 (395 out of 2097)<br>27.4% rated 3 (575 out of 2097)<br>18.8% rated 4 (395 out of 2097)<br>14% rated 5 (highest score) (293 out of 2097) |
| **Non-Citizen (international) Mentee (n=804)** | 22.1% rated 1 (lowest score) (178 out of 804)<br>18.9% rated 2 (152 out of 804)<br>27.9% rated 3 (224 out of 804)<br>16.8% rated 4 (135 out of 804)<br>14.3% rated 5 (highest score) (115 out of 804) |
| **Citizen (national) Mentee (n=1291)** | 20.1% rated 1 (lowest score) (260 out of 1291)<br>18.8% rated 2 (243 out of 1291)<br>27.1% rated 3 (350 out of 1291)<br>20.1% rated 4 (260 out of 1291)<br>13.8% rated 5 (highest score) (178 out of 1291) |
| **Did not respond to this survey Question** | 0.8% (17 out of 2114) |

| **Table S103. Mentor takes into account the biases and prejudices they bring to the mentoring relationship.** on a Likert Scale (1 lowest-5 highest) | |
|---|---|
| **Theme** | **count** |
| **Non-Citizen (international) Women Mentee (n=442)** | 23.1% rated 1 (lowest score) (102 out of 442)<br>19.5% rated 2 (86 out of 442)<br>28.5% rated 3 (126 out of 442)<br>13.6% rated 4 (60 out of 442)<br>15.4% rated 5 (highest score) (68 out of 442) |
| **Non-Citizen (international) Men Mentee (n=354)** | 20.9% rated 1 (lowest score) (74 out of 354)<br>17.2% rated 2 (61 out of 354)<br>27.7% rated 3 (98 out of 354)<br>20.9% rated 4 (74 out of 354)<br>13.3% rated 5 (highest score) (47 out of 354) |
| **Citizen (national) Women Mentee (n=830)** | 22.9% rated 1 (lowest score) (190 out of 830)<br>19.2% rated 2 (159 out of 830)<br>26.6% rated 3 (221 out of 830)<br>18.8% rated 4 (156 out of 830)<br>12.5% rated 5 (highest score) (104 out of 830) |
| **Citizen (national) Men Mentee (n=443)** | 14.4% rated 1 (lowest score) (64 out of 443)<br>18.5% rated 2 (82 out of 443)<br>28.2% rated 3 (125 out of 443)<br>23.3% rated 4 (103 out of 443)<br>15.6% rated 5 (highest score) (69 out of 443) |

| **Table S104. Mentor works respectfully and effectively with mentees whose personal background is different from his/her own (age, race, gender, class, region, nationality, culture, religion, family composition etc.)** on a Likert Scale (1 lowest-5 highest) | |
|---|---|
| **Theme** | **Count** |
| **Total (n=2102)** | 10.8% rated 1 (lowest score) (227 out of 2102)<br>11% rated 2 (231 out of 2102)<br>18.1% rated 3 (381 out of 2102)<br>23.3% rated 4 (490 out of 2102)<br>36.8% rated 5 (highest score) (773 out of 2102) |
| **Women Mentee (n=1277)** | 10.9% rated 1 (lowest score) (139 out of 1277)<br>11.6% rated 2 (148 out of 1277)<br>18.6% rated 3 (237 out of 1277)<br>22.8% rated 4 (291 out of 1277)<br>36.2% rated 5 (highest score) (462 out of 1277) |
| **Men Mentee (n=797)** | 10.5% rated 1 (lowest score) (84 out of 797)<br>10% rated 2 (80 out of 797)<br>17.2% rated 3 (137 out of 797)<br>24.1% rated 4 (192 out of 797)<br>38.1% rated 5 (highest score) (304 out of 797) |
| **Non-binary or did not disclose gender (n=24)** | 12.5% rated 1 (lowest score) (3 out of 24)<br>8.3% rated 2 (2 out of 24)<br>25% rated 3 (6 out of 24)<br>25% rated 4 (6 out of 24)<br>29.2% rated 5 (highest score) (7 out of 24) |
| **Did not respond to this survey Question** | 0.6% (12 out of 2114) |

| **Table S105. Mentor works respectfully and effectively with mentees whose personal background is different from his/her own (age, race, gender, class, region, nationality, culture, religion, family composition etc.)** on a Likert Scale (1 lowest-5 highest) | |
|---|---|
| **Theme** | **Count** |
| **Total (n=2102)** | 10.8% rated 1 (lowest score) (227 out of 2102)<br>11% rated 2 (231 out of 2102)<br>18.1% rated 3 (381 out of 2102)<br>23.3% rated 4 (490 out of 2102)<br>36.8% rated 5 (highest score) (773 out of 2102) |
| **Non-Citizen (international) Mentee (n=803)** | 13.9% rated 1 (lowest score) (112 out of 803)<br>11.3% rated 2 (91 out of 803)<br>17.7% rated 3 (142 out of 803)<br>23.7% rated 4 (190 out of 803)<br>33.4% rated 5 (highest score) (268 out of 803) |
| **Citizen (national) Mentee (n=1297)** | 8.9% rated 1 (lowest score) (115 out of 1297)<br>10.7% rated 2 (139 out of 1297)<br>18.4% rated 3 (238 out of 1297)<br>23.1% rated 4 (300 out of 1297)<br>38.9% rated 5 (highest score) (505 out of 1297) |
| **Did not respond to this survey Question** | 0.6% (12 out of 2114) |

| Table S106. Mentor works respectfully and effectively with mentees whose personal background is different from his/her own (age, race, gender, class, region, nationality, culture, religion, family composition etc.) on a Likert Scale (1 lowest-5 highest) | |
|---|---|
| **Theme** | **count** |
| **Non-Citizen (international) Women Mentee (n=442)** | 14.9% rated 1 (lowest score) (66 out of 442)<br>10.9% rated 2 (48 out of 442)<br>17.4% rated 3 (77 out of 442)<br>21.9% rated 4 (97 out of 442)<br>34.8% rated 5 (highest score) (154 out of 442) |
| **Non-Citizen (international) Men Mentee (n=353)** | 12.7% rated 1 (lowest score) (45 out of 353)<br>11.9% rated 2 (42 out of 353)<br>17.8% rated 3 (63 out of 353)<br>25.2% rated 4 (89 out of 353)<br>32.3% rated 5 (highest score) (114 out of 353) |
| **Citizen (national) Women Mentee (n=835)** | 8.7% rated 1 (lowest score) (73 out of 835)<br>12% rated 2 (100 out of 835)<br>19.2% rated 3 (160 out of 835)<br>23.2% rated 4 (194 out of 835)<br>36.9% rated 5 (highest score) (308 out of 835) |
| **Citizen (national) Men Mentee (n=444)** | 8.8% rated 1 (lowest score) (39 out of 444)<br>8.6% rated 2 (38 out of 444)<br>16.7% rated 3 (74 out of 444)<br>23.2% rated 4 (103 out of 444)<br>42.8% rated 5 (highest score) (190 out of 444) |

| Table S107. Mentor helps mentee network effectively with other scientists via collaborations or meetings? on a Likert Scale (1 lowest-5 highest) | |
|---|---|
| **Theme** | **Count** |
| **Total (n=2099)** | 17.6% rated 1 (lowest score) (370 out of 2099)<br>14.4% rated 2 (303 out of 2099)<br>16.3% rated 3 (343 out of 2099)<br>23.6% rated 4 (495 out of 2099)<br>28% rated 5 (highest score) (588 out of 2099) |
| **Women Mentee (n=1274)** | 18.8% rated 1 (lowest score) (239 out of 1274)<br>15.4% rated 2 (196 out of 1274)<br>14.5% rated 3 (185 out of 1274)<br>24.3% rated 4 (310 out of 1274)<br>27% rated 5 (highest score) (344 out of 1274) |
| **Men Mentee (n=797)** | 15.1% rated 1 (lowest score) (120 out of 797)<br>13.2% rated 2 (105 out of 797)<br>19.1% rated 3 (152 out of 797)<br>22.8% rated 4 (182 out of 797)<br>29.9% rated 5 (highest score) (238 out of 797) |
| **Non-binary or did not disclose gender (n=24)** | 33.3% rated 1 (lowest score) (8 out of 24)<br>8.3% rated 2 (2 out of 24)<br>20.8% rated 3 (5 out of 24)<br>12.5% rated 4 (3 out of 24) |

| | |
|---|---|
| | 25% rated 5 (highest score) (6 out of 24) |
| **Did not respond to this survey Question** | 0.7% (15 out of 2114) |

| **Table S108. Mentor helps mentee network effectively with other scientists via collaborations or meetings?** on a Likert Scale (1 lowest-5 highest) | |
|---|---|
| **Theme** | **Count** |
| **Total (n=2099)** | 17.6% rated 1 (lowest score) (370 out of 2099)<br>14.4% rated 2 (303 out of 2099)<br>16.3% rated 3 (343 out of 2099)<br>23.6% rated 4 (495 out of 2099)<br>28% rated 5 (highest score) (588 out of 2099) |
| **Non-Citizen (international) Mentee (n=803)** | 20.3% rated 1 (lowest score) (163 out of 803)<br>13.4% rated 2 (108 out of 803)<br>17.1% rated 3 (137 out of 803)<br>23.7% rated 4 (190 out of 803)<br>25.5% rated 5 (highest score) (205 out of 803) |
| **Citizen (national) Mentee (n=1294)** | 16% rated 1 (lowest score) (207 out of 1294)<br>15.1% rated 2 (195 out of 1294)<br>15.8% rated 3 (205 out of 1294)<br>23.5% rated 4 (304 out of 1294)<br>29.6% rated 5 (highest score) (383 out of 1294) |
| **Did not respond to this survey Question** | 0.7% (15 out of 2114) |

| **Table S109. Mentor helps mentee network effectively with other scientists via collaborations or meetings?** on a Likert Scale (1 lowest-5 highest) | |
|---|---|
| **Theme** | **count** |
| **Non-Citizen (international) Women Mentee (n=440)** | 21.4% rated 1 (lowest score) (94 out of 440)<br>15% rated 2 (66 out of 440)<br>15.9% rated 3 (70 out of 440)<br>22.7% rated 4 (100 out of 440)<br>25% rated 5 (highest score) (110 out of 440) |
| **Non-Citizen (international) Men Mentee (n=355)** | 18.6% rated 1 (lowest score) (66 out of 355)<br>11.8% rated 2 (42 out of 355)<br>17.5% rated 3 (62 out of 355)<br>25.4% rated 4 (90 out of 355)<br>26.8% rated 5 (highest score) (95 out of 355) |
| **Citizen (national) Women Mentee (n=834)** | 17.4% rated 1 (lowest score) (145 out of 834)<br>15.6% rated 2 (130 out of 834)<br>13.8% rated 3 (115 out of 834)<br>25.2% rated 4 (210 out of 834)<br>28.1% rated 5 (highest score) (234 out of 834) |
| **Citizen (national) Men Mentee (n=442)** | 12.2% rated 1 (lowest score) (54 out of 442)<br>14.3% rated 2 (63 out of 442)<br>20.4% rated 3 (90 out of 442) |

| | |
|---|---|
| | 20.8% rated 4 (92 out of 442)<br>32.4% rated 5 (highest score) (143 out of 442) |

| **Table S110. Mentor helps mentee balance work with your personal life (for example: take family and/or vacation time off).** on a Likert Scale (1 lowest-5 highest) | |
|---|---|
| **Theme** | **Count** |
| **Total (n=2110)** | 14.5% rated 1 (lowest score) (305 out of 2110)<br>12% rated 2 (254 out of 2110)<br>16.6% rated 3 (351 out of 2110)<br>23.3% rated 4 (491 out of 2110)<br>33.6% rated 5 (highest score) (709 out of 2110) |
| **Women Mentee (n=1281)** | 15.3% rated 1 (lowest score) (196 out of 1281)<br>12.1% rated 2 (155 out of 1281)<br>16.2% rated 3 (208 out of 1281)<br>22.8% rated 4 (292 out of 1281)<br>33.6% rated 5 (highest score) (430 out of 1281) |
| **Men Mentee (n=801)** | 12.4% rated 1 (lowest score) (99 out of 801)<br>12.1% rated 2 (97 out of 801)<br>17.5% rated 3 (140 out of 801)<br>24.3% rated 4 (195 out of 801)<br>33.7% rated 5 (highest score) (270 out of 801) |
| **Non-binary or did not disclose gender (n=24)** | 37.5% rated 1 (lowest score) (9 out of 24)<br>4.2% rated 2 (1 out of 24)<br>4.2% rated 3 (1 out of 24)<br>16.7% rated 4 (4 out of 24)<br>37.5% rated 5 (highest score) (9 out of 24) |
| **Did not respond to this survey Question** | 0.2% (4 out of 2114) |

| **Table S111. Mentor helps mentee balance work with your personal life (for example: take family and/or vacation time off).** on a Likert Scale (1 lowest-5 highest) | |
|---|---|
| **Theme** | **Count** |
| **Total (n=2110)** | 14.5% rated 1 (lowest score) (305 out of 2110)<br>12% rated 2 (254 out of 2110)<br>16.6% rated 3 (351 out of 2110)<br>23.3% rated 4 (491 out of 2110)<br>33.6% rated 5 (highest score) (709 out of 2110) |
| **Non-Citizen (international) Mentee (n=806)** | 15.8% rated 1 (lowest score) (127 out of 806)<br>11.9% rated 2 (96 out of 806)<br>16.6% rated 3 (134 out of 806)<br>22.5% rated 4 (181 out of 806)<br>33.3% rated 5 (highest score) (268 out of 806) |
| **Citizen (national) Mentee (n=1302)** | 13.6% rated 1 (lowest score) (177 out of 1302)<br>12.1% rated 2 (158 out of 1302)<br>16.6% rated 3 (216 out of 1302) |

| | |
|---|---|
| | 23.8% rated 4 (310 out of 1302)<br>33.9% rated 5 (highest score) (441 out of 1302) |
| **Did not respond to this survey Question** | 0.2% (4 out of 2114) |

| **Table S112. Mentor helps mentee balance work with your personal life (for example: take family and/or vacation time off).** on a Likert Scale (1 lowest-5 highest) | |
|---|---|
| **Theme** | **count** |
| **Non-Citizen (international) Women Mentee (n=442)** | 16.3% rated 1 (lowest score) (72 out of 442)<br>12.4% rated 2 (55 out of 442)<br>16.5% rated 3 (73 out of 442)<br>22.2% rated 4 (98 out of 442)<br>32.6% rated 5 (highest score) (144 out of 442) |
| **Non-Citizen (international) Men Mentee (n=356)** | 14% rated 1 (lowest score) (50 out of 356)<br>11.5% rated 2 (41 out of 356)<br>16.9% rated 3 (60 out of 356)<br>23.3% rated 4 (83 out of 356)<br>34.4% rated 5 (highest score) (122 out of 356) |
| **Citizen (national) Women Mentee (n=839)** | 14.8% rated 1 (lowest score) (124 out of 839)<br>11.9% rated 2 (100 out of 839)<br>16.1% rated 3 (135 out of 839)<br>23.1% rated 4 (194 out of 839)<br>34.1% rated 5 (highest score) (286 out of 839) |
| **Citizen (national) Men Mentee (n=445)** | 11% rated 1 (lowest score) (49 out of 445)<br>12.6% rated 2 (56 out of 445)<br>18% rated 3 (80 out of 445)<br>25.2% rated 4 (112 out of 445)<br>33.3% rated 5 (highest score) (148 out of 445) |

| **Table S113. Mentor understands his/her impact as a role model for the mentee.** on a Likert Scale (1 lowest-5 highest) | |
|---|---|
| **Theme** | **Count** |
| **Total (n=2084)** | 16.8% rated 1 (lowest score) (350 out of 2084)<br>15.5% rated 2 (322 out of 2084)<br>21% rated 3 (438 out of 2084)<br>23.4% rated 4 (488 out of 2084)<br>23.3% rated 5 (highest score) (486 out of 2084) |
| **Women Mentee (n=1262)** | 16.6% rated 1 (lowest score) (210 out of 1262)<br>15.7% rated 2 (198 out of 1262)<br>21.1% rated 3 (266 out of 1262)<br>27.7% rated 4 (287 out of 1262)<br>23.9% rated 5 (highest score) (301 out of 1262) |
| **Men Mentee (n=794)** | 16.5% rated 1 (lowest score) (131 out of 794)<br>15.2% rated 2 (121 out of 794)<br>21.2% rated 3 (168 out of 794) |

| | |
|---|---|
| | 24.8% rated 4 (197 out of 794)<br>22.3% rated 5 (highest score) (177 out of 794) |
| **Non-binary or did not disclose gender (n=24)** | 33.3% rated 1 (lowest score) (8 out of 24)<br>4.2% rated 2 (1 out of 24)<br>12.5% rated 3 (3 out of 24)<br>16.7% rated 4 (4 out of 24)<br>33.3% rated 5 (highest score) (8 out of 24) |
| **Did not respond to this survey Question** | 1.4% (30 out of 2114) |

| **Table S114. Mentor understands his/her impact as a role model for the mentee.** Analysis by citizenship status. on a Likert Scale (1 lowest-5 highest) | |
|---|---|
| **Theme** | **Count** |
| **Total (n=2084)** | 16.8% rated 1 (lowest score) (350 out of 2084)<br>15.5% rated 2 (322 out of 2084)<br>21% rated 3 (438 out of 2084)<br>23.4% rated 4 (488 out of 2084)<br>23.3% rated 5 (highest score) (486 out of 2084) |
| **Non-Citizen (international) Mentee (n=793)** | 19.8% rated 1 (lowest score) (157 out of 793)<br>16.8% rated 2 (133 out of 793)<br>21.3% rated 3 (169 out of 793)<br>21.6% rated 4 (171 out of 793)<br>20.6% rated 5 (highest score) (163 out of 793) |
| **Citizen (national) Mentee (n=1289)** | 15% rated 1 (lowest score) (193 out of 1289)<br>14.7% rated 2 (189 out of 1289)<br>20.7% rated 3 (267 out of 1289)<br>24.6% rated 4 (317 out of 1289)<br>25.1% rated 5 (highest score) (323 out of 1289) |
| **Did not respond to this survey Question** | 1.4% (30 out of 2114) |

| **Table S115. Mentor understands his/her impact as a role model for the mentee.** Analysis by citizenship status. on a Likert Scale (1 lowest-5 highest) | |
|---|---|
| **Theme** | **count** |
| **Non-Citizen (international) Women Mentee (n=434)** | 20% rated 1 (lowest score) (87 out of 434)<br>16.8% rated 2 (73 out of 434)<br>21% rated 3 (91 out of 434)<br>20.5% rated 4 (89 out of 434)<br>21.7% rated 5 (highest score) (94 out of 434) |
| **Non-Citizen (international) Men Mentee (n=351)** | 18.8% rated 1 (lowest score) (66 out of 351)<br>16.8% rated 2 (59 out of 351)<br>21.7% rated 3 (76 out of 351)<br>23.4% rated 4 (82 out of 351)<br>19.4% rated 5 (highest score) (68 out of 351) |
| **Citizen (national) Women Mentee (n=828)** | 14.9% rated 1 (lowest score) (123 out of 828)<br>15.1% rated 2 (125 out of 828)<br>21.1% rated 3 (175 out of 828) |

| | |
|---|---|
| | 23.9% rated 4 (198 out of 828)<br>25% rated 5 (highest score) (207 out of 828) |
| **Citizen (national) Men Mentee (n=443)** | 14.7% rated 1 (lowest score) (65 out of 443)<br>14% rated 2 (62 out of 443)<br>20.8% rated 3 (92 out of 443)<br>26% rated 4 (115 out of 443)<br>24.6% rated 5 (highest score) (109 out of 443) |

| **Table S116. Mentor helps mentee acquire resources (e.g., grants, fellowships, etc.)** on a Likert Scale (1 lowest-5 highest) | |
|---|---|
| **Theme** | **Count** |
| **Total (n=2102)** | 14.9% rated 1 (lowest score) (314 out of 2102)<br>11.8% rated 2 (248 out of 2102)<br>16.5% rated 3 (347 out of 2102)<br>23.3% rated 4 (489 out of 2102)<br>33.5% rated 5 (highest score) (704 out of 2102) |
| **Women Mentee (n=1275)** | 14.4% rated 1 (lowest score) (184 out of 1275)<br>12.5% rated 2 (159 out of 1275)<br>16.9% rated 3 (215 out of 1275)<br>23.1% rated 4 (295 out of 1275)<br>33.1% rated 5 (highest score) (422 out of 1275) |
| **Men Mentee (n=799)** | 15.4% rated 1 (lowest score) (123 out of 799)<br>10.8% rated 2 (86 out of 799)<br>16.1% rated 3 (129 out of 799)<br>23.8% rated 4 (190 out of 799)<br>33.9% rated 5 (highest score) (271 out of 799) |
| **Non-binary or did not disclose gender (n=24)** | 21.7% rated 1 (lowest score) (5 out of 24)<br>13% rated 2 (3 out of 24)<br>4.3% rated 3 (1 out of 24)<br>17.4% rated 4 (4 out of 24)<br>43.5% rated 5 (highest score) (10 out of 24) |
| **Did not respond to this survey Question** | 0.6% (12 out of 2114) |

| **Table S117. Mentor helps mentee acquire resources (e.g., grants, fellowships, etc.)** Analysis by citizenship status. on a Likert Scale (1 lowest-5 highest) | |
|---|---|
| **Theme** | **Count** |
| **Total (n=2102)** | 14.9% rated 1 (lowest score) (314 out of 2102)<br>11.8% rated 2 (248 out of 2102)<br>16.5% rated 3 (347 out of 2102)<br>23.3% rated 4 (489 out of 2102)<br>33.5% rated 5 (highest score) (704 out of 2102) |
| **Non-Citizen (international) Mentee (n=804)** | 18.4% rated 1 (lowest score) (148 out of 804)<br>11.4% rated 2 (92 out of 804)<br>17.5% rated 3 (141 out of 804)<br>22.9% rated 4 (184 out of 804) |

| | |
|---|---|
| | 29.7% rated 5 (highest score) (239 out of 804) |
| **Citizen (national) Mentee (n=1296)** | 12.8% rated 1 (lowest score) (166 out of 1296)<br>12% rated 2 (156 out of 1296)<br>15.7% rated 3 (204 out of 1296)<br>23.5% rated 4 (305 out of 1296)<br>35.9% rated 5 (highest score) (465 out of 1296) |
| **Did not respond to this survey Question** | 0.6% (12 out of 2114) |

**Table S118. Mentor helps mentee acquire resources (e.g., grants, fellowships, etc.)**
Analysis by citizenship status. on a Likert Scale (1 lowest-5 highest)

| Theme | count |
|---|---|
| **Non-Citizen (international) Women Mentee (n=442)** | 17.9% rated 1 (lowest score) (79 out of 442)<br>12.4% rated 2 (55 out of 442)<br>18.1% rated 3 (80 out of 442)<br>21.7% rated 4 (96 out of 442)<br>29.9% rated 5 (highest score) (132 out of 442) |
| **Non-Citizen (international) Men Mentee (n=354)** | 18.6% rated 1 (lowest score) (66 out of 354)<br>10.2% rated 2 (36 out of 354)<br>17.2% rated 3 (61 out of 354)<br>24.3% rated 4 (86 out of 354)<br>29.7% rated 5 (highest score) (105 out of 354) |
| **Citizen (national) Women Mentee (n=833)** | 12.6% rated 1 (lowest score) (105 out of 833)<br>12.5% rated 2 (104 out of 833)<br>16.2% rated 3 (135 out of 833)<br>23.9% rated 4 (199 out of 833)<br>34.8% rated 5 (highest score) (290 out of 833) |
| **Citizen (national) Men Mentee (n=445)** | 12.8% rated 1 (lowest score) (57 out of 445)<br>11.2% rated 2 (50 out of 445)<br>15.3% rated 3 (68 out of 445)<br>23.4% rated 4 (104 out of 445)<br>37.3% rated 5 (highest score) (166 out of 445) |

**Table S119. Has your (trainee mentee) key mentor made a website to introduce your group?**
Responses on whether the trainee mentee's key mentor's lab had a website to introduce the lab or not. Responses included "Yes" or "No" only.

| Theme | count |
|---|---|
| **Total (n=2097)** | 71% said Yes (1490 out of 2097)<br>29% said No (607 out of 2097) |
| **Did not respond to this survey Question** | 0.8% (17 out of 2114) |

| **Table S120. Has your (trainee mentee) key mentor posted a welcome letter on the lab/group website for prospective students?** Has your group posted a welcome letter on the lab/group website for prospective students? Example of a welcome letter: https://www.bu.edu/research/ideas-for-a-welcome-to-my-lab-letter/ https://oir.nih.gov/system/files/media/file/2021-08/journal-translational_medicine-bennett-collaboration_science_and_translational_medicine.pdf https://mentoring.ucsf.edu/sites/g/files/tkssra1151/f/wysiwyg/2022%20Welcome%20Letter%20Setting%20Expectations%20final.pdf | |
|---|---|
| **Theme** | **count** |
| **Total (n=2079)** | 13% said Yes (275 out of 2079)<br>87% said No (1804 out of 2079) |
| **Did not respond to this survey Question** | 1.6% (35 out of 2114) |

| **Table S121. Has your (trainee mentee) key mentor written a lab/group manual for current group members?** Have you written a lab/group manual for current group members? | |
|---|---|
| **Theme** | **count** |
| **Total (n=2084)** | 18% said Yes (368 out of 2084)<br>82% said No (1716 out of 2084) |
| **Did not respond to this survey Question** | 1.4% (30 out of 2114) |

| **Table S122. Have you written a lab/group manual for current group members?**<br>**Table from Faculty-to-Faculty mentoring survey study**<br>bioRxiv doi: https://doi.org/10.1101/2022.10.17.512624<br>Have you (faculty mentee) written a lab/group manual for current group members? Data from Faculty mentee survey but presented in this work for the first time. | |
|---|---|
| **Theme** | **count** |
| **Total (n=445)** | 37.5% said Yes (167 out of 445)<br>62.5% said No (278 out of 445) |
| **Women Mentee(n=237)** | 38.8% said Yes (92 out of 237)<br>61.2% said No (145 out of 237) |
| **Men Mentee(n=202)** | 36.6% said Yes (74 out of 202)<br>63.4% said No (128 out of 202) |
| **Non-binary or did not disclose gender (n=6)** | 16.7% said Yes (1 out of 6)<br>83.3% said No (5 out of 6) |
| **Did not respond to this survey Question** | 2.6% (12 out of 457) |

| **Table S123. Does your current lab/group have a website to introduce your group?** Does your (faculty mentee) current lab/group have a website to introduce your group? Data from Faculty mentee survey (bioRxiv doi: https://doi.org/10.1101/2022.10.17.512624) but presented in this work for the first time. | |
|---|---|
| **Theme** | **count** |
| **Total (n=441)** | 72.3% said Yes (319 out of 441)<br>27.7% said No (122 out of 441) |
| **Women Mentee(n=239)** | 72.4% said Yes (173 out of 239)<br>27.6% said No (66 out of 239) |
| **Men Mentee(n=202)** | 72.3% said Yes (146 out of 202)<br>27.7% said No (56 out of 202) |
| **Non-binary or did not disclose gender (n=8)** | 87.5% said Yes (7 out of 8)<br>12.5% said No (1 out of 8) |
| **Did not respond to this survey Question** | 3.5% (16 out of 457) |

| **Table S124. Has your group posted a welcome letter on the lab/group website for prospective students?** Has your (faculty mentee) group posted a welcome letter on the lab/group website for prospective students? Data from Faculty mentee survey (bioRxiv doi: https://doi.org/10.1101/2022.10.17.512624) but presented in this work for the first time. | |
|---|---|
| **Theme** | **count** |
| **Total (n=448)** | 23.2% said Yes (104 out of 448)<br>76.8% said No (344 out of 448) |
| **Women Mentee(n=239)** | 20.5% said Yes (49 out of 239)<br>79.5% said No (190 out of 239) |
| **Men Mentee(n=203)** | 26.6% said Yes (54 out of 203)<br>73.4% said No (149 out of 203) |
| **Non-binary or did not disclose gender (n=6)** | 16.7% said Yes (1 out of 6)<br>83.3% said No (5 out of 6) |
| **Did not respond to this survey Question** | 2% (9 out of 457) |

| **Table S125a part I. corresponding to Figure S2A**<br>**Ordinary one-way ANOVA, Differences in mentees satisfaction with academic career by geographic region (North America, Europe, or other continents).** Data analysis for this table includes survey respondents with "0" number of mentors. | |
|---|---|
| **ANOVA summary** | |
| F | 2.703 |
| P value | 0.0193 |
| P value summary | * |
| Significant diff. among means (P < 0.05)? | Yes |
| R squared | 0.007172 |
| **Brown-Forsythe test** | |
| F (DFn, DFd) | 0.8630 (5, 1871) |

| | | | | | |
|---|---|---|---|---|---|
| P value | 0.5052 | | | | |
| P value summary | ns | | | | |
| Are SDs significantly different (P < 0.05)? | No | | | | |
| **Bartlett's test** | | | | | |
| Bartlett's statistic (corrected) | 3.893 | | | | |
| P value | 0.5649 | | | | |
| P value summary | ns | | | | |
| Are SDs significantly different (P < 0.05)? | No | | | | |
| **ANOVA table** | **SS** | **DF** | **MS** | **F (DFn, DFd)** | **P value** |
| Treatment (between columns) | 20.24 | 5 | 4.048 | F (5, 1871) = 2.703 | P=0.0193 |
| Residual (within columns) | 2802 | 1871 | 1.497 | | |
| Total | 2822 | 1876 | | | |
| **Data summary** | | | | | |
| Number of treatments (columns) | 6 | | | | |
| Number of values (total) | 1877 | | | | |

**Table S125a part II. corresponding to Figure S2A**
**One-way ANOVA results, Multiple Comparisons, Differences in mentees satisfaction with academic career by geographic region (North America, Europe, or other continents).** Data analysis for this table includes survey respondents with "0" number of mentors.

| | | | | | | |
|---|---|---|---|---|---|---|
| **Number of families** | 1 | | | | | |
| **Number of comparisons per family** | 15 | | | | | |
| **Alpha** | 0.05 | | | | | |
| **Tukey's multiple comparisons test** | Mean Diff. | 95.00% CI of diff. | Below threshold? | Summary | Adjusted P Value | |
| Africa vs. Asia | 0.4141 | -0.1767 to 1.005 | No | ns | 0.343 | A-B |
| Africa vs. Europe | 0.3106 | -0.2151 to 0.8364 | No | ns | 0.5416 | A-C |
| Africa vs. North America | 0.1681 | -0.3481 to 0.6843 | No | ns | 0.9391 | A-D |
| Africa vs. South America | 0.2091 | -0.3900 to 0.8081 | No | ns | 0.9195 | A-E |
| Africa vs. Oceania | -0.05775 | -0.6966 to 0.5811 | No | ns | 0.9998 | A-F |
| Asia vs. Europe | -0.1034 | -0.4466 to 0.2398 | No | ns | 0.9559 | B-C |
| Asia vs. North America | -0.246 | -0.5743 to 0.08237 | No | ns | 0.2686 | B-D |

| | | | | | | | | |
|---|---|---|---|---|---|---|---|---|
| Asia vs. South America | -0.205 | -0.6525 to 0.2425 | No | ns | 0.7813 | B-E | | |
| Asia vs. Oceania | -0.4718 | -0.9713 to 0.02762 | No | ns | 0.0767 | B-F | | |
| Europe vs. North America | -0.1426 | -0.3301 to 0.04500 | No | ns | 0.2532 | C-D | | |
| Europe vs. South America | -0.1016 | -0.4588 to 0.2556 | No | ns | 0.9655 | C-E | | |
| Europe vs. Oceania | -0.3684 | -0.7889 to 0.05208 | No | ns | 0.1247 | C-F | | |
| North America vs. South America | 0.04096 | -0.3021 to 0.3840 | No | ns | 0.9994 | D-E | | |
| North America vs. Oceania | -0.2258 | -0.6343 to 0.1826 | No | ns | 0.6138 | D-F | | |
| South America vs. Oceania | -0.2668 | -0.7760 to 0.2424 | No | ns | 0.6678 | E-F | | |
| **Test details** | Mean 1 | Mean 2 | Mean Diff. | SE of diff. | n1 | n2 | q | DF |
| Africa vs. Asia | 3.563 | 3.148 | 0.4141 | 0.2071 | 48 | 128 | 2.827 | 1871 |
| Africa vs. Europe | 3.563 | 3.252 | 0.3106 | 0.1843 | 48 | 540 | 2.384 | 1871 |
| Africa vs. North America | 3.563 | 3.394 | 0.1681 | 0.181 | 48 | 966 | 1.314 | 1871 |
| Africa vs. South America | 3.563 | 3.353 | 0.2091 | 0.21 | 48 | 116 | 1.408 | 1871 |
| Africa vs. Oceania | 3.563 | 3.62 | -0.05775 | 0.2239 | 48 | 79 | 0.3647 | 1871 |
| Asia vs. Europe | 3.148 | 3.252 | -0.1034 | 0.1203 | 128 | 540 | 1.216 | 1871 |
| Asia vs. North America | 3.148 | 3.394 | -0.246 | 0.1151 | 128 | 966 | 3.022 | 1871 |
| Asia vs. South America | 3.148 | 3.353 | -0.205 | 0.1569 | 128 | 116 | 1.848 | 1871 |
| Asia vs. Oceania | 3.148 | 3.62 | -0.4718 | 0.1751 | 128 | 79 | 3.811 | 1871 |
| Europe vs. North America | 3.252 | 3.394 | -0.1426 | 0.06575 | 540 | 966 | 3.066 | 1871 |
| Europe vs. South America | 3.252 | 3.353 | -0.1016 | 0.1252 | 540 | 116 | 1.147 | 1871 |
| Europe vs. Oceania | 3.252 | 3.62 | -0.3684 | 0.1474 | 540 | 79 | 3.535 | 1871 |
| North America vs. South America | 3.394 | 3.353 | 0.04096 | 0.1202 | 966 | 116 | 0.4818 | 1871 |
| North America vs. Oceania | 3.394 | 3.62 | -0.2258 | 0.1432 | 966 | 79 | 2.23 | 1871 |
| South America vs. Oceania | 3.353 | 3.62 | -0.2668 | 0.1785 | 116 | 79 | 2.114 | 1871 |

| **Table S125b part I. corresponding to Figure S2B**<br>**Ordinary one-way ANOVA, Differences in mentees satisfaction with academic career by frequency of meeting with mentor.** Data analysis for this table includes survey respondents with "0" number of mentors. | | | | | |
|---|---|---|---|---|---|
| **ANOVA summary** | | | | | |
| F | 19.14 | | | | |
| P value | <0.0001 | | | | |
| P value summary | **** | | | | |
| Significant diff. among means (P < 0.05)? | Yes | | | | |
| R squared | 0.04856 | | | | |
| **Brown-Forsythe test** | | | | | |
| F (DFn, DFd) | 2.180 (5, 1875) | | | | |
| P value | 0.0538 | | | | |
| P value summary | ns | | | | |
| Are SDs significantly different (P < 0.05)? | No | | | | |
| **Bartlett's test** | | | | | |
| Bartlett's statistic (corrected) | 5.455 | | | | |
| P value | 0.3629 | | | | |
| P value summary | ns | | | | |
| Are SDs significantly different (P < 0.05)? | No | | | | |
| **ANOVA table** | **SS** | **DF** | **MS** | **F (DFn, DFd)** | **P value** |
| Treatment (between columns) | 137.2 | 5 | 27.45 | F (5, 1875) = 19.14 | P<0.0001 |
| Residual (within columns) | 2689 | 1875 | 1.434 | | |
| Total | 2826 | 1880 | | | |
| **Data summary** | | | | | |
| Number of treatments (columns) | 6 | | | | |
| Number of values (total) | 1881 | | | | |

| **Table S125b part II. corresponding to Figure S2B**<br>**One-way ANOVA results**, **Multiple Comparisons, Differences in mentees satisfaction with academic career by frequency of meeting with mentor.** Data analysis for this table includes survey respondents with "0" number of mentors. | | | | | | |
|---|---|---|---|---|---|---|
| **Number of families** | 1 | | | | | |
| **Number of comparisons per family** | 15 | | | | | |
| **Alpha** | 0.05 | | | | | |
| **Tukey's multiple comparisons test** | Mean Diff. | 95.00% CI of diff. | Below threshold? | Summary | Adjusted P Value | |
| As necessary vs. Weekly | -0.04366 | -0.2393 to 0.1519 | No | ns | 0.9882 | A-B |
| As necessary vs. Every 2 weeks | -0.1006 | -0.3414 to 0.1402 | No | ns | 0.8411 | A-C |

| | | | | | | | | |
|---|---|---|---|---|---|---|---|---|
| As necessary vs. Monthly | 0.07741 | -0.2408 to 0.3956 | No | ns | 0.9827 | A-D | | |
| As necessary vs. A few times per year | 0.8158 | 0.5091 to 1.123 | Yes | **** | <0.0001 | A-E | | |
| As necessary vs. Never | 0.8673 | 0.3439 to 1.391 | Yes | **** | <0.0001 | A-F | | |
| Weekly vs. Every 2 weeks | -0.05692 | -0.2931 to 0.1793 | No | ns | 0.9834 | B-C | | |
| Weekly vs. Monthly | 0.1211 | -0.1937 to 0.4358 | No | ns | 0.8826 | B-D | | |
| Weekly vs. A few times per year | 0.8595 | 0.5563 to 1.163 | Yes | **** | <0.0001 | B-E | | |
| Weekly vs. Never | 0.9109 | 0.3897 to 1.432 | Yes | **** | <0.0001 | B-F | | |
| Every 2 weeks vs. Monthly | 0.178 | -0.1667 to 0.5227 | No | ns | 0.6816 | C-D | | |
| Every 2 weeks vs. A few times per year | 0.9164 | 0.5823 to 1.250 | Yes | **** | <0.0001 | C-E | | |
| Every 2 weeks vs. Never | 0.9678 | 0.4280 to 1.508 | Yes | **** | <0.0001 | C-F | | |
| Monthly vs. A few times per year | 0.7384 | 0.3448 to 1.132 | Yes | **** | <0.0001 | D-E | | |
| Monthly vs. Never | 0.7899 | 0.2113 to 1.368 | Yes | ** | 0.0014 | D-F | | |
| A few times per year vs. Never | 0.05146 | -0.5209 to 0.6238 | No | ns | 0.9998 | E-F | | |
| **Test details** | Mean 1 | Mean 2 | Mean Diff. | SE of diff. | n1 | n2 | q | DF |
| As necessary vs. Weekly | 3.411 | 3.454 | -0.04366 | 0.06857 | 577 | 647 | 0.9005 | 1875 |
| As necessary vs. Every 2 weeks | 3.411 | 3.511 | -0.1006 | 0.08442 | 577 | 309 | 1.685 | 1875 |
| As necessary vs. Monthly | 3.411 | 3.333 | 0.07741 | 0.1116 | 577 | 144 | 0.9814 | 1875 |
| As necessary vs. A few times per year | 3.411 | 2.595 | 0.8158 | 0.1075 | 577 | 158 | 10.73 | 1875 |
| As necessary vs. Never | 3.411 | 2.543 | 0.8673 | 0.1835 | 577 | 46 | 6.685 | 1875 |
| Weekly vs. Every 2 weeks | 3.454 | 3.511 | -0.05692 | 0.08281 | 647 | 309 | 0.9721 | 1875 |
| Weekly vs. Monthly | 3.454 | 3.333 | 0.1211 | 0.1103 | 647 | 144 | 1.552 | 1875 |
| Weekly vs. A few times per year | 3.454 | 2.595 | 0.8595 | 0.1063 | 647 | 158 | 11.44 | 1875 |
| Weekly vs. Never | 3.454 | 2.543 | 0.9109 | 0.1827 | 647 | 46 | 7.05 | 1875 |
| Every 2 weeks vs. Monthly | 3.511 | 3.333 | 0.178 | 0.1208 | 309 | 144 | 2.083 | 1875 |
| Every 2 weeks vs. A few times per year | 3.511 | 2.595 | 0.9164 | 0.1171 | 309 | 158 | 11.07 | 1875 |
| Every 2 weeks vs. Never | 3.511 | 2.543 | 0.9678 | 0.1892 | 309 | 46 | 7.233 | 1875 |

| | | | | | | | | |
|---|---|---|---|---|---|---|---|---|
| Monthly vs. A few times per year | 3.333 | 2.595 | 0.7384 | 0.138 | 144 | 158 | 7.569 | 1875 |
| Monthly vs. Never | 3.333 | 2.543 | 0.7899 | 0.2028 | 144 | 46 | 5.508 | 1875 |
| A few times per year vs. Never | 2.595 | 2.543 | 0.05146 | 0.2006 | 158 | 46 | 0.362 7 | 1875 |

**Table S125c part I. corresponding to Figure S2C**
**Ordinary one-way ANOVA, Differences in mentees satisfaction with academic career by quality of mentorship match.** Data analysis for this table includes survey respondents with "0" number of mentors.

| **ANOVA summary** | | | | | |
|---|---|---|---|---|---|
| F | 499.7 | | | | |
| P value | <0.0001 | | | | |
| P value summary | **** | | | | |
| Significant diff. among means (P < 0.05)? | Yes | | | | |
| R squared | 0.5154 | | | | |
| **Brown-Forsythe test** | | | | | |
| F (DFn, DFd) | 26.75 (4, 1879) | | | | |
| P value | <0.0001 | | | | |
| P value summary | **** | | | | |
| Are SDs significantly different (P < 0.05)? | Yes | | | | |
| **Bartlett's test** | | | | | |
| Bartlett's statistic (corrected) | 91.85 | | | | |
| P value | <0.0001 | | | | |
| P value summary | **** | | | | |
| Are SDs significantly different (P < 0.05)? | Yes | | | | |
| **ANOVA table** | **SS** | **DF** | **MS** | **F (DFn, DFd)** | **P value** |
| Treatment (between columns) | 1460 | 4 | 365 | F (4, 1879) = 499.7 | P<0.0001 |
| Residual (within columns) | 1372 | 1879 | 0.7304 | | |
| Total | 2832 | 1883 | | | |
| **Data summary** | | | | | |
| Number of treatments (columns) | 5 | | | | |
| Number of values (total) | 1884 | | | | |

**Table S125c part II. corresponding to Figure S2C**
**One-way ANOVA results, Multiple Comparisons, Differences in mentees satisfaction with academic career by quality of mentorship match.** Data analysis for this table includes survey respondents with "0" number of mentors.

| | | | | | |
|---|---|---|---|---|---|
| **Number of families** | 1 | | | | |
| **Number of comparisons per family** | 10 | | | | |
| **Alpha** | 0.05 | | | | |
| **Tukey's multiple comparisons test** | Mean Diff. | 95.00% CI of diff. | Below threshold ? | Summ ary | Adjusted P Value |

| | | | | | | |
|---|---|---|---|---|---|---|
| Excellent vs. Very good | 0.5501 | 0.3994 to 0.7008 | Yes | **** | <0.0001 | A-B |
| Excellent vs. Good | 1.217 | 1.054 to 1.379 | Yes | **** | <0.0001 | A-C |
| Excellent vs. Fair | 1.904 | 1.726 to 2.083 | Yes | **** | <0.0001 | A-D |
| Excellent vs. Poor | 2.573 | 2.393 to 2.752 | Yes | **** | <0.0001 | A-E |
| Very good vs. Good | 0.6666 | 0.5082 to 0.8249 | Yes | **** | <0.0001 | B-C |
| Very good vs. Fair | 1.354 | 1.179 to 1.529 | Yes | **** | <0.0001 | B-D |
| Very good vs. Poor | 2.023 | 1.846 to 2.199 | Yes | **** | <0.0001 | B-E |
| Good vs. Fair | 0.6877 | 0.5027 to 0.8726 | Yes | **** | <0.0001 | C-D |
| Good vs. Poor | 1.356 | 1.170 to 1.542 | Yes | **** | <0.0001 | C-E |
| Fair vs. Poor | 0.6684 | 0.4679 to 0.8689 | Yes | **** | <0.0001 | D-E |

| **Test details** | Mean 1 | Mean 2 | Mean Diff. | SE of diff. | n1 | n2 | q | DF |
|---|---|---|---|---|---|---|---|---|
| Excellent vs. Very good | 4.382 | 3.832 | 0.5501 | 0.05519 | 455 | 507 | 14.1 | 1879 |
| Excellent vs. Good | 4.382 | 3.166 | 1.217 | 0.05939 | 455 | 380 | 28.97 | 1879 |
| Excellent vs. Fair | 4.382 | 2.478 | 1.904 | 0.06535 | 455 | 274 | 41.21 | 1879 |
| Excellent vs. Poor | 4.382 | 1.81 | 2.573 | 0.06581 | 455 | 268 | 55.29 | 1879 |
| Very good vs. Good | 3.832 | 3.166 | 0.6666 | 0.05799 | 507 | 380 | 16.26 | 1879 |
| Very good vs. Fair | 3.832 | 2.478 | 1.354 | 0.06408 | 507 | 274 | 29.89 | 1879 |
| Very good vs. Poor | 3.832 | 1.81 | 2.023 | 0.06454 | 507 | 268 | 44.32 | 1879 |
| Good vs. Fair | 3.166 | 2.478 | 0.6877 | 0.06773 | 380 | 274 | 14.36 | 1879 |
| Good vs. Poor | 3.166 | 1.81 | 1.356 | 0.06817 | 380 | 268 | 28.13 | 1879 |
| Fair vs. Poor | 2.478 | 1.81 | 0.6684 | 0.07342 | 274 | 268 | 12.87 | 1879 |

**Table S125d part I. corresponding to Figure S2D**
**Ordinary one-way ANOVA, Differences in mentees satisfaction with academic career by gender and citizenship status in country of research.** Data analysis for this table includes survey respondents with "0" number of mentors.

| **ANOVA summary** | |
|---|---|
| F | 4.232 |
| P value | 0.0021 |
| P value summary | ** |
| Significant diff. among means (P < 0.05)? | Yes |
| R squared | 0.009969 |
| **Brown-Forsythe test** | |

| | | | | | |
|---|---|---|---|---|---|
| F (DFn, DFd) | 2.260 (4, 1681) | | | | |
| P value | 0.0606 | | | | |
| P value summary | ns | | | | |
| Are SDs significantly different (P < 0.05)? | No | | | | |
| **Bartlett's test** | | | | | |
| Bartlett's statistic (corrected) | 3.275 | | | | |
| P value | 0.5128 | | | | |
| P value summary | ns | | | | |
| Are SDs significantly different (P < 0.05)? | No | | | | |
| **ANOVA table** | **SS** | **DF** | **MS** | **F (DFn, DFd)** | **P value** |
| Treatment (between columns) | 25.06 | 4 | 6.265 | F (4, 1681) = 4.232 | P=0.0021 |
| Residual (within columns) | 2489 | 1681 | 1.48 | | |
| Total | 2514 | 1685 | | | |
| **Data summary** | | | | | |
| Number of treatments (columns) | 5 | | | | |
| Number of values (total) | 1686 | | | | |

**Table S125d part II. corresponding to Figure S2D**
**One-way ANOVA results**, **Multiple Comparisons, Differences in mentees satisfaction with academic career by gender and citizenship status in country of research.** Data analysis for this table includes survey respondents with "0" number of mentors.

| | | | | | | |
|---|---|---|---|---|---|---|
| **Number of families** | 1 | | | | | |
| **Number of comparisons per family** | 10 | | | | | |
| **Alpha** | 0.05 | | | | | |
| **Tukey's multiple comparisons test** | Mean Diff. | 95.00% CI of diff. | Below threshold ? | Summ ary | Adjusted P Value | |
| W - int vs. W - nat | -0.2386 | -0.4507 to -0.02660 | Yes | * | 0.0183 | A-B |
| W - int vs. M - int | -0.2103 | -0.4779 to 0.05732 | No | ns | 0.2012 | A-C |
| W - int vs. M - nat | -0.3226 | -0.5657 to -0.07956 | Yes | ** | 0.0028 | A-D |
| W - int vs. NB | 0.148 | -0.5249 to 0.8208 | No | ns | 0.975 | A-E |
| W - nat vs. M - int | 0.02834 | -0.2171 to 0.2738 | No | ns | 0.9979 | B-C |
| W - nat vs. M - nat | -0.08398 | -0.3024 to 0.1345 | No | ns | 0.832 | B-D |
| W - nat vs. NB | 0.3866 | -0.2778 to 1.051 | No | ns | 0.5046 | B-E |
| M - int vs. M - nat | -0.1123 | -0.3851 to 0.1604 | No | ns | 0.7936 | C-D |
| M - int vs. NB | 0.3583 | -0.3259 to 1.042 | No | ns | 0.6083 | C-E |

| | | | | | | |
|---|---|---|---|---|---|---|
| M - nat vs. NB | 0.4706 | -0.2043 to 1.146 | No | ns | 0.3155 | D-E |

| Test details | Mean 1 | Mean 2 | Mean Diff. | SE of diff. | n1 | n2 | q | DF |
|---|---|---|---|---|---|---|---|---|
| W - int vs. W - nat | 3.148 | 3.387 | -0.2386 | 0.07765 | 392 | 657 | 4.346 | 1681 |
| W - int vs. M - int | 3.148 | 3.358 | -0.2103 | 0.098 | 392 | 254 | 3.035 | 1681 |
| W - int vs. M - nat | 3.148 | 3.471 | -0.3226 | 0.08901 | 392 | 357 | 5.126 | 1681 |
| W - int vs. NB | 3.148 | 3 | 0.148 | 0.2464 | 392 | 26 | 0.8492 | 1681 |
| W - nat vs. M - int | 3.387 | 3.358 | 0.02834 | 0.0899 | 657 | 254 | 0.4458 | 1681 |
| W - nat vs. M - nat | 3.387 | 3.471 | -0.08398 | 0.08 | 657 | 357 | 1.485 | 1681 |
| W - nat vs. NB | 3.387 | 3 | 0.3866 | 0.2433 | 657 | 26 | 2.247 | 1681 |
| M - int vs. M - nat | 3.358 | 3.471 | -0.1123 | 0.09988 | 254 | 357 | 1.59 | 1681 |
| M - int vs. NB | 3.358 | 3 | 0.3583 | 0.2505 | 254 | 26 | 2.022 | 1681 |
| M - nat vs. NB | 3.471 | 3 | 0.4706 | 0.2472 | 357 | 26 | 2.693 | 1681 |

**Table S125e part I. corresponding to Figure S2E**
**Ordinary one-way ANOVA, Differences in mentees satisfaction with academic career by gender.** Data analysis for this table includes survey respondents with "0" number of mentors.

| ANOVA summary | | | | | |
|---|---|---|---|---|---|
| F | 3.093 | | | | |
| P value | 0.0456 | | | | |
| P value summary | * | | | | |
| Significant diff. among means (P < 0.05)? | Yes | | | | |
| R squared | 0.003662 | | | | |
| **Brown-Forsythe test** | | | | | |
| F (DFn, DFd) | 1.932 (2, 1683) | | | | |
| P value | 0.1452 | | | | |
| P value summary | ns | | | | |
| Are SDs significantly different (P < 0.05)? | No | | | | |
| **Bartlett's test** | | | | | |
| Bartlett's statistic (corrected) | 0.9021 | | | | |
| P value | 0.6369 | | | | |
| P value summary | ns | | | | |
| Are SDs significantly different (P < 0.05)? | No | | | | |
| **ANOVA table** | **SS** | **DF** | **MS** | **F (DFn, DFd)** | **P value** |
| Treatment (between columns) | 9.204 | 2 | 4.602 | F (2, 1683) = 3.093 | P=0.0456 |
| Residual (within columns) | 2504 | 1683 | 1.488 | | |
| Total | 2514 | 1685 | | | |

| **Table S125e part II. corresponding to Figure S2E<br>One-way ANOVA results, Multiple Comparisons, Differences in mentees satisfaction with academic career by gender.** Data analysis for this table includes survey respondents with "0" number of mentors. | | | | | | | | |
|---|---|---|---|---|---|---|---|---|
| **Number of families** | 1 | | | | | | | |
| **Number of comparisons per family** | 3 | | | | | | | |
| **Alpha** | 0.05 | | | | | | | |
| **Tukey's multiple comparisons test** | Mean Diff. | 95.00% CI of diff. | Below threshold ? | Summ ary | Adjusted P Value | | | |
| W vs. M | -0.1265 | -0.2721 to 0.01916 | No | ns | 0.1037 | A-B | | |
| W vs. NB | 0.2974 | -0.2707 to 0.8655 | No | ns | 0.4367 | A-C | | |
| M vs. NB | 0.4239 | -0.1491 to 0.9969 | No | ns | 0.1923 | B-C | | |
| **Test details** | Mean 1 | Mean 2 | Mean Diff. | SE of diff. | n1 | n2 | q | DF |
| W vs. M | 3.297 | 3.424 | -0.1265 | 0.06208 | 1049 | 611 | 2.881 | 1683 |
| W vs. NB | 3.297 | 3 | 0.2974 | 0.2422 | 1049 | 26 | 1.737 | 1683 |
| M vs. NB | 3.424 | 3 | 0.4239 | 0.2443 | 611 | 26 | 2.454 | 1683 |

| **Table S125f. corresponding to Figure S2F<br>Ordinary one-way ANOVA, Differences in mentees satisfaction with academic career by citizenship status in country of research.** Data analysis for this table includes survey respondents with "0" number of mentors. | |
|---|---|
| **ANOVA summary** | |
| Column B | Nat |
| vs. | vs. |
| Column A | Int |
| **Unpaired t-test** | |
| P value | 0.0025 |
| P value summary | ** |
| Significantly different (P < 0.05)? | Yes |
| One- or two-tailed P value? | Two-tailed |
| t, df | t=3.032, df=1658 |
| **How big is the difference?** | |
| Mean of column A | 3.231 |
| Mean of column B | 3.416 |
| Difference between means (B - A) ± SEM | 0.1855 ± 0.06118 |
| 95% confidence interval | 0.06552 to 0.3055 |
| R squared (eta squared) | 0.005515 |
| **F test to compare variance** | |
| F, DFn, Dfd | 1.111, 645, 1013 |
| P value | 0.137 |

| | |
|---|---|
| P value summary | ns |
| Significantly different (P < 0.05)? | No |
| **Data analyzed** | |
| Sample size, column A | 646 |
| Sample size, column A | 646 |

**Table S126a. corresponding to Figure S3A**
**One-way ANOVA results, Multiple Comparisons, Differences in quality of match with mentor (on a poor-to-excellent Likert scale) by gender and citizenship status in country of research.**
Data analysis for this table includes survey respondents with "0" number of mentors.

| | |
|---|---|
| **Number of families** | 1 |
| **Number of comparisons per family** | 6 |
| **Alpha** | 0.05 |

| **Tukey's multiple comparisons test** | Mean Diff. | 95.00% CI of diff. | Below threshold ? | Summary | Adjusted P Value | |
|---|---|---|---|---|---|---|
| W - int vs. W - nat | -0.319 | -0.5242 to -0.1137 | Yes | *** | 0.0004 | A-B |
| W - int vs. M - int | -0.1029 | -0.3521 to 0.1463 | No | ns | 0.7132 | A-C |
| W - int vs. M - nat | -0.3502 | -0.5846 to -0.1158 | Yes | *** | 0.0007 | A-D |
| W - int vs. NB | 0.2161 | -0.005582 to 0.4378 | No | ns | 0.0592 | B-C |
| W - nat vs. M - int | -0.03122 | -0.2362 to 0.1737 | No | ns | 0.9796 | B-D |
| W - nat vs. M - nat | -0.2473 | -0.4963 to 0.001646 | No | ns | 0.0523 | C-D |

| **Test details** | Mean 1 | Mean 2 | Mean Diff. | SE of diff. | n1 | n2 | q | DF |
|---|---|---|---|---|---|---|---|---|
| W - int vs. W - nat | 3.081 | 3.4 | -0.319 | 0.07982 | 443 | 837 | 5.651 | 2074 |
| W - int vs. M - int | 3.081 | 3.184 | -0.1029 | 0.09693 | 443 | 353 | 1.501 | 2074 |
| W - int vs. M - nat | 3.081 | 3.431 | -0.3502 | 0.09118 | 443 | 445 | 5.431 | 2074 |
| W - int vs. NB | 3.4 | 3.184 | 0.2161 | 0.08622 | 837 | 353 | 3.545 | 2074 |
| W - nat vs. M - int | 3.4 | 3.431 | -0.03122 | 0.07971 | 837 | 445 | 0.554 | 2074 |
| W - nat vs. M - nat | 3.184 | 3.431 | -0.2473 | 0.09683 | 353 | 445 | 3.612 | 2074 |

| Table S126b. corresponding to Figure S3B<br>**Two-tailed Mann-Whitney test results, Women mentees (Column A) versus Men mentees (Column B) on the quality of their match with mentor.** Data analysis for this table excludes survey respondents with "0" number of mentors. | |
|---|---|
| **Mann-Whitney test** | |
| P value | 0.6294 |
| Exact or approximate P value? | Approximate |
| P value summary | ns |
| Significantly different (P < 0.05)? | No |
| One- or two-tailed P value? | Two-tailed |
| Sum of ranks in column A, B | 1324293 , 835788 |
| Mann-Whitney U | 0.6294 |
| Difference between medians | |
| Median of column A | 4.000, n=1280 |
| Median of column B | 4.000, n=798 |
| Difference: Actual | 0 |
| Difference: Hodges-Lehmann | 0 |

| Table S126c. corresponding to Figure S3C<br>**Two-tailed Mann-Whitney test results, International mentees (Column A) versus National mentees (Column B) on the quality of their match with mentor.** Data analysis for this table excludes survey respondents with "0" number of mentors. | |
|---|---|
| **Mann-Whitney test** | |
| P value | <0.0001 |
| Exact or approximate P value? | Approximate |
| P value summary | **** |
| Significantly different (P < 0.05)? | Yes |
| One- or two-tailed P value? | Two-tailed |
| Sum of ranks in column A, B | 767776 , 1392305 |
| Mann-Whitney U | 450570 |
| Difference between medians | |
| Median of column A | 3.000, n=796 |
| Median of column B | 4.000, n=1282 |
| Difference: Actual | 1 |
| Difference: Hodges-Lehmann | 0 |

| Table S127a. corresponding to Figure S5A<br>**Two-tailed Mann-Whitney test results, National women mentees (Column B) versus International women mentees (Column A) on the quality of their match with mentor.** Data analysis for this table excludes survey respondents with "0" number of mentors. | |
|---|---|
| **Mann-Whitney test** | |
| P value | <0.0001 |
| Exact or approximate P value? | Approximate |
| P value summary | **** |
| Significantly different (P < 0.05)? | Yes |
| One- or two-tailed P value? | Two-tailed |
| Sum of ranks in column A, B | 259703 , 561419 |
| Mann-Whitney U | 161357 |
| Difference between medians | |
| Median of column A | 3.000, n=443 |
| Median of column B | 4.000, n=838 |
| Difference: Actual | 1 |
| Difference: Hodges-Lehmann | 0 |

| **Table S127b. corresponding to Figure S5B**<br>**Two-tailed Mann-Whitney test results, National women mentees (Column B) versus International women mentees (Column A) on constructiveness of their interactions with mentor.** Data analysis for this table excludes survey respondents with "0" number of mentors. | |
|---|---|
| **Mann-Whitney test** | |
| P value | 0.0093 |
| Exact or approximate P value? | Approximate |
| P value summary | ** |
| Significantly different (P < 0.05)? | Yes |
| One- or two-tailed P value? | Two-tailed |
| Sum of ranks in column A, B | 270162 , 545841 |
| Mann-Whitney U | 171816 |
| Difference between medians | |
| Median of column A | 1.000, n=443 |
| Median of column B | 1.000, n=834 |
| Difference: Actual | 0 |
| Difference: Hodges-Lehmann | 0 |

| **Table S127c. corresponding to Figure S5C**<br>**Two-tailed Mann-Whitney test results, National women mentees (Column B) versus International women mentees (Column A) on their academic satisfaction.** Data analysis for this table excludes survey respondents with "0" number of mentors. | |
|---|---|
| **Mann-Whitney test** | |
| P value | 0.0033 |
| Exact or approximate P value? | Approximate |
| P value summary | ** |
| Significantly different (P < 0.05)? | Yes |
| One- or two-tailed P value? | Two-tailed |
| Sum of ranks in column A, B | 212248 , 461132 |
| Mann-Whitney U | 135220 |
| Difference between medians | |
| Median of column A | 3.000, n=392 |
| Median of column B | 4.000, n=768 |
| Difference: Actual | 1 |
| Difference: Hodges-Lehmann | 0 |

| **Table S127d. corresponding to Figure S5D**<br>**Two-tailed Mann-Whitney test results, National women mentees (Column B) versus International women mentees (Column A) on their career optimism.** Data analysis for this table excludes survey respondents with "0" number of mentors. | |
|---|---|
| **Mann-Whitney test** | |
| P value | |
| Exact or approximate P value? | 0.0132 |
| P value summary | Approximate |
| Significantly different (P < 0.05)? | * |
| One- or two-tailed P value? | Yes |
| Sum of ranks in column A, B | Two-tailed |
| Mann-Whitney U | 214077 , 459303 |
| Difference between medians | |
| Median of column A | 3.000, n=391 |
| Median of column B | 4.000, n=769 |
| Difference: Actual | 1 |
| Difference: Hodges-Lehmann | 0 |

| Table S127e. corresponding to Figure S5E<br>**Two-tailed Mann-Whitney test results, National men mentees (Column B) versus International men mentees (Column A) on the quality of their match with mentor.** Data analysis for this table excludes survey respondents with "0" number of mentors. | |
|---|---|
| **Mann-Whitney test** | |
| P value | 0.0112 |
| Exact or approximate P value? | Approximate |
| P value summary | * |
| Significantly different (P < 0.05)? | Yes |
| One- or two-tailed P value? | Two-tailed |
| Sum of ranks in column A, B | 133027 , 185775 |
| Mann-Whitney U | 70546 |
| Difference between medians | |
| Median of column A | 3.000, n=353 |
| Median of column B | 4.000, n=445 |
| Difference: Actual | 1 |
| Difference: Hodges-Lehmann | 0 |

| Table S127f. corresponding to Figure S5F<br>**Two-tailed Mann-Whitney test results, National men mentees (Column B) versus International men mentees (Column A) on constructiveness of their interactions with mentor.** Data analysis for this table excludes survey respondents with "0" number of mentors. | |
|---|---|
| **Mann-Whitney test** | |
| P value | 0.0027 |
| Exact or approximate P value? | Approximate |
| P value summary | ** |
| Significantly different (P < 0.05)? | Yes |
| One- or two-tailed P value? | Two-tailed |
| Sum of ranks in column A, B | 132942 , 185061 |
| Mann-Whitney U | 70814 |
| Difference between medians | |
| Median of column A | 1.000, n=352 |
| Median of column B | 1.000, n=445 |
| Difference: Actual | 0 |
| Difference: Hodges-Lehmann | 0 |

| Table S127g. corresponding to Figure S5G<br>**Two-tailed Mann-Whitney test results, National men mentees (Column B) versus International men mentees (Column A) on their academic satisfaction.** Data analysis for this table excludes survey respondents with "0" number of mentors. | |
|---|---|
| **Mann-Whitney test** | |
| P value | 0.2205 |
| Exact or approximate P value? | Approximate |
| P value summary | ns |
| Significantly different (P < 0.05)? | No |
| One- or two-tailed P value? | Two-tailed |
| Sum of ranks in column A, B | 110548 , 151902 |
| Mann-Whitney U | 61093 |
| Difference between medians | |
| Median of column A | 4.000, n=314 |
| Median of column B | 4.000, n=410 |
| Difference: Actual | 0 |
| Difference: Hodges-Lehmann | 0 |

**Table S127h. corresponding to Figure S5H**
**Two-tailed Mann-Whitney test results, National men mentees (Column B) versus International men mentees (Column A) on their career optimism.** Data analysis for this table excludes survey respondents with "0" number of mentors.

| **Mann-Whitney test** | |
|---|---|
| P value | |
| Exact or approximate P value? | 0.1175 |
| P value summary | Approximate |
| Significantly different (P < 0.05)? | ns |
| One- or two-tailed P value? | No |
| Sum of ranks in column A, B | Two-tailed |
| Mann-Whitney U | 110507 , 152668 |
| Difference between medians | |
| Median of column A | 4.000, n=316 |
| Median of column B | 4.000, n=409 |
| Difference: Actual | 0 |
| Difference: Hodges-Lehmann | 0 |

**Table S128a. corresponding to Figure S4A**
**Two-tailed Mann-Whitney test results, Women (Column B) versus Men mentees (Column A) on their career optimism.** Data analysis for this table excludes survey respondents with "0" number of mentors.

| **Mann-Whitney test** | |
|---|---|
| P value | 0.0019 |
| Exact or approximate P value? | Approximate |
| P value summary | ** |
| Significantly different (P < 0.05)? | Yes |
| One- or two-tailed P value? | Two-tailed |
| Sum of ranks in column A, B | 718019 , 1059537 |
| Mann-Whitney U | 386157 |
| Difference between medians | |
| Median of column A | 4.000, n=725 |
| Median of column B | 4.000, n=1160 |
| Difference: Actual | 0 |
| Difference: Hodges-Lehmann | 0 |

**Table S128b. corresponding to Figure S4B**
**Two-tailed Mann-Whitney test results, National (Column B) versus International mentees (Column A) on quality of their match with mentor.** Data analysis for this table excludes survey respondents with "0" number of mentors.

| **Mann-Whitney test** | |
|---|---|
| P value | <0.0001 |
| Exact or approximate P value? | Approximate |
| P value summary | **** |
| Significantly different (P < 0.05)? | Yes |
| One- or two-tailed P value? | Two-tailed |
| Sum of ranks in column A, B | 767776 , 1392305 |
| Mann-Whitney U | 450570 |
| Difference between medians | |
| Median of column A | 3.000, n=796 |
| Median of column B | 4.000, n=1282 |
| Difference: Actual | 1 |
| Difference: Hodges-Lehmann | 0 |

| **Table S128c. corresponding to Figure S4C**<br>**Two-tailed Mann-Whitney test results, National (Column B) versus International mentees (Column A) on constructiveness of their interactions with mentor.** Data analysis for this table excludes survey respondents with "0" number of mentors. | |
|---|---|
| **Mann-Whitney test** | |
| P value | 0.0001 |
| Exact or approximate P value? | Approximate |
| P value summary | *** |
| Significantly different (P < 0.05)? | Yes |
| One- or two-tailed P value? | Two-tailed |
| Sum of ranks in column A, B | 785228 , 1366548 |
| Mann-Whitney U | 468818 |
| Difference between medians | |
| Median of column A | 1.000, n=795 |
| Median of column B | 1.000, n=1279 |
| Difference: Actual | 0 |
| Difference: Hodges-Lehmann | 0 |

| **Table S128d. corresponding to Figure S4D**<br>**Two-tailed Mann-Whitney test results, National (Column B) versus International mentees (Column A) on their academic satisfaction.** Data analysis for this table excludes survey respondents with "0" number of mentors. | |
|---|---|
| **Mann-Whitney test** | |
| P value | 0.0001 |
| Exact or approximate P value? | Approximate |
| P value summary | *** |
| Significantly different (P < 0.05)? | Yes |
| One- or two-tailed P value? | Two-tailed |
| Sum of ranks in column A, B | 1184214 , 1395642 |
| Mann-Whitney U | 585249 |
| Difference between medians | |
| Median of column A | 3.000, n=1094 |
| Median of column B | 4.000, n=1177 |
| Difference: Actual | 1 |
| Difference: Hodges-Lehmann | 0 |

| **Table S128e corresponding to Figure S4E**<br>**Two-tailed Mann-Whitney test results, National (Column B) versus International mentees (Column A) on their career optimism.** Data analysis for this table excludes survey respondents with "0" number of mentors. | |
|---|---|
| **Mann-Whitney test** | |
| P value | 0.0094 |
| Exact or approximate P value? | Approximate |
| P value summary | ** |
| Significantly different (P < 0.05)? | Yes |
| One- or two-tailed P value? | Two-tailed |
| Sum of ranks in column A, B | 637733 , 1137938 |
| Mann-Whitney U | 387455 |
| Difference between medians | |
| Median of column A | 4.000, n=707 |
| Median of column B | 4.000, n=1177 |
| Difference: Actual | 0 |
| Difference: Hodges-Lehmann | 0 |

| **Table S129a. corresponding to Figure S6A**<br>**One-way ANOVA results**, **Multiple Comparisons, whether mentor builds confidence in mentee (on a 1- to 5 Likert scale) by gender and citizenship status in country of research.** Data analysis for this table includes survey respondents with "0" number of mentors. | | | | | | | | |
|---|---|---|---|---|---|---|---|---|
| **Number of families** | 1 | | | | | | | |
| **Number of comparisons per family** | 6 | | | | | | | |
| **Alpha** | 0.05 | | | | | | | |
| **Tukey's multiple comparisons test** | Mean Diff. | 95.00% CI of diff. | Below threshold ? | Summ ary | Adjusted P Value | | | |
| W - int vs. W - nat | -0.2032 | -0.4318 to 0.02546 | No | ns | 0.1019 | A-B | | |
| W - int vs. M - int | - 0.0448 4 | -0.3222 to 0.2325 | No | ns | 0.9758 | A-C | | |
| W - int vs. M - nat | -0.2779 | -0.5386 to - 0.01723 | Yes | * | 0.0314 | A-D | | |
| W - int vs. NB | 0.1583 | -0.08815 to 0.4048 | No | ns | 0.3499 | B-C | | |
| W - nat vs. M - int | - 0.0747 6 | -0.3024 to 0.1528 | No | ns | 0.8331 | B-D | | |
| W - nat vs. M - nat | -0.2331 | -0.5096 to 0.04340 | No | ns | 0.1327 | C-D | | |
| **Test details** | Mean 1 | Mean 2 | Mean Diff. | SE of diff. | n1 | n2 | q | DF |
| W - int vs. W - nat | 3.1 | 3.304 | -0.2032 | 0.0889 2 | 438 | 830 | 3.231 | 2059 |
| W - int vs. M - int | 3.1 | 3.145 | -0.04484 | 0.1079 | 438 | 351 | 0.588 | 2059 |
| W - int vs. M - nat | 3.1 | 3.378 | -0.2779 | 0.1014 | 438 | 444 | 3.876 | 2059 |
| W - int vs. NB | 3.304 | 3.145 | 0.1583 | 0.0958 6 | 830 | 351 | 2.336 | 2059 |
| W - nat vs. M - int | 3.304 | 3.378 | -0.07476 | 0.0885 2 | 830 | 444 | 1.194 | 2059 |
| W - nat vs. M - nat | 3.145 | 3.378 | -0.2331 | 0.1075 | 351 | 444 | 3.065 | 2059 |

| **Table S129b. corresponding to Figure S6B**<br>**One-way ANOVA results**, **Multiple Comparisons, whether mentor coordinates their mentorship with mentee 's other mentors (on a 1- to 5 Likert scale) by gender and citizenship status in country of research.** Data analysis for this table includes survey respondents with "0" number of mentors. | | | | | |
|---|---|---|---|---|---|
| **Number of families** | 1 | | | | |
| **Number of comparisons per family** | 6 | | | | |
| **Alpha** | 0.05 | | | | |
| **Tukey's multiple comparisons test** | Mean Diff. | 95.00% CI of diff. | Below threshold ? | Summ ary | Adjusted P Value |

| | | | | | | |
|---|---|---|---|---|---|---|
| W - int vs. W - nat | -0.1898 | -0.3954 to 0.01576 | No | ns | 0.0824 | A-B |
| W - int vs. M - int | -0.05116 | -0.3013 to 0.1989 | No | ns | 0.9528 | A-C |
| W - int vs. M - nat | -0.2553 | -0.4897 to -0.02089 | Yes | * | 0.0265 | A-D |
| W - int vs. NB | 0.1387 | -0.08379 to 0.3611 | No | ns | 0.3773 | B-C |
| W - nat vs. M - int | -0.06548 | -0.2701 to 0.1392 | No | ns | 0.8438 | B-D |
| W - nat vs. M - nat | -0.2041 | -0.4535 to 0.04520 | No | ns | 0.1518 | C-D |

| **Test details** | Mean 1 | Mean 2 | Mean Diff. | SE of diff. | n1 | n2 | q | DF |
|---|---|---|---|---|---|---|---|---|
| W - int vs. W - nat | 2.949 | 3.139 | -0.1898 | 0.07995 | 430 | 815 | 3.357 | 2018 |
| W - int vs. M - int | 2.949 | 3 | -0.05116 | 0.09727 | 430 | 341 | 0.7439 | 2018 |
| W - int vs. M - nat | 2.949 | 3.204 | -0.2553 | 0.09117 | 430 | 436 | 3.96 | 2018 |
| W - int vs. NB | 3.139 | 3 | 0.1387 | 0.08651 | 815 | 341 | 2.267 | 2018 |
| W - nat vs. M - int | 3.139 | 3.204 | -0.06548 | 0.07959 | 815 | 436 | 1.163 | 2018 |
| W - nat vs. M - nat | 3 | 3.204 | -0.2041 | 0.09697 | 341 | 436 | 2.977 | 2018 |

**Table S129c. corresponding to Figure S6C**
**One-way ANOVA results, Multiple Comparisons, whether mentor helps mentee to strategize their career goals (on a 1- to 5 Likert scale) by gender and citizenship status in country of research.** Data analysis for this table includes survey respondents with "0" number of mentors.

| | |
|---|---|
| **Number of families** | 1 |
| **Number of comparisons per family** | 6 |
| **Alpha** | 0.05 |

| **Tukey's multiple comparisons test** | Mean Diff. | 95.00% CI of diff. | Below threshold ? | Summary | Adjusted P Value | |
|---|---|---|---|---|---|---|
| W - int vs. W - nat | -0.2428 | -0.4612 to -0.02447 | Yes | * | 0.0223 | A-B |
| W - int vs. M - int | -0.2986 | -0.5632 to -0.03410 | Yes | * | 0.0196 | A-C |
| W - int vs. M - nat | -0.3894 | -0.6390 to -0.1397 | Yes | *** | 0.0004 | A-D |
| W - int vs. NB | -0.05578 | -0.2907 to 0.1792 | No | ns | 0.9288 | B-C |
| W - nat vs. M - int | -0.1465 | -0.3646 to 0.07154 | No | ns | 0.3095 | B-D |

| | | | | | | |
|---|---|---|---|---|---|---|
| W - nat vs. M - nat | -0.09074 | -0.3550 to 0.1735 | No | ns | 0.8138 | C-D |

| Test details | Mean 1 | Mean 2 | Mean Diff. | SE of diff. | n1 | n2 | q | DF |
|---|---|---|---|---|---|---|---|---|
| W - int vs. W - nat | 2.84 | 3.083 | -0.2428 | 0.08493 | 438 | 831 | 4.044 | 2058 |
| W - int vs. M - int | 2.84 | 3.139 | -0.2986 | 0.1029 | 438 | 353 | 4.105 | 2058 |
| W - int vs. M - nat | 2.84 | 3.23 | -0.3894 | 0.09709 | 438 | 440 | 5.672 | 2058 |
| W - int vs. NB | 3.083 | 3.139 | -0.05578 | 0.09138 | 831 | 353 | 0.8632 | 2058 |
| W - nat vs. M - int | 3.083 | 3.23 | -0.1465 | 0.08481 | 831 | 440 | 2.443 | 2058 |
| W - nat vs. M - nat | 3.139 | 3.23 | -0.09074 | 0.1028 | 353 | 440 | 1.248 | 2058 |

**Table S129d. corresponding to Figure S6D**
**One-way ANOVA results, Multiple Comparisons, whether mentor helps mentee to strategize their career goals (on a 1- to 5 Likert scale) by gender and citizenship status in country of research.** Data analysis for this table includes survey respondents with "0" number of mentors.

| | |
|---|---|
| **Number of families** | 1 |
| **Number of comparisons per family** | 6 |
| **Alpha** | 0.05 |

| **Tukey's multiple comparisons test** | Mean Diff. | 95.00% CI of diff. | Below threshold? | Summary | Adjusted P Value | |
|---|---|---|---|---|---|---|
| W - int vs. W - nat | -0.2288 | -0.4444 to -0.01311 | Yes | * | 0.0326 | A-B |
| W - int vs. M - int | -0.2436 | -0.5045 to 0.01735 | No | ns | 0.0773 | A-C |
| W - int vs. M - nat | -0.3494 | -0.5954 to -0.1035 | Yes | ** | 0.0015 | A-D |
| W - int vs. NB | -0.01481 | -0.2465 to 0.2169 | No | ns | 0.9984 | B-C |
| W - nat vs. M - int | -0.1207 | -0.3354 to 0.09400 | No | ns | 0.4711 | B-D |
| W - nat vs. M - nat | -0.1059 | -0.3660 to 0.1542 | No | ns | 0.722 | C-D |

| **Test details** | Mean 1 | Mean 2 | Mean Diff. | SE of diff. | n1 | n2 | q | DF |
|---|---|---|---|---|---|---|---|---|
| W - int vs. W - nat | 2.816 | 3.045 | -0.2288 | 0.08387 | 435 | 825 | 3.857 | 2049 |
| W - int vs. M - int | 2.816 | 3.06 | -0.2436 | 0.1015 | 435 | 352 | 3.394 | 2049 |
| W - int vs. M - nat | 2.816 | 3.166 | -0.3494 | 0.09565 | 435 | 441 | 5.167 | 2049 |

| | | | | | | | | |
|---|---|---|---|---|---|---|---|---|
| W - int vs. NB | 3.045 | 3.06 | -0.01481 | 0.09011 | 825 | 352 | 0.2324 | 2049 |
| W - nat vs. M - int | 3.045 | 3.166 | -0.1207 | 0.0835 | 825 | 441 | 2.044 | 2049 |
| W - nat vs. M - nat | 3.06 | 3.166 | -0.1059 | 0.1012 | 352 | 441 | 1.48 | 2049 |

**Table S129e. corresponding to Figure S14e**
**One-way ANOVA results**, **Multiple Comparisons, whether mentor is aware of their biases (on a 1- to 5 Likert scale) by gender and citizenship status in country of research.** Data analysis for this table includes survey respondents with "0" number of mentors.

| | |
|---|---|
| **Number of families** | 1 |
| **Number of comparisons per family** | 6 |
| **Alpha** | 0.05 |

| **Tukey's multiple comparisons test** | Mean Diff. | 95.00% CI of diff. | Below threshold? | Summary | Adjusted P Value | |
|---|---|---|---|---|---|---|
| W - int vs. W - nat | -0.003985 | -0.2036 to 0.1956 | No | ns | >0.9999 | A-B |
| W - int vs. M - int | -0.09685 | -0.3386 to 0.1449 | No | ns | 0.7317 | A-C |
| W - int vs. M - nat | -0.2826 | -0.5105 to -0.05481 | Yes | ** | 0.0079 | A-D |
| W - int vs. NB | -0.09287 | -0.3080 to 0.1223 | No | ns | 0.6836 | B-C |
| W - nat vs. M - int | -0.2787 | -0.4781 to -0.07921 | Yes | ** | 0.0019 | B-D |
| W - nat vs. M - nat | -0.1858 | -0.4274 to 0.05580 | No | ns | 0.1969 | C-D |

| **Test details** | Mean 1 | Mean 2 | Mean Diff. | SE of diff. | n1 | n2 | q | DF |
|---|---|---|---|---|---|---|---|---|
| W - int vs. W - nat | 2.787 | 2.791 | -0.003985 | 0.07763 | 442 | 829 | 0.07259 | 2064 |
| W - int vs. M - int | 2.787 | 2.884 | -0.09685 | 0.09401 | 442 | 354 | 1.457 | 2064 |
| W - int vs. M - nat | 2.787 | 3.07 | -0.2826 | 0.08862 | 442 | 443 | 4.511 | 2064 |
| W - int vs. NB | 2.791 | 2.884 | -0.09287 | 0.08369 | 829 | 354 | 1.569 | 2064 |
| W - nat vs. M - int | 2.791 | 3.07 | -0.2787 | 0.07757 | 829 | 443 | 5.08 | 2064 |
| W - nat vs. M - nat | 2.884 | 3.07 | -0.1858 | 0.09397 | 354 | 443 | 2.796 | 2064 |

**Table S129f. corresponding to Figure S6F**
**One-way ANOVA results, Multiple Comparisons, whether mentor is aware of their biases (on a 1- to 5 Likert scale) by gender and citizenship status in country of research.** Data analysis for this table includes survey respondents with "0" number of mentors.

| | |
|---|---|
| **Number of families** | 1 |
| **Number of comparisons per family** | 6 |
| **Alpha** | 0.05 |

| **Tukey's multiple comparisons test** | Mean Diff. | 95.00% CI of diff. | Below threshold? | Summary | Adjusted P Value | |
|---|---|---|---|---|---|---|
| W - int vs. W - nat | -0.1355 | -0.3475 to 0.07657 | No | ns | 0.355 | A-B |
| W - int vs. M - int | -0.1242 | -0.3811 to 0.1328 | No | ns | 0.5998 | A-C |
| W - int vs. M - nat | -0.1355 | -0.3475 to 0.07657 | No | ns | 0.355 | A-D |
| W - int vs. NB | 0.01128 | -0.2169 to 0.2394 | No | ns | 0.9993 | B-C |
| W - nat vs. M - int | 0 | -0.1760 to 0.1760 | No | ns | >0.9999 | B-D |
| W - nat vs. M - nat | -0.01128 | -0.2394 to 0.2169 | No | ns | 0.9993 | C-D |

| **Test details** | Mean 1 | Mean 2 | Mean Diff. | SE of diff. | n1 | n2 | q | DF |
|---|---|---|---|---|---|---|---|---|
| W - int vs. W - nat | 2.833 | 2.969 | -0.1355 | 0.08247 | 438 | 833 | 2.323 | 2453 |
| W - int vs. M - int | 2.833 | 2.958 | -0.1242 | 0.09995 | 438 | 353 | 1.757 | 2453 |
| W - int vs. M - nat | 2.833 | 2.969 | -0.1355 | 0.08247 | 438 | 833 | 2.323 | 2453 |
| W - int vs. NB | 2.969 | 2.958 | 0.01128 | 0.08874 | 833 | 353 | 0.1798 | 2453 |
| W - nat vs. M - int | 2.969 | 2.969 | 0 | 0.06847 | 833 | 833 | 0 | 2453 |
| W - nat vs. M - nat | 2.958 | 2.969 | -0.01128 | 0.08874 | 353 | 833 | 0.1798 | 2453 |
